# Ts66Yah, an upgraded Ts65Dn mouse model for Down syndrome, for only the region homologous to Human chromosome 21

**DOI:** 10.1101/2022.06.06.494940

**Authors:** Arnaud Duchon, Maria del Mar Muñiz Moreno, Claire Chevalier, Valérie Nalesso, Philippe Andre, Marta Fructuoso-Castellar, Mary Mondino, Chrystelle Po, Vincent Noblet, Marie-Christine Birling, Marie-Claude Potier, Yann Herault

## Abstract

Down syndrome is caused by trisomy of human chromosome 21 (Hsa21). The understanding of phenotype-genotype relationships, the identification of driver genes and various proof-of-concepts for therapeutics have benefited from mouse models. The premier model, named Ts(17^16^)65Dn/J (Ts65Dn), displayed phenotypes related to the human DS features. It carries an additional minichromosome with the *Mir155 t*o *Zbtb21* region of mouse chromosome 16 (Mmu16), homologous to Hsa21, encompassing around 90 genes, fused to the centromeric part of mouse chromosome 17 (Mmu17) from*Pisd-ps2/Scaf8* to *Pde10a*, containing 46 genes, not related to Hsa21. Here, we report the investigation of a new model, Ts66Yah, generated by CrispR/Cas9 without the genomic region unrelated to Hsa21 on the minichromosome. As expected, Ts66Yah replicated DS cognitive features. However, certain phenotypes related to increased activity, spatial learning and molecular signatures, were changed suggesting genetic interactions between the *Mir155-Zbtb21* and the *Scaf8-Pde10a* interval. Thus, Ts66Yah mice have a stronger construct and face validity for mimicking consequences of DS genetic overdosage. Furthermore, this report is the first to demonstrate genetic interactions between triplicated regions homologous to Hsa21 and others unrelated to Hsa21.

## INTRODUCTION

Knowledge of the pathophysiology of Down syndrome (DS), commonly known as trisomy 21, has been acquired from mouse models. Animal research has allowed the identification of several major driver genes linked to the clinical features found in people with DS, such as DYRK1A and CBS, and made possible the pre-clinical validation of therapies with several drug candidates (Duchon and Herault 2016; Herault *et al*. 2017; Nakano-Kobayashi *et al*. 2017; Faundez *et al*. 2018; Neumann *et al*. 2018; Nguyen *et al*. 2018; Marechal *et al*. 2019). Over the last decades, even more complex mouse models have been generated carrying one human chromosome 21 almost complete (O’Doherty *et al*. 2005; Kazuki *et al*. 2020), and various segmental duplications of regions homologous to human chromosome 21 (Hsa21; Hsa for Homo Sapiens) making it possible to dissect genotype-phenotype relationships in DS (Herault *et al*. 2017; Duchon *et al*. 2021).

These studies were pioneered and strongly driven by the use of the Ts(17^16^)65Dn/J (Ts65Dn) mouse line (Davisson *et al*. 1990; Davisson *et al*. 1993; Reeves *et al*. 1995) with more than 500 publications found in the PUBMED database (April 2022). Originally, the Ts(17^16^)65Dn/J line was generated by the irradiation of DBA/2J males, then crossed with C57BL/6J females, with the progeny being checked for chromosomal abnormalities (Davisson *et al*., 1990; Davisson *et al*., 1993). Translocations for mouse chromosome 16 (Mmu16 for Mus musculus) homologous to Hsa21 were then mated with the C57BL/6J x C3H/HeJ F1 hybrid line. The genetic background of B6C3HF1 was selected to obtain large litters and maintain higher transmission. The translocated minichromosome Ts(17^16^)65Dn/J, encompasses 90 protein coding genes (PGCs) from *Mir155* to *Zbtb21*, linked to the centromeric part of the Mmu17 chromosome which contains a region non-homologous to Hsa21, from Pisd-ps2 to *Pde10a*, with approximately 46 PGCs (MuÑiz Moreno *et al*. 2020)). The region Pisd-ps2 to *Pde10a* is homologous to the region between *SCAF8* and *PDE10A*, found on human chromosome 6. It encompasses many genes involved in neuronal function (*CEP43, NOX3, PDE10A, RNASET2b, SERAC1*, TULP4), in genetic disorders : ARID1B, Coffin-Siris syndrome 1 (www.orpha.net; ORPHA:1465) or in the microdeletion syndrome 6q25.2–q25.3; (ORPHA:251056)(Nagamani *et al*. 2009); *SERAC1* deficiency and MEGDEL syndrome (ORPHA: 352328); Recessive mutations of *GTF2H5* and trichothiodystrophy (ORPHA:33364); RSPH3 involved of primary ciliary dyskinesia (ORPHA:244), and *PDE10A* linked to infantile movement disorders (ORPHA:494541 and 494526).

Due to the trisomy of the Pisd-ps2 to *Pde10a* region in the Ts65Dn, the genetic validity of the model has been a topic of debate for quite some time. On the one hand, the presence of a supernumerary freely segregating chromosome may contribute to producing additional phenotypes as compared to models with intrachromosomic duplications (Goodliffe *et al*. 2016; Olmos-Serrano *et al*. 2016); on the other hand, the presence of 60 genes non-homologous to Hsa21 may have a phenotypic impact not related to DS, and too often neglected in various studies. Somehow, the effect of this additional triplicated segment has been set aside and its contribution to Ts65Dn phenotypes remains undetermined.

To solve this dilemma and have a model closer to DS, we developed a new line named Ts66Yah, derived from the Ts65Dn lineage but no longer carrying the duplicated centromeric part of Mmu17. Here, we report its first phenotypic and molecular characterization.

## RESULTS

### Structure of the *Scaf8*-*Pde10a* proximal region of mouse chromosome 17

Before creating the Ts66Yah model we looked at the *Scaf8*-*Pde10a* region. We found it quite rearranged in several mouse lines according to the mouse genome informatics database. In particular we closely investigated the corresponding segments in the DBA/2J line used to generate the first Ts65Dn chromosome, and also in C57BL/6J and C3H/HeJ (as a proxy) of the mouse lines used to breed the Ts65Dn mouse carriers. Overall, the region was globally conserved in its organization but several loci were affected (Fig. S1A). More precisely, the Snx9 locus was different in size in the 3 models, and more perturbed in C3H/HeJ. Remarkably, the genetic interval encompassing Tulp4, Tmem181a and Sytl3 were not found in the DBA/2J genome and other loci for Rpsh3 and Rnaset2 were duplicated differently in the 3 genetic backgrounds (Fig. S1B). Interestingly, the region is well preserved in humans on Chromosome 6 with an organization similar to that of C57BL/6J, except for large inversions near the *Pde10a* and Rps6ka2-Rnaset2 genes that occurred during evolution.

### Creation, validation, and transmission of the new Ts66Yah minichromosome

Thus, we decided to remove the centromeric segment of Mmu17 located on the Ts65Dn minichromosome using the CrispR/Cas9 technique. Embryos obtained from in vitro fertilization, taking sperm from selected fertile males from the Ts65Dn “1924” line (Shaw *et al*. 2020) and wild-type F1B6C3B oocytes, were injected with CrispR/Cas9 and the pairs of selected gRNAs based on the CRISPOR score (Concordet and Haeussler 2018)(Fig. 1A). One founder carrying the recombined minichromosome, with the deletion of the centromeric part of Mmu17, was selected in the progeny and crossed with C57BL/6NCrl females. Two offspring, one male and one female, were used to start the new colony. The extent of the deletion was characterized by Sanger sequencing of the PCR fragment encompassing the deleted region (Fig. 1B-C), leaving a piece of Mmu16 encompassing the *Mir155* to the end of the telomeric Mmu16 (about 13,856,661 bp). We characterized the new breakpoint between the genomic base number 3,071,436 on Mmu17 and the base 84,354,894 on Mmu16 (UCSC Genome Browser on Mouse Dec. 2011 (GRCm38/mm10) Assembly).

**Fig. 1.**
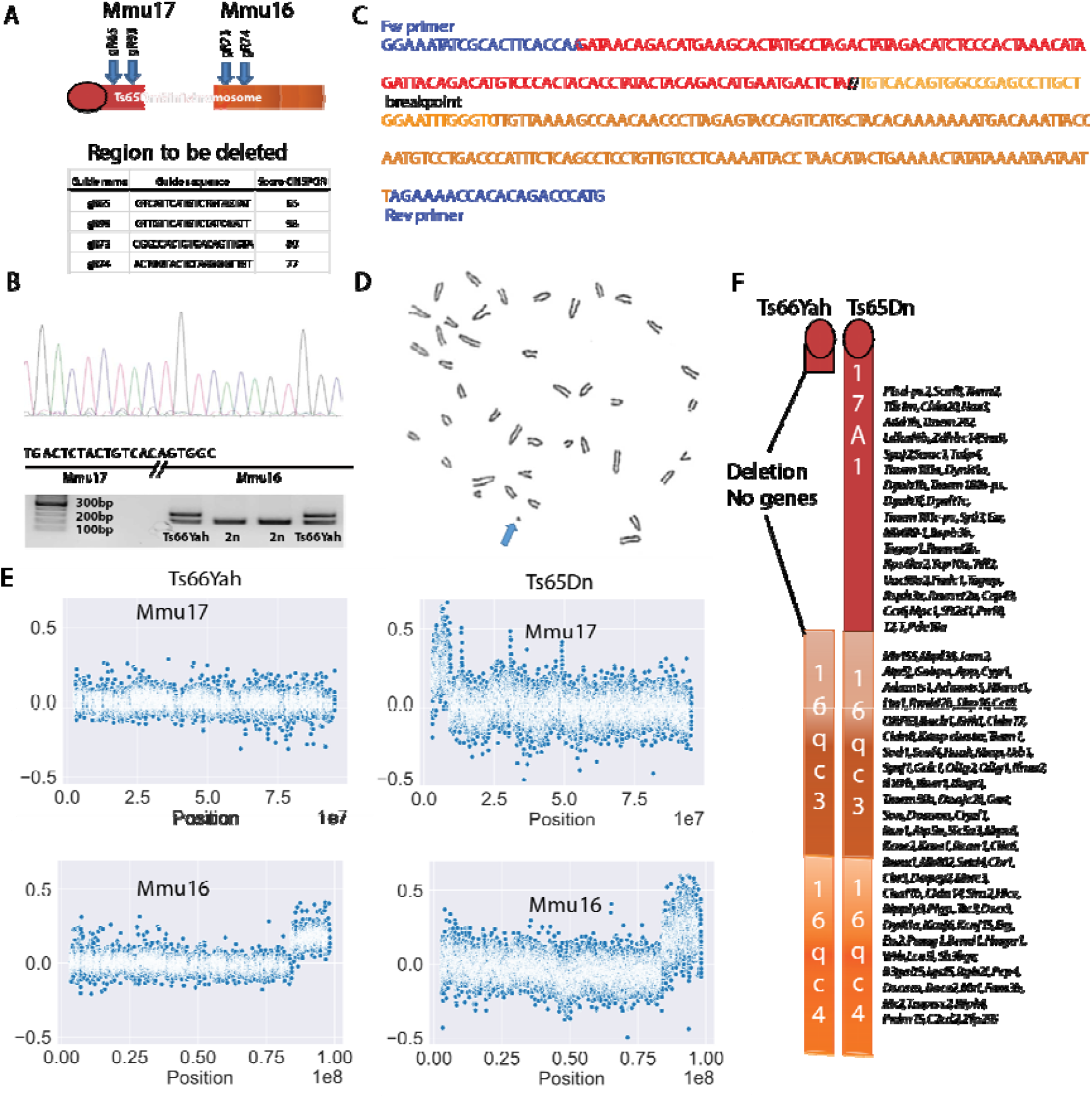
Generation and validation of the new Ts66Yah mouse model. A. Representation of the deletion produced in Ts65Dn using CrispR/Cas9 and two pairs of gRNA. B. Sequence electropherogram and PCR amplification products from the genotyping of Ts66Yah mice. C. Genomic sequence of the new junction found in the deleted minichromosome of Ts66Yah mice. D. One metaphase spread showing the presence of an additional minichromosome in Ts66Yah fibroblast. E. Comparative genomic hybridization (log2) of genomic DNA from Ts66Yah mice versus wild-type (180K probes) compared to Ts65Dn mice (2 100K probes). F. Comparison of the Ts66Yah and Ts65Dn minichromosomes.

We confirmed the presence of an independent chromosome on metaphase spreads in Ts66Yah embryonic fibroblasts isolated from 14.5 days post coitum (dpc) mutant embryos (Fig. 1D). We also performed a-CGH to check the copy number of the Mmu16 and Mmu17 chromosomes both in the Ts66Yah and Ts65Dn models (Fig. 1E). We were able to confirm that the Ts66Yah model indeed lacks the increase in gene dosage for the telomeric part of the Mmu17 region from Pisd-ps2 to *Pde10a* seen in the Ts65Dn model (Fig. 1E-F), while the copy number of the region located on Mmu16 homologous to Hsa21 increased. Moreover, we checked that the change in the Mmu16 copy number in the Ts66Yah mice was located upstream of *Mrpl39* (D_UCHON_ *et al*. 2011), as expected, between two probes targeting the Mmu16 genetic interval 84,325,686-84,356,080 from the mouse genome assembly GRCm38/mm10.

We also checked the genetic transmission of the minichromosome. After establishing the line, the transmission rate (Table 1) began to stabilize at about 30% on both F1B6C3B and C57BL/6J (B6J) genetic background. We thus decided to maintain the mixed genetic background used for the Ts65Dn mice, with the sighted C3H (C3B) males crossed with B6J females, namely B6C3B, that we commonly use in our laboratory. Currently, we have a stable ratio of transmission from both male and female germlines (Table 1), although not all the Ts66Yah males are fertile. We hypothesize that the increased fertility of Ts66Yah was a consequence of the selection of fertile Ts65Dn males when generating the model. To test this, we performed a sperm analysis to evaluate the quality of the sperm by looking at different parameters such as the concentration of spermatozoids, their motility, their velocity and progressivity, for both Ts66Yah and Ts65Dn lines (Fig. S2). As expected, the Ts65Dn mice showed poorer performance than wild-type littermates for all the sperm parameters analyzed. Similar differences were also observed in the Ts66Yah male sperm, with lower quality than the wild-type (wt) littermates. Nonetheless, we were able to isolate 31 fertile individuals out of the 49 males, tested for fertility, in the Ts66Yah line compared to 3 fertile individuals out of 39 males in Ts65Dn, which is below the ranges reported previously for Ts65Dn in two centres (Moore *et al*. 2010).

**Table 1:** Transmission of the Ts66Yah minichromosome in the F1B6C3B and B6J genetic backgrounds. (*some crosses were used during two calendar years and only count once for the total)

| Genetic Background | Period | Transmission rate | Individuals | Litters | Breeding Crosses | Transmission rate | Individuals | Litters | Breeding Crosses |
| --- | --- | --- | --- | --- | --- | --- | --- | --- | --- |
| **B6C3B** |  | Female Ts x wt male |  |  |  | Male Ts x wt females |  |  |  |
|  | 2017 | 44% | 27 | 5 | 2 | 35% | 49 | 5 | 2 |
|  | 2018 | 33% | 87 | 21 | 6 | 25% | 71 | 16 | 6 |
|  | 2019 | 29% | 70 | 14 | 8 | 34% | 120 | 14 | 4 |
|  | 2020 | 26% | 295 | 69 | 17 | 36% | 36 | 3 | 4 |
|  | 2021 | 37% | 349 | 75 | 17 | 31% | 45 | 7 | 6 |
|  | 2022 | 36% | 215 | 37 | 8 | 33% | 66 | 9 | 4 |
|  | **Total** | **33%** | **1043** | **221** | **58\*** | **34%** | **387** | **54** | **26\*** |
| **B6J** |  | Female Ts x wt male |  |  |  | Male Ts x wt females |  |  |  |
|  | **2018-2019** | **30%** | **51** | **9** | **4\*** | **32%** | **53** | **10** | **3\*** |

### The Ts66Yah mice displayed lower locomotor activity compared to the Ts65Dn mice in many tests

To study the behavioral phenotypes under the same environmental conditions, we produced separate cohorts for the two DS models from crosses of wt B6C3B males with Ts66Yah or Ts65Dn females. Then, we challenged the Ts66Yah, Ts65Dn and wild-type littermate animals to a battery of behavioral tests.

First, we assessed their adaptation to a novel environment and exploration activity in a brightly illuminated (120 Lux) square open field. The Ts66Yah mutant mice showed the same pattern of exploration as their wild-type littermates, with no significant differences between the two groups for either the distance travelled or the number of rears (Fig. S3A). In contrast, the Ts65Dn mice travelled further, notably more in the peripheral zone, and performed fewer rears in the central zone compared to their wild-type littermates. These results therefore indicate that only the Ts65Dn mutants, but not the Ts66Yah, presented an increase in anxiety-like behavior, as emphasized by the reduced activity in the center of the arena, and the increased activity in the peripheral zone of the open field.

We then wondered if the circadian activity of the mutant animals was altered, as reported previously for the Ts65Dn model (Ruby *et al*. 2010). We therefore tested the activity of the mice for 46 hours (2 nights). Ts66Yah and Ts65Dn mutants exhibited a similar pattern of circadian activity compared to their control littermates (Fig. S4A). However, we observed that the distance travelled by the Ts65Dn mutants was increased in the light and dark phases compared to the control mice, whereas the activity level of the Ts66Yah mice was comparable to the controls. This result was confirmed by the number of rears, which was also increased in the Ts65Dn line, even in the habituation phase. Similar increases in activity were observed in the number of Y Maze arm entries (Fig. S3C). Thus, the hyperactivity phenotype detected in the Ts65Dn mutants in several paradigms was absent in the

### Ts66Yah. Robust cognitive deficits in the Ts66Yah mouse models

Several learning and memory deficits have been described in the Ts65Dn mouse line by different teams, and are considered as robust and reproducible phenotypes (Reeves *et al*. 1995; Martinez-Cue *et al*. 2002; Fernandez *et al*. 2007; GarcÍa-Cerro *et al*. 2017; Aziz *et al*. 2018; Duchon *et al*. 2021). Thus, we tested how much those robust phenotypes were present in the Ts66Yah individuals.

First, we started to analyze alterations linked to the normal innate behavior of rodents with the nest building task, which is sensitive to hippocampal lesions and is an ancillary parameter assessed to predict cognitive defects (Heller *et al*. 2014). The organization of the nest was scored after one night with a scale from 0, equivalent to the absence of a nest, to 5 when a full dome is raised. For both lines, most control disomic mice built a nest with a dome (score higher than 3; Fig. 2A) nevertheless the nesting score was lower for both trisomic lines compared to the control littermates. Intriguingly, 9 out of 25 Ts66Yah males built a nest with a score above 3 while only 2 males out of 12 reach this stage in the Ts65Dn. Further investigation would be needed to confirm this observation.

**Fig. 2.**
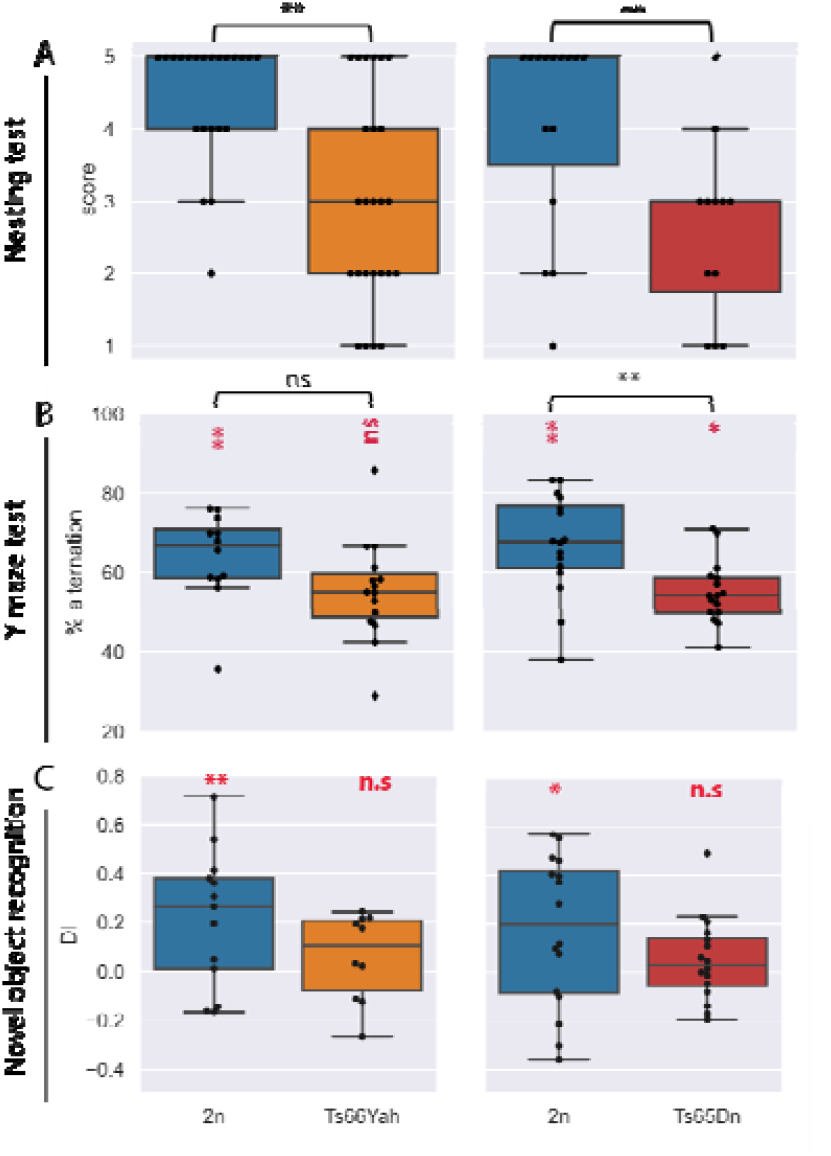
Nesting activities and working memories display similar changes in males from Ts66Yah and Ts65Dn models. A. Mice with trisomy showed deficits in building a nest while a majority of 2n mice were able to build a nest (22 2n with 25 Ts66Yah males and 15 2n with 12 Ts65Dn males). B. Although there was no significant difference in the percentage of alternation between 2n and Ts66Yah males, level was significantly above 50% (chance level) only in 2n and not inTs66Yah mice. Conversely, the Ts65Dn line showed a strong difference in males, with a significant lower % of alteration for the trisomic group as compared to chance level, as well as a reduced spontaneous alternation compared to 2n mice. C. In the NOR, the discrimination index analysis indicated that Ts66Yah and Ts65Dn males were not able to distinguish the novel object (DI was close to 0). Box plots with the median and quartiles. Statistical significance of differences between genotype was inferred by a two-tailed t-test or Kruskal-Wallis non-parametric test (table S1), * p<0.05. **p<0.01. ***p<0.001. Two tailed one sample t-test result versus 0 for DI in C. or versus 50 for Y maze in B., was indicated in red.

Then, we assessed spatial working memory by placing the mice in a Y-maze and left them to freely explore the three arms. The total number of entries to any arm was scored as an index of locomotor activity. In the Ts66Yah mouse line, the percentage of alternation of the mutant mice was significantly different from chance level but no strong difference was observed between trisomic and control individuals. On the contrary, there was a strong difference for the Ts65Dn mutants who showed a lower percentage of spontaneous alternation than their control littermates, with a percentage of alteration for the mutant group significantly different from 50% (Fig. 2B).

Afterwards, we evaluated the episodic non-spatial recognition memory using the novel object recognition (NOR) paradigm with a retention time of 24h. During the acquisition session, there was a difference in the exploration time between both Ts65Dn and Ts66Yah mutant animals versus their respective control groups (Fig S3D). Indeed, both DS model mice spent more time to explore the objects. During the test session, 24 h after the acquisition session, when one of the two familiar objects was replaced by a new one, both DS mice models explored both objects at the same rate and did not show any preference for the novel object contrary to the control mice (Fig. 2C). The Y-maze and object recognition phenotypes were reproduced in independent groups of male and female mice, bred and tested in a second laboratory. Both trisomic males and females showed the same defects in the Y maze and the novel object recognition without any sex effect (Fig. S5).

In the next step, we focused on the spatial reference memory in the Morris water maze (MWM) task. Mice must learn how to escape from a circular pool of opaque water by localizing a hidden platform (PF), set at a single fixed location, using distal spatial cues. Three different sessions were organized according to the diagram shown in Fig. 3A. First, a standard learning phase session where a hidden platform is presented was performed, followed by a reversal session to detect memory flexibility. Both sessions ended with a probe test. Finally, a visible session was performed to assess the general capacity of the mice to perform the test and asses any visual or physical impairments. We measured the velocity, thigmotaxis and time spent by each individual to reach the platform. For the Ts66Yah line (Fig. 3B-I), there was no significant difference between the mutant and control group for thigmotaxis and swimming speed; however, the Ts66Yah mutants took longer to reach the platform during the sessions, indicating poorer performance than the control group. Apart from this, Ts66Yah mice learned the PF location at the last block of the trial with half the latency as over the first block of the trial. The same profile was observed in the reversal phase.

**Fig. 3.**
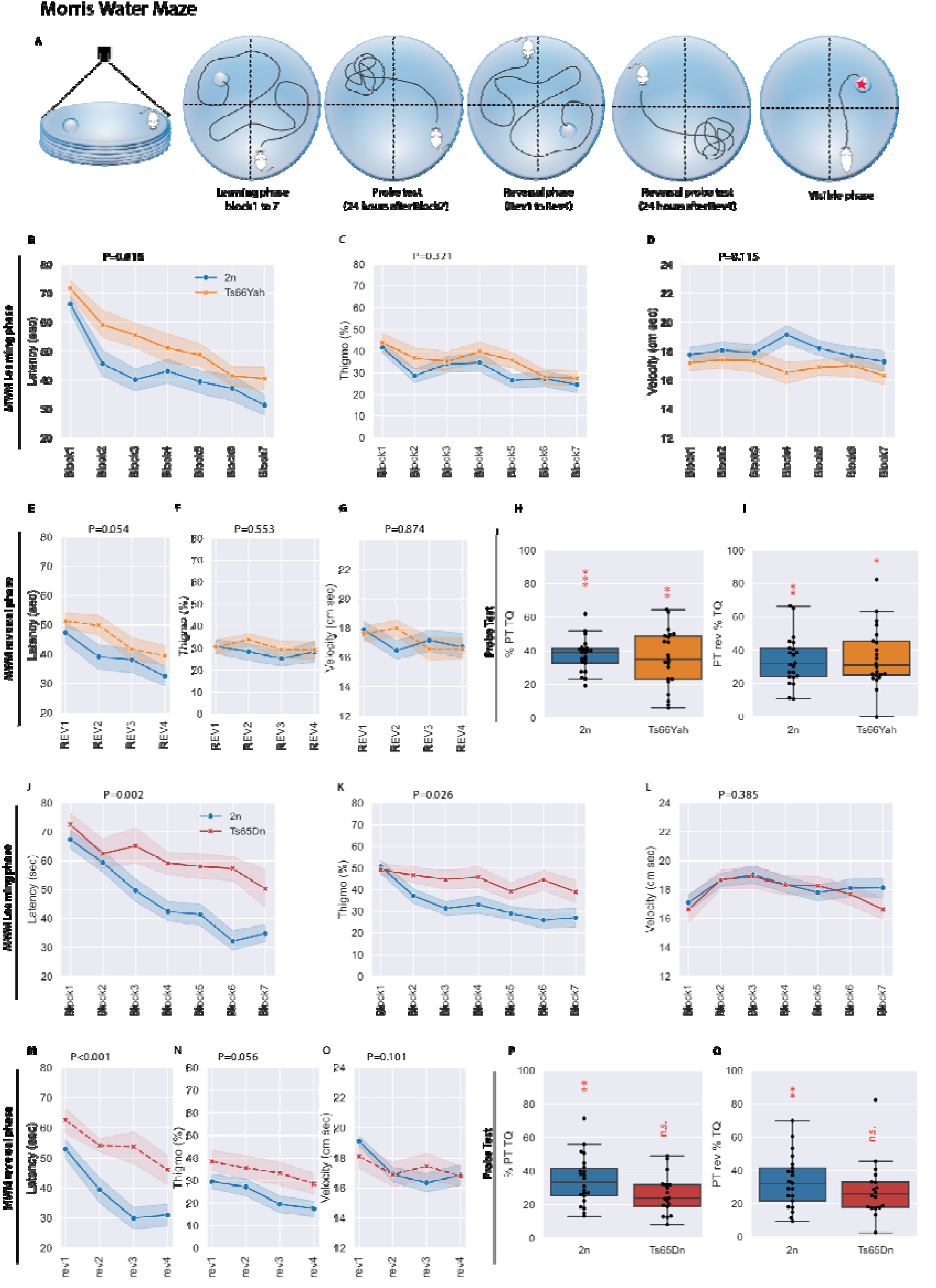
Different effects on spatial learning and memory in the Morris water maze between males from Ts66Yah and Ts65Dn mice. A. Schematic representation of the test. B and J. Both trisomic mice (22 2n with 22 Ts66Yah and 20 2n with 18 Ts65Dn) exhibited a delay in acquisition during the learning phase of the test, resulting in increased latency to find the PF. In addition, the Ts65Dn mice presented increased thigmotaxic behavior (C and K) while velocity was stable regardless of the genotype (D and L). In the probe test (H and P), only the Ts65Dn mice did not present increased exploration of the target quadrant, indicating a clear deficit in reference memory not observed in Ts66Yah mice. During the reversal phase, the augmented latency to find the OF was close to the significant level for the Ts66Yah mice (E), whereas the difference was clearly increased for the Ts65Dn mice. M. There was no difference in velocity (D,L and L,O) for both lines and the thigmotaxic behavior was not found in the Ts66Yah males compared to Ts65Dn (C,F versus K,N). For the reversal probe test, once again only the Ts65Dn mice did not present preferences for the target quadrant (I.Q). Box plots with the median and quartiles. Statistical significance of differences between groups was inferred by an ANOVA for repeated measures (table S1) * p<0.05. **p<0.01. ***p<0.001. One sample t-test result versus 25% is indicated in red in the graph for PT.

The retention memory of place location was assessed during a single probe trial (PT) with no platform available, 24h after the last training session. In this probe test, all the mice remembered where the platform was located after the two learning sessions (learning and reversal phase). As expected, the Ts65Dn mutants (Fig. 3J-Q) presented an increased level of thigmotaxis compared to the controls and an increased latency, indicating that the mutants had difficulties in learning the PF location. This conclusion was supported by the analysis of the reversal phase data. In addition, the Ts65Dn mutants did not present any preference for the target quadrant, with a level of time exploration close to 25% for both the learning and reversal phases.

Efficacy depends on swimming strategies to search for the PF. Thus, we analyzed the strategies to search for the PF undertaken by each individual along the test phase. Only the Ts65Dn mice showed a low and constant high level of non-spatial navigation while the level of spatial strategy increased both in the 2n mice and in the Ts66Yah animals during learning in the sessions of both the initial learning and reversal phase but not in Ts65Dn mice (Fig. S4C). In the visible session (Fig. S4C), the Ts66Yah latency to reach the PF was higher that of the control mice, with a wider distribution. However, the mean of the latency to reach the platform was statistically inferior in both lines (Fig. S4C) to the mean of the latency in the last session of the reversal phase when the platform was hidden (Fig. 3I or 3Q, P<0.001). This indicates that the mice did not have any deficit that could have prevented them from learning and passing the test. Overall, our results confirmed a severe impaired spatial learning in Ts65Dn mice but not in the Ts66Yah mice.

Finally, both mutant lines were tested for contextual associative memory deficits in the Pavlovian fear conditioning test. To induce fear responses, the mice were placed in a “fear context”, a context box where they were subjected to an electric shock. When the animals were re-subjected to the shock 24 h later in the same “fear context”, the level of freezing in all groups was increased compared to the habituation session, indicating that all groups developed a behavioral fear response during the training session (Fig. S4B). The Ts65Dn mutants presented a lower level of responses than the control group, especially at the end of the context session (the last 2 minutes of the session named CONT3). On the contrary, the Ts66Yah mutants did not show any difference in freezing time compared to the control group. Thus, we concluded that the hyperactivity of Ts65Dn mutants interfered with this test: the Ts65Dn mutants would not be subject to a memory failure but instead could not stop moving during a session lasting 2min.

Identification of discriminating phenotypic variables between Ts66Yah and Ts65Dn Considering the differences observed in several phenotypic behavioral variables (Table S1) we wanted to assess the importance of each variable for the genotype-based classification using GDAPHEN (see Materials and Methods). First, we found that several variables considered for the analysis (Table S2) were highly correlated (more than 86% correlation) and thus were removed from the respective downstream analysis for both Ts65Dn and Ts66Yah with their respective wild-type controls (Table S3). The statistical classifiers GLM-Net and RF identified several variables that discriminated the trisomic from the control individual in each model separately (Fig. 4). Using both classifiers, the three most important variables to discriminate Ts66Yah versus wild-type individuals were the “sperm concentration”, the “sperm:progressive”, and the “percentage of time spent in the target quadrant in the 1^st^ probe test” (MWM:PTRev_TQ; GLM-NET; table S4) with the “nesting score” (RF; table S4; Fig. 4A). In addition, the wt and trisomic individuals were easily separated in principal component analysis (PCA; Fig. 4B) with 67% of the variance explained with the 3 first dimensions (Fig. 4C). To discriminate Ts65Dn versus wild-type individuals, the situation was different with both classifiers (Fig. 4D, table S4). The most important variables were also the ”“FC:Precue1 (GLM-Net) and the “sperm concentration” (RF), then the “FC:Cont3” for both and in third position “MWM:PT1_TQ” (GLM-Net) with the “FC:Precue1” for RF (table S4; Fig. 4D). About 59% of the variance was explained with the 3 first dimensions of the PCA to separate Ts65Dn and wt individuals (Fig. 4F).

**Fig 4:**
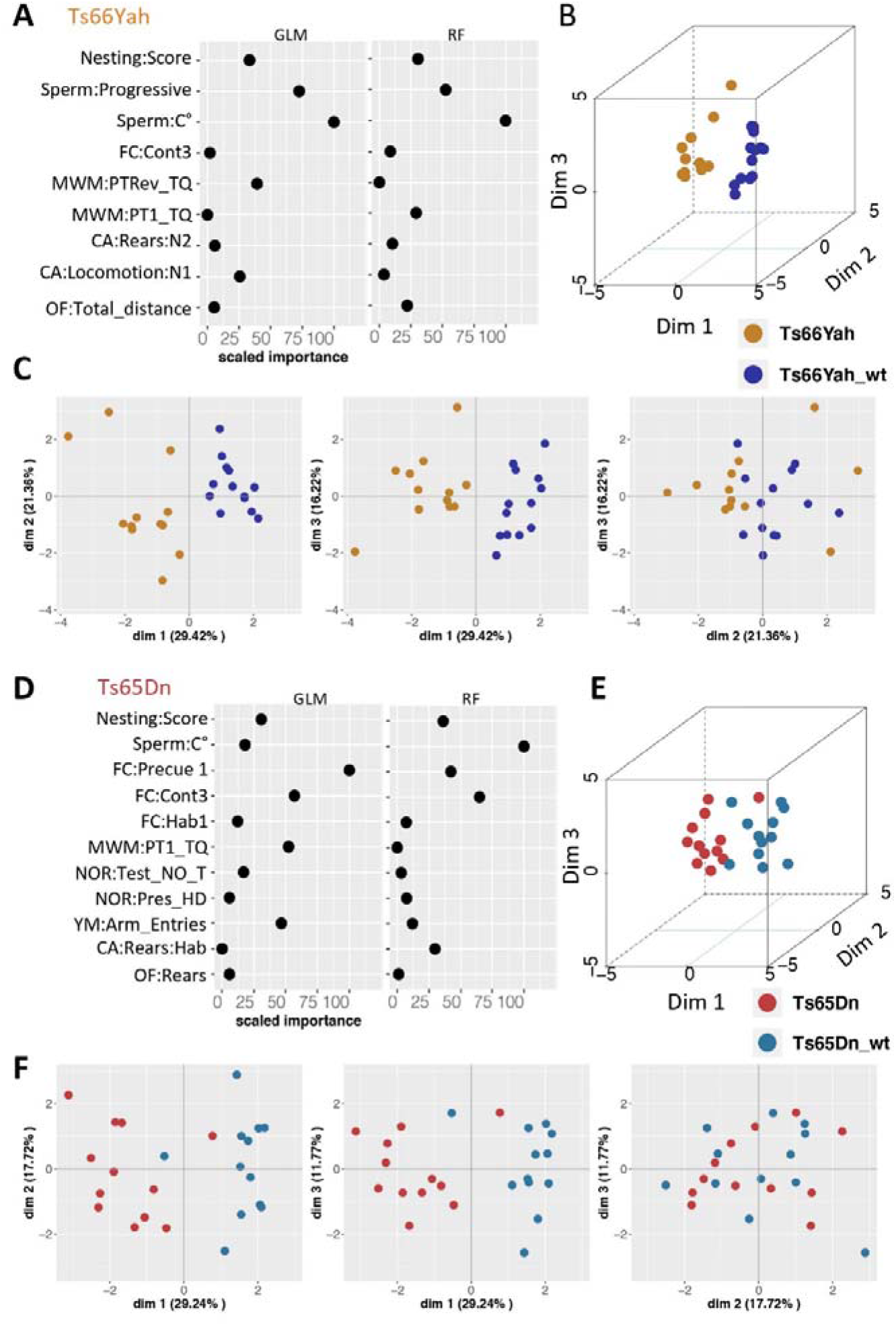
Identification of the strongest phenotypic variables contributing to the genotype discrimination in males from the Ts66Yah (A-C) and Ts65Dn (D-F) models. A-D. Importance of each explanatory phenotypic variable in the genotype discrimination. The selected variables were those known to contribute more than 30% to the genotype discrimination. All measures of importance are scaled to obtain a maximum value of 100 for the variable contributing most to the discrimination in the comparison of Ts66Yah DS mutants versus Wild-types. B-C. 3D-PCA plots showing the individual animals clustering on the 3D space based on the PCA analyses performed with all the phenotypic variables and colored based on genotype and model as follows: in dark blue the Ts66Yah wild-types and in yellow Ts66Yah DS mutants. C-F. Individual component map. The distribution in 2D space of the individual observation coordinates calculated based on the PCA analysis performed after the MFA of the MFAmix function.

Then we analyzed the discriminating variable combining all the genotypes (Table S4a, Fig. S6). We found more behavioral variables to discriminate the wt and trisomic individuals. Apart from the first one, “Sperm:Progressive”, the next six variables were “FC:Hab2”, “MWM:PTRev_TQ”, “NOR:Test_NO”, “YM:Spont_Alter”, “CA:Rears:N2”CA:Locomotion:Light”, although the percentage of variance explained was even lower in the 3 first dimensions, with only 46%. The multiple factor analysis (MFA) computed the correlation between the qualitative or quantitative variables grouped by test or ungrouped, and the principal component dimensions. For both DS models, the MFA and the cosine diagrams highlighted almost the same variables as the main contributors to differentiate between Ts65Dn or Ts66Yah and their respective wild-types.

### Comparison of morphological alterations in the brain and the skull of Ts66Yah and Ts65Dn mice

To detect brain morphological alterations of specific regions in the two DS models, we conducted an MRI study. We did not observe statistically significant changes for the whole brain volume. Nevertheless, to be as accurate as possible, we considered the whole brain volume and performed a Z-score standardization (Fig. 5 and S7). Globally, we observed that changes in the morphology of specific brain structures as well as the direction of the changes (increase or decrease of volume) were the same in both DS lines. However, the amplitude of the changes was less severe but statistically significant in Ts66Yah than in Ts65Dn (for example at ventricles, rest of midbrain, fimbria, superior colliculi; Fig. 5 and S7A). Principal component analysis (PCA; Fig. S7B) comparing mutants with their respective 2n controls for both lines, pointed to a more pronounced difference between the Ts65Dn and their controls than for the Ts66Yah mutants and their controls.

**Fig. 5:**
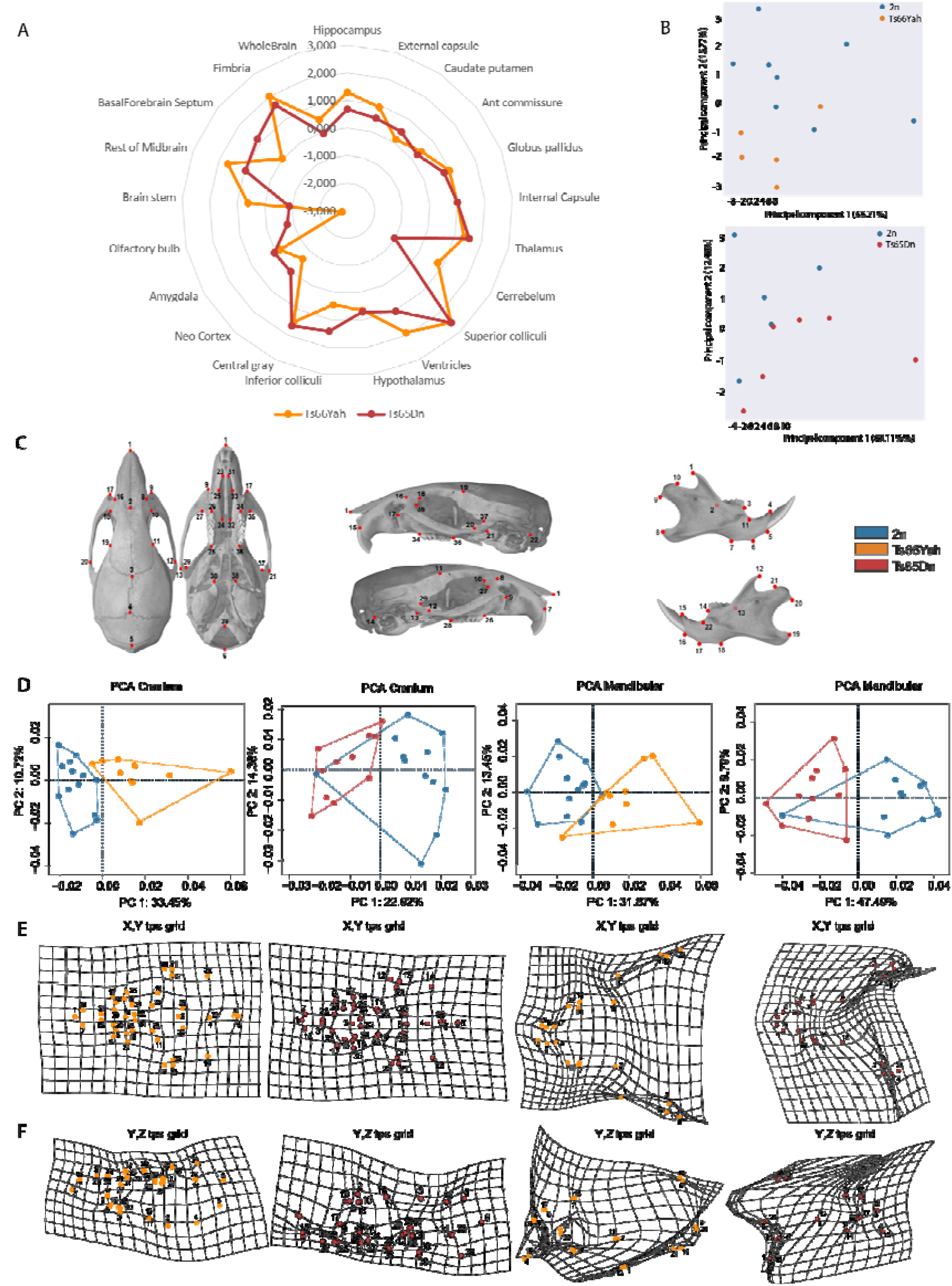
Comparing morphological changes in the brain (MRI) and skull (CT scans) of males from the Ts66Yah and Ts65Dn mouse lines. We analyzed different brain regions/structures taking into consideration the whole brain volume. A. the Z-score was calculated as the mean of control (wt) mice minus the mean of transgenic mice divided by each type of wt and trisomic (Ts). Changes were similar between the Ts66Yah (2n: n=6. 7, Ts: n=7; males) and Ts65Dn (2n: n=5, Ts n=6: males) although the amplitude of the changes was less drastic in Ts66Yah than in Ts65Dn. B. PCA analysis indicated that Ts65Dn were more affected than their respective wt compared to the Ts66Yah line. C. The landmarks used for craniofacial analysis. D-F. Cranio-facial analysis. D. ACP analysis after generalized Procrustes indicated that for cranial skull and mandibular, the 2n group (n=13) and the Ts66Yah (n=10) male group were well separated while the Ts65Dn (n=15) males were less well separated from their control 2n littermates (n=16). E and F. The shape differences between the means of groups was visualized graphically by obtaining the average landmark coordinates for each group and the overall mean and plotting the differences as thin-plate spline transformation grids for the 2 axes. The XY axis was less affected than the YZ axis for both skull and mandible.

People with DS have very specific craniofacial changes which have also been found in Ts65Dn (Richtsmeier *et al*. 2000; Starbuck *et al*. 2014). Thus, we investigated the Ts66Yah, searching for similar craniofacial changes using a landmark based analysis (Table S5, Fig. 5C). PCA computed on Euclidian distance calculated from well-defined landmarks (Fig. 5D) showed that control and trisomic mice were well separated in the Ts66Yah line compared to the Ts65Dn line. The shape differences between groups for skulls and mandibles can be visualized graphically, showing that the deformation was more pronounced in the Y axis than in the X axis (Fig. 5E-F) with the Ts65Dn skull smaller in global size and with more pronounced deformations. Altogether, our brain and skull morphology results demonstrated convergence with DS features in both Ts66Yah and Ts65Dn but with additional changes due to the presence of the trisomy of the *Scaf8*-*Pde10a* genetic interval in Ts65Dn.

### The Ts66Yah and Ts65Dn models show a strong tissue specific dysregulation profile in the entorhinal cortex and hippocampus with differences in functional alterations

The hippocampus (HIP) and the entorhinal cortex (EC) are two brain regions contributing to novel object recognition (Cohen and Stackman 2015; Leal and Yassa 2015; Schultz *et al*. 2015). Because NOR is affected in both the Ts66Yah and Ts65Dn DS models (This study; (Duchon *et al*. 2021)), we wanted to identify genes and molecular pathways altered in the two models and the two brain regions. Thus, we analyzed the expression profile in both the Ts66Yah and Ts65Dn DS models by focusing on the HIP and EC using RNA-Seq. After normalizing the raw data, we were able to confirm the quality of all the biological replicates as the samples clustered well as a function of their genotype using PCA clustering and the Euclidian distance of all the DEGs identified (Fig. S8 A and B). Then, we performed the differential expression analysis (DEA) and identified 1902 and 2220 DEGs in Ts66Yah HIP and EC respectively, and 1836 and 1691 DEGs in the HIP and EC of Ts65Dn (Fig. S8C, Table S6a). All the DEGs found in both tissues were spread along all the chromosomes (Table S6b). Interestingly, the misregulation was very specific in the HIP and the EC, with only 417 and 382 genes found misregulated in both regions (HIP⍰EC) in Ts66Yah and Ts65Dn respectively. First, regarding the Ts66Yah model, the 417 genes identified commonly dysregulated in both tissues were spread along all the chromosomes and more than 50% were non-coding and pseudogene genomic elements. Of these, 72 had a tissue-specific regulatory sense and more than 60% were non-coding and pseudogenes. Moreover, only 54 genes from Mmu16 were identified as DEGs in both tissues. We found similar results for Ts65Dn with 382 common genes between the regions and 34 from Mmu16. Of these 45 have a tissue-specific sense of regulation and more than 70% are non-coding and pseudogenes. Thus, with only around 20% of genes shared between the two brain regions, we concluded that strong tissue specific dysregulation is found in both the Ts65Dn and Ts66Yah models. More globally, we compared the number of DEGs found in both tissues, in Ts65Dn vs Ts66Yah (Fig. 6A). We found 272 common DEGs in the HIP and 282 in the EC (28 are triplicated in Ts65Dn), of these 97 and 105 DEGs follow opposing regulatory senses in both models for the HIP and the EC, respectively (34 are triplicated in Ts65Dn). Those opposing regulated genes could very well be responsible for the differences observed in behavioral analyses.

**Fig. 6.**
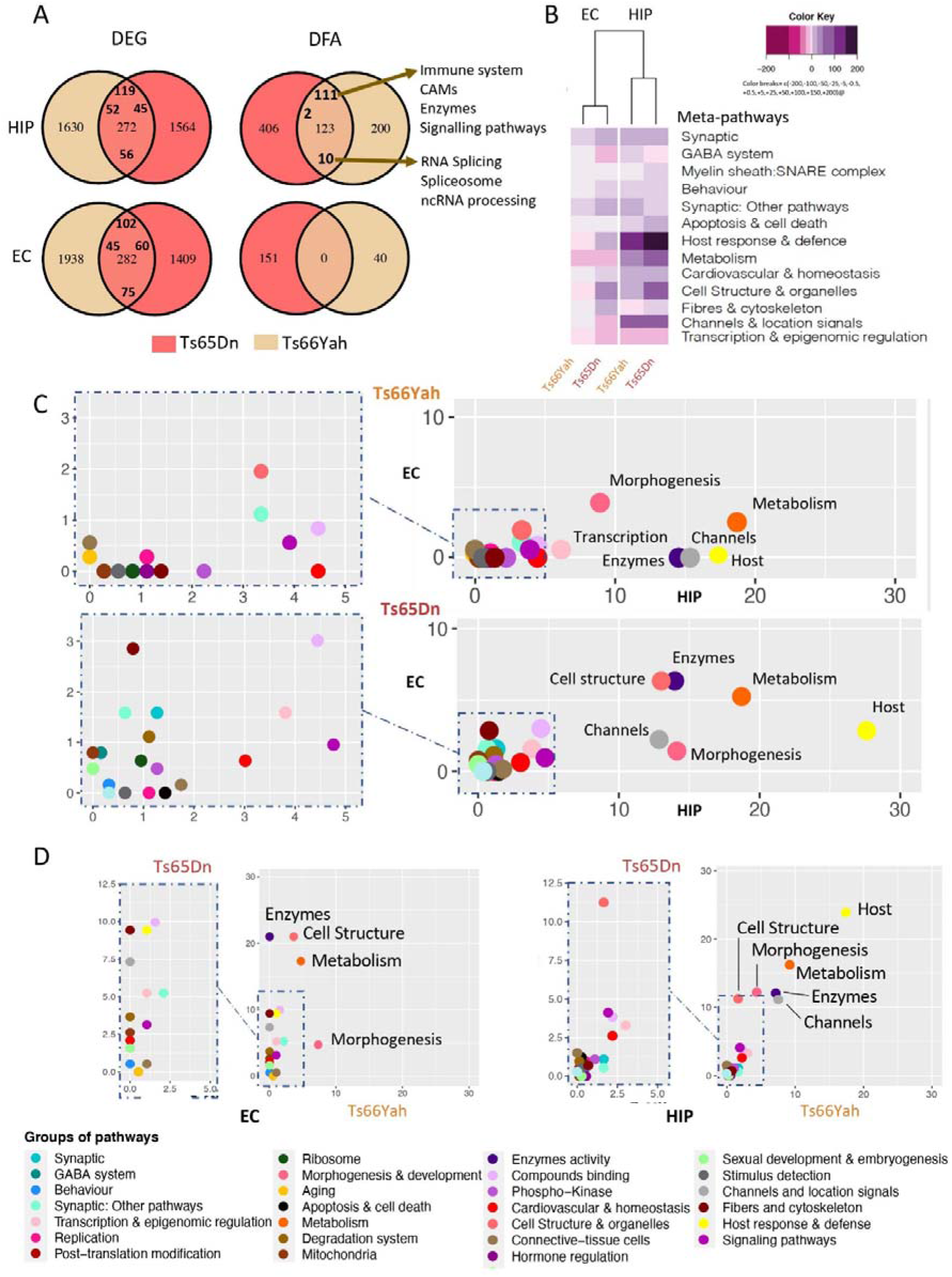
Functional analysis of expressed genes and pathways altered in the Ts66Yah compared to Ts65Dn DS models in male hippocampi and entorhinal cortices. A. Venn diagrams for the DEGs found in common between the Ts66Yah and Ts65Dn hippocampi (HIP) and entorhinal cortex (EC). Right panel highlighting the model specific and common pathways altered between Ts66Yah and Ts65Dn HIP samples in the upper part and the EC datasets of both the Ts65Dn and Ts66Yah models in the lower part. B. Heatmap representation of the number and regulation sense of the meta-pathways found in the Ts65Dn and Ts66Yah HIPand EC. The color key breaks represent the number of pathways within the meta-pathways. C. Scatter-plot showing the inter-tissue comparison of the percentage of pathways included on each meta-pathway, normalized by the total number of unique pathways per meta-pathway for Ts66Yah (upper panel) and the Ts65Dn (lower panel) on the x-axis and y-axis representing the HIP and the EC for the Ts66Yah or Ts65Dn models, respectively. D. Similar representation showing the inter-model comparison with the percentage of pathways included on each meta-pathway (group of pathways) normalized by the total number of unique pathways per meta-pathway found in the HIP of Ts66Yah (x-axis) in comparison with Ts65Dn (y-axis).

Then, we checked the fold change of triplicated genes along the chromosome regions on Mmu16 and 17. We detected a similar fold change for most of the genes located in the Mmu16 region, homologous to human chromosome 21, in the HIP and EC (Fig. S9; Table S7). Out of the 133 protein-coding genes (PCG) that are triplicated and located in the segment on Mmu16, 90 and 66 showed compensated expression in the Ts66Yah HIP and EC (67 and 49% of the PC triplicated genes), whereas 78 and 100 genes were compensated in the Ts65Dn tissues (58 and 75% respectively; Table S6a), thereby supporting previous observations in which approximately half of the triplicated genes are compensated or partially compensated in partial DS models, with a strong effect of the tissue/organ analyzed (AÏt Yahya-Graison *et al*. 2007; Duchon *et al*. 2021). As expected with the dose effect, the fold changes of the 51 genes (PCG and non-coding), located on the proximal part of Mmu17, were around 1.02+/-0.25 and 1.01+/-0.25 (mean+/-SD) respectively in the Ts66Yah HIP and EC, while they were between 1.54+/-1.22 and 1.63+/-1.2 in the Ts65Dn Hip and EC. Altogether the spearman correlation comparing TEGs from Mmu16 in the HIP or EC in both models was low (between 26 to 30%) and was completely loss when looking at TEGs from the centromeric region of the Mmu17 (Fig. S8D). Nevertheless, a few genes from this centromeric interval of Mmu17, triplicated in Ts65Dn but not in Ts66Yah, showed a distinct pattern of expression in the Ts66Yah hippocampi. As expected, 9 genes, Ezr, *Pde10a, Scaf8*, Tiam2, Tfb1m, Zdhhc14, Synj2, Serac1 and Gtf2h5, were differentially expressed in Ts65Dn but not in Ts66Yah; 2 genes, Ccr6 and Tagap, upregulated in Ts65Dn, were downregulated in Ts66Yah.

We verified by qRT-PCR the expression of several genes in the HIP of control and mutant mice. We confirmed the overexpression of Dyrk1a, Sod1 and Sh3bgr as expected in both models. We also verified that *Pde10a* and Ezr from the Mmu17 region were only upregulated in the Ts65Dn model HIP (Fig. S10) while Ccr6 was downregulated in Ts66Yah and upregulated in the Ts65Dn HIP. Finally, we also identified genes from Mmu16, syntenic to Hsa21, with an increase of at least 0.5-fold in Ts65Dn compared to the Ts66Yah HIP, with Kcne1, Kcnj15, Map3k7cl, and in the EC with Sh3bgr, Itgb2l, Ripk4 or Ripply3, suggesting that the increased dosage of the centromeric Mmu17 genes interfere with the regulation of these genes.

On the functional side, we identified 323 and 493 pathways altered in the hippocampi of the Ts66Yah and Ts65Dn models, and 40 and 135 pathways in the EC of both models (Table 3 and table S8). This observation suggested a more profound impact in the hippocampi versus the EC, and in the Ts65Dn versus the Ts66Yah conditions (Fig. 6B). In addition, a milder effect was found in both DS models in the EC samples in comparison with the HIP, and no common pathway was found between Ts65Dn and Ts66Yah in the EC (Fig. 6A). Next, we identified common alterations per brain region in both DS models; in the HIP, 123 common pathways were altered following the same regulatory sense. Of these, 111 were upregulated involving mainly the immune system, cell adhesion molecules and signaling pathways. However, only 10 were downregulated linked to RNA splicing and ncRNA processing (Fig. 6C-D, table S8). The heatmap showing the average number of pathways contributing to meta-pathways with regulatory orientation for the Ts65Dn and Ts66Yah HIP and EC models, supported strong brain region-specific alteration observed in the DEA in both models (Fig. 6B). Interestingly, the samples were clustered by brain regions and not by models. Inside each meta-pathway we identified those that showed a different intensity in the number of pathways altered or in the regulatory sense between both DS models.

In addition, the majority of altered pathways in the HIP for both models were upregulated as expected (Duchon *et al*. 2021). In the EC a lower number of pathways grouped on each meta-pathway were identified compared to the HIP. As such, the HIP showed a higher number of altered pathways and a higher deregulation of “host response”, “channels”, “metabolism” and “cell structure and organelles”. Interestingly the effect was stronger in Ts65Dn compared to Ts66Yah (Fig. 6B). To better understand the region-specific alterations, we compared the HIP and EC meta-pathway maps for each model (Fig. 6C) and between the DS models (Fig. 6D). A few meta-pathways, such as “metabolism”, “morphogenesis and development”, “channels and location signals” and “enzyme activity”, were strongly affected in both DS models, with slightly different levels in the HIP and EC (Fig. 6C). Surprisingly the “cell structure and organelle” were more strongly affected in Ts65Dn than in Ts66Yah, and the “host response” was more affected in both the Ts66Yah and Ts65Dn HIP but only in the EC of Ts65Dn. Moreover, when we compared the HIP and the EC alteration map (Fig. 6D) between Ts66Yah and Ts65Dn, we observed that Ts65Dn showed stronger alterations than Ts66Yah in the “Host and immune response”, “Morphogenesis”, “Metabolism and Cell structure” related pathways in the Hippocampi. In addition, the alteration in pathways in the EC was higher in the Ts65Dn model considering the number of total pathways found altered (151 in Ts65Dn, 40 in Ts66Yah) and involved mainly the “Metabolism”, “Enzymes”, “Cell structure”, “Fibers and Cytoskeleton” and “Host and Immune response” meta-pathways.

To further understand the nature of the Ts65Dn and Ts66Yah phenotype divergence, we built a regulatory and PPI network, noted RegPPINet, using all the genes identified by the GAGE pathway analysis (Luo *et al*. 2009), for Ts65Dn and Ts66Yah, and known to contribute to the synaptic meta-pathway group. After performing a betweenness based centrality analysis on the joined Ts65Dn-Ts66Yah RegPPINet we identified three main sub-networks (Fig. 7A): the “MHC-immune response” gathered members of the major histocompatibility complex (MHC) and of the IFN response, and is almost Ts65Dn specific, whereas the two others, “RHO” and “Morphogenesis”, were more affected in the Ts66Yah model. Furthermore, our network analyses here in Ts66Yah and Ts65Dn support the role of the DS subnetworks linked to RHOA, SNARE proteins (Vamps and Sec proteins interactors), DYRK1A and NPY (Fig. S11), which are deeply intertwined, as previously identified in 7 other DS mouse models (Duchon *et al*. 2021).

**Fig. 7.**
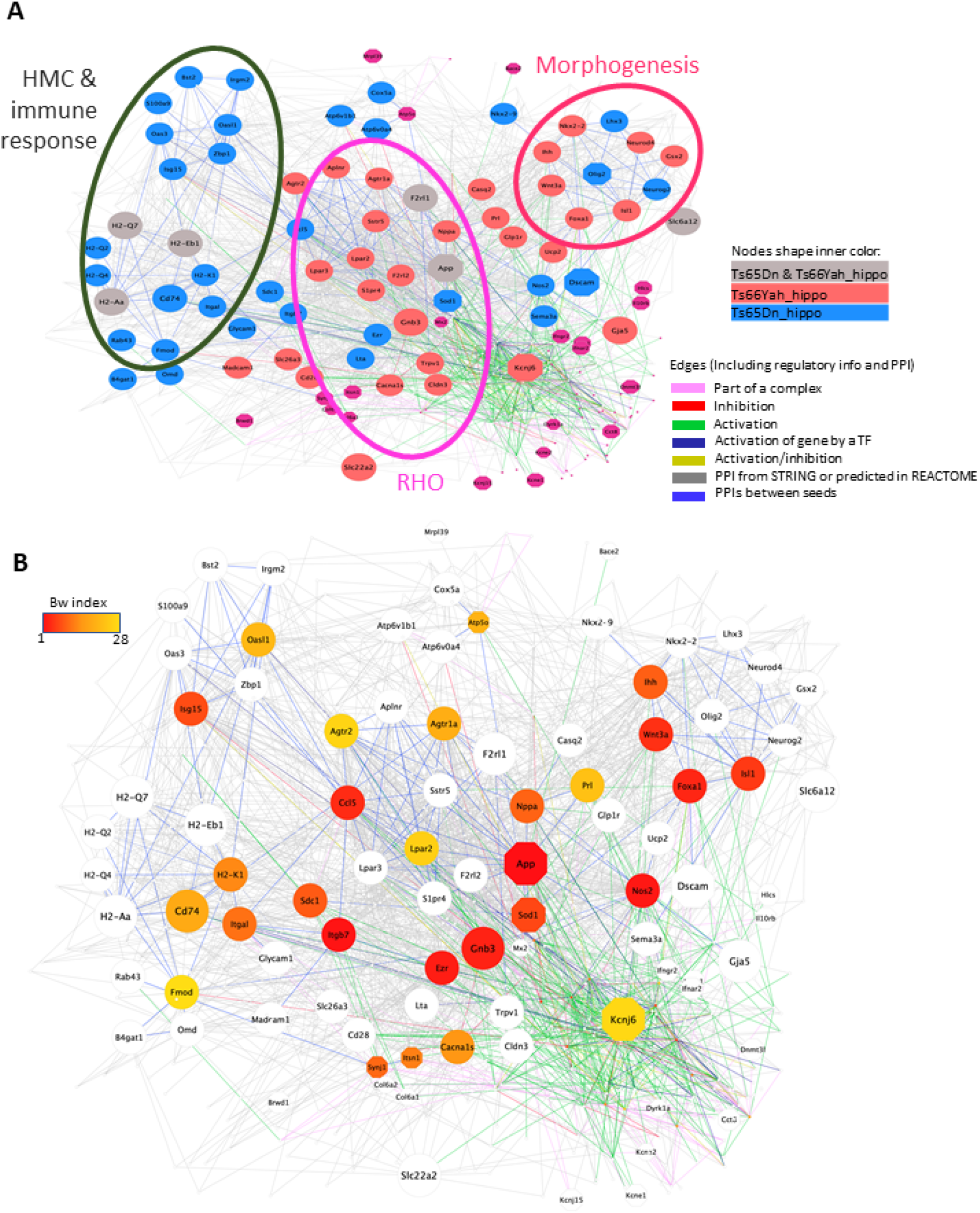
Central Protein-protein interaction and regulatory gene connectivity network (RegPPINets) involved in the synaptic meta-pathway identified in Ts65Dn and Ts66Yah males, highlighting the top 30 genes identified by betweenness centrality analysis. A. Central RegPPINets of genes involved in the synaptic meta-pathway identified in Ts65Dn and Ts66Yah, highlighting the main subnetworks found. B. The same central RegPPINets highlighting in color from red to yellow the proteins more central for the communication flow over the network identified by the centrality analysis using the betweenness index.

Then we focused on the EC and HIP Ts66Yah RegPPINet. Strikingly, the pathway dysregulation in the EC unraveled a mild effect of the trisomic model over this tissue structure with 40 pathways affected, mainly morphogenesis, metabolism, cell structure and synaptic meta-pathways (Fig 6; Fig. S12). The Ts66Yah EC RegPPINet revealed several well-connected subnetworks (Fig. S12B) linked mainly to “morphogenesis”, “metabolism” and “host response & defense” (Fig. S12C). Conversely, a more severe dysregulation with 323 altered pathways was revealed in the HIP (Table 4 and S08, Fig S13A). Although less functional alteration was found in Ts66Yah than in Ts65Dn, the stronger changes were found at the level of “Host response & defense” and “Metabolism” meta-pathways with important relations with genes also belonging to morphogenesis and synaptic related pathways (Fig. S13D). The network analysis identified several central genes distributed over all the subnetworks (Fig. S13B, C) highlighting the strong connectivity and importance of each subnetwork in the Ts66Yah HIP. Additionally, in the most closely connected subnetwork including the top central genes, we identified App and G protein-coupled receptor 3 for lysophosphatidic acid (LPA) LPAR3 and LPAR2, which were closely connected (Fig. S13C).

Furthermore, most of the genes contributed to the alteration in different meta-pathways or functionalities due to gene multifunctionality, again pointing out the relevance of improving our understanding of comorbidity phenotypes in DS (Fig. S13D).

## DISCUSSION

Here we report the general characterization of the new Ts66Yah line and compared several outcomes with the well-known Ts65Dn DS mouse model. Ts66Yah is certainly more genetically valid for DS studies than Ts65Dn as it does not contain trisomy of the *Scaf8*-*Pde10a* centromeric Mmu17 region unrelated to Hsa21. The general transmission of the recombined Ts66Yah chromosome was similar to the Ts65Dn, showing that the centromeric part of Mmu17 does not play a major role in the sterility of trisomic males. The male germline transmission property in the Ts66Yah may have arisen from the construction of the models and should be monitored to see if it is maintained over generations. Interestingly, sperm quality was also similar, thus probably the fertility issue is more connected to sperm fertility as such rather than the parameters of concentration, mobility and velocity checked here.

More importantly, both DS models were found to replicate very classical DS cognitive features. Similarities were found in behavior phenotypes in the two models for the nesting, object recognition and spontaneous alternation tests. In particular, defects in object recognition have been observed in many different labs for the Ts65Dn line and have been replicated in two independent laboratories for Ts66Yah. Thus, we must consider this phenotype as a very robust one. It would be very interesting to see if other phenotypes found in the Ts65Dn are also reproduced in Ts66Yah. For example, the impairment in spatial learning was found in the two models during the MWM test but no spatial memory phenotype was observed in Ts66Yah compared to Ts65Dn. In addition, we found that the Ts66Yah line did not reproduce the anxiety and hyperactivity phenotypes observed in the open field with the Ts65Dn line. Similarly, circadian activity was normal in Ts66Yah whereas we found increased activity for Ts65Dn, as observed previously in light-dark condition (Ruby *et al*. 2010). Although hyperactivity may be influenced by gene-environment interaction (Itohara *et al*. 2015), increased locomotor activity is common to many mouse mutant models, i.e. 678 significant genes out of 8641 mutants tested in the International Mouse Phenotyping consortium. Hyperactivity could be a direct consequence of abnormal involuntary movement or motor coordination, or indirect consequences, such as associated with stress, being in a novel environment or being influenced when raised by or in contact with mutant individuals. This increased activity has a major impact on behavioral testing in Ts65Dn, as shown recently by the contribution of distance travelled and thigmotaxis to novel object discrimination (Sierra *et al*. 2021). This phenomenon should depend on genes located in Mmu17 either directly or indirectly, because Ts66Yah was not affected. According to the Mouse Genome Informatics database, mutants from five genes of the region, Arid1b, Cep43, Nox3, *Pde10a*, and Serac1, are linked to activity, motor and stereotypic behavior and are strong candidates for which overdosage may enhance the activity of the Ts65Dn mouse model. Similarities were also observed in brain and craniofacial morphologies, with convergence found in both DS models. Nonetheless, the severity of the trisomy in these two features were slightly different: The Ts65Dn brain was larger than the Ts66Yah one in many different regions while the shape of the cranium was affected differently, with more severe brachycephaly and increased mid-face shortening in Ts66Yah, compared to Ts65Dn, a phenotype closer to human DS features.

Another intriguing result came from the expression analysis in the two brain regions, the HIP and EC, which are both involved in learning and memory. First, there was little overlap in the DEGs in the two models, and while some pathways were found dysregulated similarly in the hippocampus of both models, no real pathway convergence was observed in the EC, with a more severe impact for Ts65Dn compared to Ts66Yah, presumably due to the trisomy of the centromeric region of Mmu17.

Altogether the results from learning and behavior, craniofacial and brain morphologies, and gene expression in two brain regions, highlighted the interference of the *Scaf8*-*Pde10a* trisomy in the Ts65Dn phenotypes. Considering the breeding of the two models on F1B6C3B, it would be highly recommended to genotype for the zygosity of the *Scaf8*-*Pde10a* region that can be found as heterozygous for B6 and C3B alleles, or homozygotes for both alleles. In addition, it would be important to control for any recombination event in the small region to reduce the risk of generating a subline with specific genotype in this region. Interestingly some gene like EZR, encoded in the Mmu17 interval, is known to interact with the RHOA pathway, and is able to downregulate RHOA activity. Reversely, RHO can activate EZR activity and lead to new specific phenotypes during neuritogenesis (Yonemura *et al*. 2002; Matsumoto *et al*. 2014; Hatano *et al*. 2018). *EZR* was found to be a major hub in the top 5 influential hippocampal genes. We may therefore hypothesize that the EZR overexpression found in the Ts65Dn brain acts on the downregulation of the RHOA pathway found altered in DS models (Duchon *et al*. 2021). For the lower activity phenotypes found in Ts66Yah compared to Ts65Dn, Serac1, *Pde10a* and *Ccr6*, whose loss-of-function impaired locomotor activity, are interesting candidates while the defect in *Kcne1* induced hyperactivity. This means that the dose of those genes may explain the hyperactivity found more severe in Ts65Dn than in Ts66Yah.

In the network analysis of both the Ts65Dn and Ts66Yah hippocampi, we confirmed the dysregulation in the immune response, RHOA and morphogenesis pathways, as detected previously in several DS models (Duchon *et al*. 2021). From this common network, five proteins, encoded by the Mmu16 region homologous to Hsa21, appeared to have stronger effects, based on the betweenness centrality index (APP, SOD1, KCNJ6, SYNJ1, ITSN1) while others had less impact (ATP5o, BACE2, BRWD1, DYRK1A, DSCAM, HLCS, IL10RB, INFAR2, INFGR2, KCNE1, KCNE2, KCNJ15, MRPL39, MX2, OLIG2), supporting the multigenic dimension in DS models. Interestingly, genes from other regions homologous to Hsa21 were also detected in the hippocampal network, like COL6A1, COL6A2 and DNMT3L.

The altered immune response has been reported previously in DS models (Ling *et al*. 2014; Guedj *et al*. 2016; Duchon *et al*. 2021). Here we found that both “synaptic” and “host & immune response” meta-pathways were closely connected by protein-protein interaction between several members and more interestingly, some of the genes had dual synaptic and host response functionality such as App, Cldn19, F2rl1 (involved in the behavioral neurological phenotype), *Ihh, Trpv1, Nppa, Wnt3a, Isl1*, and a few genes coding for cell adhesion molecules, such as *H2-Q6* or *H2-Aa*.

Compared with the parental Ts65Dn, the Ts66Yah line revealed the importance of the *Scaf8*-*Pde10a* trisomy. Interestingly this region varies quite often in the mouse genetic inbred backgrounds used to generate and maintain the Ts65Dn minichromosome. Originally the translocated minichromosome was from DBA/2J but then it was crossed to B6J and later bred to (C57BL/6J x C3H/HeJ)F1 (Davisson *et al*. 1993). Unfortunately, the follow-up of the DBA/2J alleles on Ts65Dn has never been done although there was a proposal to use a SNP in *Snx9* to detect the DBA/2J allele in the proximal part of the Ts65Dn minichromosome (Lorenzi *et al*. 2010). This variation in allelic composition of the *Scaf8*-*Pde10a* region may contribute to the fluctuation of the phenotypes observed from stocks to stocks in the Ts65Dn (Shaw *et al*. 2020). Interestingly Shaw *et al* (2020) demonstrated several differences in the phenotypes of the Ts65Dn mouse lines 1924 and 5252 from different batches that have been further separated in 2010 to correct for retinal degeneration at the Jax laboratory: the 1924 subline keeping the original Ts65Dn genetic background and the 5252 rederived in a F1B6EiC3Sn.BLiAF1/J genetic background. Here too, no tracing of the Ts65Dn genotype was done in the sublines. Nevertheless, our results obtained here from the Ts65Dn 5252 subline (ordered in 2018) are coherent for the MWM memory phenotype found also in the cryopreserved stock in 2010, close to the time the 1924 and 5252 were separated. Thus, the loss of MWM memory phenotype observed in the Ts66Yah mouse compared to the Ts65Dn 5252 subline is indeed a direct consequence of the rescue of the disomy of the *Scaf8*-*Pde10a* region. Nevertheless, more investigation should be done to control the other embryonic phenotypes found altered in the 5252 compared to other Ts65Dn sublines (Shaw *et al*. 2020).

As shown here, DS related phenotypes in mouse models could be altered by change in genetic dosage of another genomic region, the *Scaf8*-*Pde10a* genetic interval. Conversely, this region may change penetrance or expressivity of features in individual with DS. Indeed, the *SCAF8*-*PDE10A* on human chromosome 6 is also subjected to copy number variation according to the DECIPHER database (Firth *et al*. 2009). In human, the region is rearranged in the 3’ part (Fig S1B) and contains the *ARID1B, SERAC1, GTF2H5, RPSH3* and *PDE10A* disease genes. Thus, the additional trisomy of *Scaf8*-*Pde10a* has a strong effect on the expression of several DS related features in models. This phenomenon of additive effect due to genetic interaction has been observed in other copy number variation diseases (Pizzo *et al*. 2019).

Overall it highlights the need of complete evaluation of the genetic background in individual with DS, to define potential interaction with other candidate disease-associated variants and to get a better understanding of the DS complexity. Our report introduced a new model with a freely segregating chromosome with a stronger genetic validity for DS, phenotypic variation in 3 main areas affected in individual with DS. Therefore, we hope that the DS research community will consider working with the newTs66Yah mouse model for DS, although this model remains partial DS model since the triplicated *Mrpl39* to *Zbtb21* region encompass 102/187 of the Hsa21 orthologous protein coding genes. The Ts66Yah model should be reduce interference with the Mmu17 region unrelated to Hsa21 in Ts65Dn, and obtain more refine analysis of DS during lifetime, closer to the DS genetic condition.

## MATERIALS AND METHODS

All the experiments were performed in accordance with the Directive of the European Parliament: 2010/63/EU, revising/replacing Directive 86/609/EEC and with French Law (Decret no. 2013-118 01 and its supporting annexes which came into force on 1 February 2013) relating to the protection of animals used in scientific experimentation. YH was the principal investigator of this study (accreditation 67-369) in our animal facility (Agreement C67-218-40). Experimental procedures for the use of animals for research were approved by the Ministry of National Education, Higher Education and Research and with the agreement of the local ethical committee Com’Eth (no. 17) under the accreditation number APAFIS#13127-2018012210465339 v5 and APAFIS#21969-2019091215444738 v3.

### Mouse lines

The Ts65Dn (Ts(17^16^)65Dn) mice analysed in the study was obtained from the Jackson Laboratory in 2018 from the 5252 subline (Shaw *et al*. 2020), was kept in an F1 B6C3B genetic background (with the C3B line as a C3H/HeN congenic line for the BALB/c allele at the Pde6b locus (Hart *et al*. 2005; Hoelter *et al*. 2008). Ts65Dn mice were genotyped according to published protocols (Duchon *et al*. 2011). Two primers were selected on both sides of the breakpoint to amplify a fragment of 396 bp (with forward primer Fw_wtTs65Dn: GACTTAGTAAGAGCAAGTGGC and reverse primer Rev_Ts65Dn: AGGTAGAAAGATGTGAGGACAC), and a third primer was designed on the reverse strand of chromosome 17 to amplify a fragment of 290 bp (GGGCAACACTGGATCAATC).

The generation of the new mouse line was done based on Ts65Dn “1924” line (Shaw *et al*. 2020). Briefly, Ts65Dn male mice were selected and used for in vitro fertilization to produce fertilized eggs that were injected with two pairs of gRNAs (Fig. 1A) with one pair located on the centromeric region of Mmu17 and the other in the proximity of the breakpoint in the Ts65Dn minichromosome and on the Mmu16 region. The Ts66Yah line was obtained at PHENOMIN-ICS using the CRISPR/Cas9 technology. The CRISMERE approach was selected to specifically obtain the deletion of the 6.2 megabases region located on the Ts65Dn minichromosome. Single guide RNAs were selected using the CRISPOR web site (https://doi.org/10.1093/nar/gky354) in order to generate double strand breaks close to Chr17 centromere; gR65 and gR93 (GRCm39 17:3121550-3122030); at >43 kb from SR-related and CTD-associated factor 8) and 3.8 kb 3’ of the break point of the minichromosome; gR73 and gR74(Chr16:84151598-84152123); at 363 kb of *Mrpl39*-39S ribosomal protein L39, mitochondrial isoform 1) to avoid the repeats that are present at the Ts65 break point. Both guide RNAs and Cas9 mRNA were synthesized by in vitro transcription (Birling *et al*. 2017). Microinjection of CRISPR reactive was performed in the pronuclei of fertilized oocytes obtain after in vitro fertilization of B6C3 F1 females with the sperm of aTs65Dn fertile male. Ninety one oocytes were fertilized and microinjected with CRISPR/Cas9 reactive (the 4 sgRNAs and Cas9 mRNA). Fifty-three embryos that developed in 2-cell embryos were reimplanted in 3 foster CD1 females. Fifteen pups were born and genotyped by junction PCR. One male pup had a clear new junction corresponding to the expected deletion. This male was still positive with the Ts65Dn junction, showing that this animal was mosaic. Noteworthy, out of the 15 pups born, only 3 had the Ts65 minichromosome junction. Droplet digital PCR was performed on the positive pup and a control pup with a probe located on Dyrk1a (located on chr16 and the minichromosome) and Snx9 (located on chr17 and the minichromosome). Two copies of WT were detected with both probes on the WT control animal. On the new junction positive animal, 3 copies were clearly detected with the Dyrk1a probe located on Chr16 while a decrease to 2.5 copies of WT was observed with the Snx9 probe located on Chr17. These results confirmed the fact that the founder was mosaic and the intact Ts65 minichromosome was still present in some cells of the animal.

At 8 weeks of age, the male founder was bred with 2 wild type C57BL/6NCrl females. Both WT females gave birth to a single litter with a total of 13 pups, no other pups were born afterwards. Five pups had the same new junction than observed on the founder whereas the Ts65 minichromosome junction was not observed on any of these F1 pups. For the 5 pups with the new junction, only 2 copies of WT were observed with the Snx9 probe (Chr17) and 3 copies were detected with both *App* and *Dyrk1a* probes (both located on Chr16), confirming the presence of the engineered minichromosome in the pups. The recombined chromosome was selected to propagate upon breeding. As such the full name of the mouse line should be Ts66YahIcs, shortened here to Ts66Yah.

Ts66Yah mice were genotyped with a specific primer encompassing the new break point between Mmu17 and Mmu16, with a forward primer Ts66Yah_wt-tg_up (GGAAATATCGCACTTCACCAA) and a reverse primer Ts66Yah_tg_dw (CATGGGTCTGTGTGGTTTTCT) to amplify a fragment of 322 bp. A third reverse primer was designed on the reverse strand of chromosome 16 to amplify a wt fragment of 234 bp (TCTAGGATCAGTGGGACTTTTGT).

### Metaphase spread

Fibroblasts obtained from 2 Ts66Yah embryos were treated with 0.02 μg/ml colcemide for 2 h. Cells were then trypsinized, and the cell pellet was incubated in 0.56 % KCl for 20 min in 5 % CO2 at 37 °C (hypotonic shock). Cells were then fixed in methanol-acetic acid 3V/1V for 20 min at room temperature, then washed three times with methanol-acetic acid and concentrated in a small volume. Drops of cell suspension were then plated on glass slides at 50 °C. The cells were then allowed to dry and stained with Giemsa 4 % as described previously (Codner *et al*. 2016). We analysed 20 metaphase spreads.

### Comparative genomic hybridization

To confirm the increased copy number, we performed a comparative genomic hybridization (CGH) of the Ts65Dn and Ts66Yah models with their wild-type controls.

For Ts65Dn, these CGH data have been previously published from the original “1924” Ts65Dn mouse line (Duchon *et al*. 2011), however, they were reused in this study for comprehensive comparison with CGH data from Ts66Yah. For Ts65Dn, the CGH was undertaken using NimbleGen mouse HD2 oligonucleotide arrays. Comparative analysis was done using DNA extracts from one wild-type animal that were fluorescently labeled with Cy5 and from one animal bearing the duplication labeled with Cy3. After sonication and labeling, DNA was hybridized to the CGH array, followed by washing the slide according to the manufacturer’s instructions (Roche NimbleGen, Madison, WI, USA). Slides were scanned using a G2565 scanner at 3-lm resolution (Agilent Technologies, Palo Alto, CA, USA), and array images were analyzed using NimbleScan v2.5 software (Roche NimbleGen), with default parameters incorporating spatial correction. Arrays include 2,100,000 isothermal probes 50–75 bp in length with a median spacing of 1.1 kb throughout the genome.

For the Ts66Yah, labeling was performed using the SureTag DNA Labeling Kit (Agilent Technologies) from a microgram of genomic DNA (based on Qubit assays). The samples were digested for 2 hours at 37 ° C with restriction enzymes Alu I and Rsa I. After an inactivation step, the DNAs were denatured for 3 minutes at 98 ° C. The use of random primers and an exo-klenow fragment permit to mark samples with incorporation of dUTP coupled to cyanine 5 fluorochromes (Cy5) or cyanine 3 (Cy3). The targets thus synthesized were purified on 30kDa columns (Agilent Technologies). WT and trisomic samples were respectively labeled with Cy 3 and Cy5. Before carrying out the hybridization, the absorbances of the labeled DNAs were measured at 260 nm (DNA), 550 nm (cyanine 3) and at 650 nm (cyanine 5) with the spectrophotometer NanoDrop ND-1000. These values were used to assess performance and specific activity, the yield was be between 9 and 14 μg. After labeling, the DNAs were hybridized on CGH 4 × 180 K mouse slides (AMADID 027411, Agilent Technologies). Finally, the slides were scanned with G2505C scanner (Agilent Technologies).

### Sperm Analysis

Sperm analysis was done on 4 to 4.5 months old males with IVOS (Hamilton Thorne) apparatus. After euthanasia, the vasa deferentia and cauda epididymis were dissected and sperm was sampled. The quality and quantity of semen was estimated according to four main parameters: concentration (millions/ml), motility (%), rapid cells (%) and progressivity (%). For this analysis, sperm was diluted 20 times in prewarm (37°C) COOK solution (COOK ®, ref K-RVFE-50 COOK France). The suspension was gently agitated and placed for 3 to 4 minutes in a CO2 incubator before analysis with the IVOS system.

### Behavior analysis pipeline labo 1

We generated several experimental animal cohorts by selecting mice from litters containing a minimum of two male pups. One cohort was used for nesting activity, working memory in the Y maze, exploration of a novel environment in the open-field, spatial memory in the Morris water maze, and circadian activity. Three independent cohorts of animals were used for sperm, cranio-facial analysis and object recognition memory. Moreover, we also confirmed some phenotypes in another laboratory with a complete independent group of mice (17 males and 15 females 2n compared to 11 males and 14 females Ts66Yah). Independently of this line, a cohort of Ts65Dn mice was also built and assessed in the same behavioral pipeline, in order to permit direct comparison of both lines.

After weaning at 4 weeks of age, animals were sorted by litters into 39 × 20 × 16 cm cages (Green Line, Techniplast, Italy) where they had free access to purified water and food (D04 chow diet, Safe, Augy, France). The temperature was maintained at 23±1 °C and the light cycle was controlled as 12 h light and 12 h dark (lights on at 7 am). On testing days, animals were transferred to the antechambers of the experimental room 30 min before the start of the experiment. All the experiments were performed between 8:00 AM and 4:00 PM. A resting period of 2 days to 1 week was used between two consecutive tests.

A series of behavioral experiments were conducted on male mice with ages ranging from 1.8 months on starting to 4.5 months for the last test. The tests were administered in the following order: Circadian activity, nesting, Y-maze, square open field, open field, novel object recognition, Morris water maze (standard hidden and reversal) and fear conditioning (contextual and cue). Behavioral experimenters were blinded to the genetic status of the animals. All the tests were performed with the experimenter out of the animal’s sight. Further experiments for the evaluation of the spontaneous alternation and NOR made in a different laboratory (Fig. S5), where the two sexes were evaluated. The procedures for the test are detailed in the supplementary information.

Circadian activity (CA) was measured to assess spontaneous activity behavior over the complete light/dark cycle. The actimeter (Imétronic, Pessac, France) is composed of 8 individual boxes (11 × 21 × 18 cm3), each of them equipped with an array of four parallel horizontal infrared beams and linked to a computer allowing recordings of photocell beam breaks, providing automated measures of position and locomotor activity. Mice were put into cages at 11 am on the first day and removed the next day at 7 pm. The light cycle was controlled as 12 h light and 12 h dark (lights on at 7 am). The 32 hours of testing were divided into three different phases: the habituation phase (from 11 am to 7 pm on the first day); the night/dark phase (from 7 pm on the first day to 7 am on the second day); and the day/light phase (from 7 am to 7 pm on the second day).

Nesting test was performed by placing the mice individually in clean new housing cages two hours before the dark phase and the results were assessed the next morning. Normal bedding covered the floor to a depth of 0.5 cm. Each cage was supplied with a ‘Nestlet’, a 5 cm square of pressed cotton batting. The nests were assessed on a 5-point scale. 1-The Nestlet was largely untouched (>90% intact). 2-The Nestlet was partially torn up (50-90% remaining intact). 3-The Nestlet was mostly shredded but often there was no identifiable nest site: < 50% of the Nestlet remained intact but < 90% was within a quarter of the cage floor area, i.e. the cotton was not gathered into a nest but spread around the cage. 4-An identifiable but flat nest: > 90% of the Nestlet was torn up, the material was gathered into a nest within a quarter of the cage floor area, but the nest was flat with walls higher than mouse body height (curled up on its side) on less than 50% of its circumference. 5-A (near) perfect nest: > 90% of the Nestlet was torn up, the nest was a crater, with walls higher than mouse body height for more than 50% of its circumference (Deacon 2006).

Short-term memory was assessed by recording spontaneous alternation in the Y-maze test (Hughes 2004). The Y-maze test is based on the innate preference of animals to explore an arm that has not been explored previously, a behavior that, if occurring with a frequency greater than 50%, is called spontaneous alternation behavior (SAB). The maze was made of three enclosed plastic arms, each 40×9×16cm, set at an angle of 120° to each other in the shape of a Y. The wall of each arm had a different pattern to encourage SAB. Animals were placed at the end of one arm (this initial arm was alternated within the group of mice to prevent arm placement bias), facing away from the center, and allowed to freely explore the apparatus for 8 min under moderate lighting conditions (70 lux in the center-most region). The time sequences of entries in the 3 arms were recorded, (considering the mouse enters an arm when all four paws were inside the arm). Alternation was determined from successive entries into the three arms on overlapping triplet sets in which three different arms are entered. The number of alternations was then divided by the number of alternation opportunities, namely, total arm entries minus one. In addition, total entries were scored as an index of locomotor activity.

A square open-field (OF) was used to evaluate mice adaptation to a novel environment under stressful conditions. Mice were tested in an automated square OF (44.3 × 44.3 × 16.8 cm) made of PVC with transparent walls, a black floor, and covered with translucent PVC (Panlab, Barcelona, Spain). The open field arena was divided into central and peripheral regions and was homogeneously illuminated at 150 Lux. Each mouse was placed on the periphery of the open field and allowed to explore the apparatus freely for 30 min. The distance travelled, the number of rearing episodes, and the stereotypes in the central and peripheral zones were recorded over the 30 min test session. Stereotypes is the number of samples where the position of the subject is different from its position during the previous sample and equal to its position during the 2^nd^ sample back in time. The distance is the total distance (cm) travelled in the corresponding zone.

The novel object recognition task is based on the innate tendency of rodents to explore novel objects over familiar ones (Bevins and Besheer 2006). This test was done 24 hours after the last OF session performed in the same arena. On Day 1, mice were free to explore 2 identical objects for 10 min. After this acquisition phase, mice returned to their home cage for a 24-hour retention interval. Their memory was evaluated on day 2, using one familiar object (of those already experienced during the acquisition phase) and one novel object which were placed in the arena with the mice free to explore the two objects for a 10 min period. Between trials and subjects, the different objects were cleaned with 70° ethanol to reduce olfactory cues. To avoid a preference for one of the two objects, the new one was different and counterbalanced between the different animal groups and genotypes. Similarly, to avoid a location preference, the emplacement of the novel object compared to the familiar one (left or right) was counterbalanced too. Object exploration was manually scored and defined as the orientation of the nose to the object, allowing a distance < 1 cm. For the retention phase, the percentage of time exploring familiar vs novel objects was calculated to assess memory performance.

Spatial learning can be analyzed using the Morris Water Maze (MWM) task. This test was designed for the animals to learn to navigate a swimming tank (150 cm of diameter) filled with opaque water, following the most direct path to a hidden submerged platform when starting from different, random locations around the perimeter of the tank. Two principal axes of the maze were defined, each line bisecting the maze perpendicular to one another to create an imaginary ‘+’. The end of each line demarcates four cardinal points: North (N), South (S), East (E) and West (W) and four quadrants (NE, NW, SE, and SW). The use of distal cues provides the most effective strategy to accomplish this task and avoid the aversive effect of cold water (22°C). This ability is controlled by hippocampal-dependent spatial cognition. The partially trisomic mice were trained in the standard version of the water maze as previously described (Vorhees and Williams 2006; Arbogast *et al*. 2015). This standard version contains 2 different phases, a learning phase (using 7 acquisition sessions) and a probe test (to assess memory performance). Each acquisition session contained 4 trials in which mice were placed at one of the starting locations in random order (N, S, E, and W) and were allowed to swim until they located the platform situated in the target quadrant. Mice failing to find the platform within 60 s were gently guided and placed on it for 20 s (the same period of time as the successful animals). At the end of each learning and reversal phase, a probe test was done, with the platform removed and the time spent in the target and non-target quadrants as well as the number of platform annulus crossings during 60 s were recorded.

At the end of this standard version, memory flexibility was tested with a reversal phase, in which the PF was positioned in the opposite quadrant (using 4 acquisition sessions) and a probe test. Reversal learning in the MWM reveals whether or not animals can extinguish their initial learning and acquire a direct path to the new goal position. Finally, a cue session was done to validate the test and to determine the swimming speed and visual ability using the visible platform, clearly indicated by a visible cue (black flag). All the trials were recorded with a video tracking system (Ethovision, Wageningen, Netherlands). The swim paths for each mouse in each trial of the MWM test were categorized manually into one of the following search strategies. Thigmotaxis when mice have persistent swim along the wall of the pool. “Random” when mice swim over the entire area of the pool in straight swims. “Scanning” when the search path is restricted to the central area of the pool. “Chaining” when mice swim in a circular manner at a fixed distance from the wall. “Focal target” when swim search is restricted to the target quadrant. “Non-focal target” when mice search the PF in an incorrect quadrant. “Directly” when mice swim directly to the PF. Only “Focal search” and “directly” are considered as spatial search.

To further challenge hippocampus mediated cognitive behaviors we used the fear conditioning test. Fear conditioning (FC) is an associative learning paradigm for measuring aversive learning and memory where a neutral conditioned stimulus (CS) such as light and tone are paired with an aversive unconditioned stimulus (US) such as mild shock to the paw. Animals associate the spatial context cues with the CS concomitantly. After conditioning, the CS or the spatial context elicits a central state of fear in the absence of the US, translated into a reduced locomotor activity or total lack of movement (freezing) response (see supplementary information FC). Thus, immobility time is used as a measure of learning/memory performances (Crawley 1999; Goeldner *et al*. 2009).

Experiments were conducted in four operant chambers (28 × 21 × 22 cm) with a metal bar floor linked to a shocker (Coulbourn Instruments, Allentown US). Chambers were dimly lit with a permanent house-light and equipped with a speaker for tone delivery and infra-red activity monitor. The experimental procedure encompassed 3 sessions over 2 days where the activity/inactivity behaviour was monitored continuously and the duration of inactivity per 2 s was collected. In day 1, for the conditioning session, the mouse was allowed to acclimate for 4 min, then a light/tone (10 kHz, 80-dB) CS was presented for 20s and terminated by a mild shock in the paw, US (1sec, 0.4 mA). After the paw shock, animals were left in the chamber for another 2 minutes. We defined total freezing time in first 2min and 4min and 2min immediately after paw shock as PRE1, PRE2 and POST, respectively. In Day 2, the fear to context was tested by bringing back the mouse into the same chamber and allowing it to explore for 6 minutes without presentation of the light/auditory CS. The movements of the animal were monitored to detect freezing behaviour consequence of recognizing the chamber as the spatial context (contextual learning). The total freezing time was calculated per 2min time block as CONT2, CONT4 and CONT6. Finally, the cue testing was performed 5 hours after the context testing. Animals were tested in modified conditioning chambers with walls and floor of different colour and texture. The mouse could habituate for 2 minutes to the chamber and then it was subjected to light and auditory cues for 2 minutes to evaluate conditioning fear. The total freezing time was calculated by 2min block as PRECUE1, CUE1, PRECUE2 and CUE2.⍰

### Behavioral analysis labo 2

Three months-old Ts66Yah and 2n male and female mice were used in this study. The general health of mice was regularly checked throughout the experimental period. All experiments on animals were conducted in accordance with the ethical standards of French and European laws (European Communities Council Directive of 24 November 1986).

The spontaneous alternation behavior (SAB) was assessed in a Y maze (wall height 19.5 cm, arm length 26 cm, and arm width 6.3 cm). Each arm was covered with different cues in its walls (arm “A” with squares, arm “B” with lines, and “C” arm with triangles). Mice were introduced into the maze alternating the arm of the entry between mice. The 8-min test was video-recorded. An experimenter blind to the genotype analyzed the number of entries in each arm, and the number of complete alternations. The percentage of alternation was calculated as: (number of spontaneous alternations /(total number of arm entries −2)) × 100. The total mice used for this test were Ts66Yah-female = 12; Ts66Yah-male = 11; 2n-female = 17; 2n-male = 17.

The Nobel Object Recognition (NOR) task was conducted in a V-maze apparatus (adapted for Catuara-Solarz *et al*., 2016) with black walls (Y-maze adapted, wall height 19.5 cm, arm length 26 cm, and arm width 6.3 cm). Each day, mice were introduced into the maze placed in the center (between the two arms of the V-maze). The first day, mice were subjected to a 10-min habituation session during which they were allowed to explore the maze without any objects. The next day, mice went through a 10-min familiarization session in which two identical objects were situated at the end of each arm attached to the wall and the floor with adhesive tape. The next day, the recognition test session was conducted consisting of a 10-min trial in which a new object substituted one of the objects used at familiarization. The recognition of the new object was assessed by calculating the discrimination index (DI) by the following formula: DI = [(time exploring the novel object – time exploring the familiar object) /total exploration time)]. Two different pairs of objects were used. For each mouse, the type of object and the location of the novel object were randomized. The sessions were video-recorded and analyzed by an experimenter blind to the genotype. Mice exploring objects less than 3 sec in either of the phase were excluded from the analysis. When mice climbed on an object, the time the mice spent on the object as not counted as exploration in manual scoring. Between each trial the arena and objects were cleaned with Aniospray (Dutscher) to reduce olfactory cue. The total mice used for this test were Ts66Yah-female = 11; Ts66Yah-male = 11; 2n-female = 14; 2n-male = 13. For 2n vs Ts66Yah comparisons, we used a t-test.

### Statistical analysis

For each data set, we performed the Shapiro–Wilk test and Quantile-Quantile plots to analyze if the data were normally distributed and the Brown-Forsythe test to ascertain the homogeneity of variances. If the p-value was greater than the significance level (0.05), we assumed the normality and equal variance. In this case, statistical significance of differences between groups was inferred by a two-tailed T test between genotype or ANOVA for repeated measures (MWM, FC). The post hoc tests (Tukey Test) were conducted only if F in RMANOVA achieved a 0.05 level. In the case of datasets where the assumptions of normality or homogeneity of variances were not fulfilled, we did the Kruskal-Wallis non-parametric test. We performed a one sample two tail t-test respectively for the discrimination index (DI) of the NOR versus no discrimination (0%), or for the percentage of spontaneous alternation versus 50% (hazard) in the Y maze, or 25% (hazard) for the probe test in the MWM. One Ts66Yah was excluded from Y maze experiment because the number of visited arms was less than 5 and one wt animal from Ts65Dn group in MWM PT reversal was excluded because he showed a floating behaviour for more than 90% of time. The figures and the statistical analysis for labo 2 were done using GraphPad Prism 9. The phenotypes were compared between genotypes in male and female mice separately. The comparisons between 2n and Ts66Yah mice were made using a two tailed t-test. The percentage of alternation and the discrimination index (NOR) was analyzed using a two tailed one-sample t-test.

### Identify further the explanatory phenotyping variables in the trisomic lines

We used “Gdaphen” for “Genotype discrimination using phenotypic features”. Gaphen is a R pipeline that allows the identification of the most important predictive qualitative and quantitative variables for genotype discrimination in phenotypic based datasets without any prior hypothesis (available on github https://github.com/YaH44/GDAPHEN). More detailed are available in the supplementary data and in Muniz Moreno *et al* (ref).

Gdaphen is a R pipeline that allows the identification of the most important predictor qualitative and quantitative variables for genotype discrimination in animal models of different diseases. We used Gdaphen an unpublished R package developed in our lab (https://github.com/YaH44/GDAPHEN/releases/tag/Public) to identify the phenotypic explanatory variables of those recorded during the analysis more relevant to discriminate between genotypes mutant and wild-type from the Ts66Yah and Ts65Dn models. Moreover, we also identified which recorded variables were more relevant to discriminate between each specific model wild-types or mutants to more deeply understand the differences and similarities in the relevance of the alterations in the phenotypic characterization performed for those mice models.

Gdaphen takes as input data an excel table containing on the rows the info per animal and on each column all the variables recorded. As for some tests several variables were recorded, we grouped those variables with the same group label and identified the importance for the discrimination of i) each variable alone, ii) the overall contribution of the group.

Pre-processing steps were carried out to get the data into shape for the analysis, after which Gdaphen was able to first identify the highly correlated variables (more than a r=0.75) and remove them for downstream analysis. For example, in the sperm tests several variables measured for the test were highly correlated with each other. The sperm percentage of rapid cells and the percentage of cell motility showed a coefficient of correlation r=0.987. Similarly, the percentage of rapid spermatozoa (“Sperm:Rapid_cells “) was 0.97 correlated with the “percentage of progressive cells (“Sperm:Progressive”). Thus, we decided to keep only the two variables: the sperm concentration (“Sperm:C°”, millions/ml)” and the percentage of progressive cells(“Sperm:Progressive”).

We decided to use two different classifiers to answer two different questions: I) a GLM or GLM-Net model that will allow us to identify which phenotypic variables or “predicting variables” are able to discriminate due to the fact that their linear combination influences the value of the dependent variable response; II) a random forest, noted RF, unsupervised algorithm that will be able to identify relevant phenotypic variables for the discrimination even though they may not originate from a linear distribution or exponential distribution family or have a linear relationship. Both functions are taken from the caret R and nnet R packages.

This method is able to deal with groups of both qualitative and quantitative variables recorded from the same individuals. The MFA performs a normalization or “weighting” on each group by dividing all the variables belonging to the group by the first eigenvalue from the principal component analysis (PCA) of the group. Then a PCA on all the weighted variables is applied so we can identify the correlation between the grouped or ungrouped qualitative or quantitative variables, the principal component dimensions, and identify the individual coordinates of each observation on the PCA dimensions. The method is implemented using the MFAmix function from the PCAmixdata R package. Moreover, we chose a vectorization visualization approach like that implemented in PCAmixdata where we included the cosine similarity distance to further highlight the parameters that follow the same trajectory as Genotype. Consequently, these parameters contribute to the separation of the individual data on the same dimensions defined by their cosine similarity distance. We analyzed three different numbers of phenotypic predictor variables: i) all phenotypic variables; ii) the phenotypic variables left after removing the highly correlated ones (correlation higher than 75%); and iii) the phenotypic variables contributing more than a 30% in the discrimination after running the MFA analysis using all the variables and observing the correlation between the quantitative ungrouped phenotypic variables with the main three dimensions of the PCA. Our reasoning is to try to decrease the noise added by variables that do not strongly contribute to the discrimination, decrease the complexity of the model and the calculations, and increase the power on the discrimination since a lower number of variables are considered. Then we calculated the variance of the data that we were able to explain using the first 10 dimensions and the accuracy of the models to answer how well they can correctly predict each individual observation in the class of the dependent variable. We ran the Gdaphen pipeline to perform the genotype discrimination analyses on: 1) Ts65Dn mutants and controls; 2) Ts66Yah mutants and their respective controls; 3) Ts65Dn and Ts66Yah mutants and their respective control phenotypic data. In all these analyses, the model built using the phenotypic predictor variables known to contribute more than 30% to the discrimination was always able to explain a higher percentage of the variance in the data.

### Magnetic Resonance Imaging

Males from a dedicated cohort, at the age between 3 to 4.5 months, were anesthetized and perfused with 30 ml of room temperature 1X Phosphate Buffer Saline (PBS) supplemented with 10% (% w/v) heparine and 2mM of ProHance Gadoteridol (Bracco Imaging, Courcouronnes, France) followed by 30ml of 4% PFA supplemented with 2mM of ProHance Gadoteridol. Then the brain structure was dissected and kept in PFA 4% 2mM ProHance overnight at 4°C. The next day, each specimen was transferred into 1X PBS 2mM ProHance until imaging. Just prior to imaging, the brains were removed from the fixative and placed in plastic tubes (internal diameter 1 cm, volume 13 mL) filled with a proton-free susceptibility-matching fluid (Fluorinert® FC-770, Sigma-Aldrich, St. Louis, MO). Images of excised brains were acquired on a 7T BioSpec animal MRI system (Bruker Biospin MRI GmbH, Ettlingen, Germany), with an actively decoupled quadrature-mode mouse brain surface coil for signal reception and a 86-mm birdcage coil for transmission, both supplied by Bruker. Two imaging protocols were used. The first protocol consisted of a 3D T2-weighted rapid-acquisition with relaxation enhancement (RARE). The parameters for this sequence were: TR 325 ms, TE 32 ms, Rare factor = 6, interecho spacing 10.667 ms, 92 kHz bandwidth. The second imaging protocol consisted of a 3D T2*-weighted Fast Low Angle (FLASH) sequence with the following parameters: TR 50 ms, TE 25 ms, FA 50 degrees, 28 kHz bandwidth. The output image matrixes for both sequences were 195 × 140 × 90 over a field of view 19.5 × 14.0 × 9.0 mm3 yielding an isotropic resolution of 100 μm and were reconstructed using ParaVision 6.0.1. Each MRI image was segmented into twenty anatomical structures according to a multi-atlas label propagation framework. To this end, the ten manually segmented in-vitro MR images from the MRM NeAt Mouse Brain Database <u>(ht</u>t<u>p://brainatlas.mbi.ufl.edu/)</u> were considered (Ma *et al*. 2005). The image processing pipeline consisted of the following steps: (i) a skull-stripping step was first performed using the tissue brain segmentation method provided in SPMMouse (http://www.spmmouse.org) (Sawiak *et al*. 2009); (ii) each MR image was then corrected for bias field in homogeneity using N4ITK (Tustison *et al*. 2010); (iii) the ten anatomically annotated images from the MRM NeAt Mouse Brain Database were registered in a deformable way on each mouse image using the ANTs registration toolbox (http://stnava.github.io/ANTs/) (Avants *et al*. 2008); (iv) anatomical labels were finally fused using the simultaneous truth and performance level estimation (STAPLE) (Warfield *et al*. 2004). By this way, the volumes of the twenty anatomical structures as well as the whole brain were computed for each image modality and each mouse. Finally, the volumes computed from the two images modalities were averaged out to obtain the final volume associated to each mouse. Morphological MRI images were compared across groups with a region-based analysis. The resulting region-based volume estimations were averaged out for each animal before the statistical analysis (see supplementary information MRI). For Ts65Dn, these data have been previously published by our group (Duchon *et al*. 2021), were reanalysed with the same analysis pipeline for comprehensive comparison of both models.

### Morphometrics

Mice used for the craniofacial study were sacrificed at 18 weeks of age and carcasses were skinned, eviscerated and store in Ethanol 96%. Cranium morphology was assessed using the Quantum μCT scanner (Perkin Elmer, Whaltham, USA). All scans were performed with an isotropic voxel size of 20 μm, 160 μA tube current and 90 kV tube voltage. We applied a common approach to shape analysis named geometric morphometrics GM using the Geomorph software package in the R statistical computing environment (Adams and OtÁrola-castillo 2013). This approach used the coordinates of 39 relevant cranial landmarks that were recorded using Landmark software (Institute for Data Analysis and Visualization (IDAV) group at the University of California, Davis, Table S5). A generalized Procrustes analysis was then used to superimpose the specimens on a common coordinate system by holding their position, size and orientation constant. From the Procrustes-aligned coordinates, a set of shape variables was obtained which can be used in multivariate statistical analyses. Graphical methods were used to visualize patterns of shape variation. Taking a different approach, the principal component analysis (CPA) is a mathematical procedure that transforms a number of correlated variables into a number of uncorrelated variables. This permitted visualizing patterns of shape variation in shape space.

### QRT-PCR

cDNA synthesis was performed using the SuperScript® VILO™ cDNA Synthesis Kit (Invitrogen, Carlsbad, CA). PCRs were performed with TaqMan® Universal Master Mix II and pre-optimized TaqMan® Gene Expression assays (Applied Biosystems, Waltham, Massachusetts, USA), consisting of a pair of unlabeled PCR primers and a TaqMan® probe with an Applied Biosystems™ FAM™ dye label on the 5’ end and minor groove binder (MGB) and nonfluorescent quencher (NFQ) on the 3’ end (listed in Supplementary Table 4). mRNA expression profiles were analyzed by real-time quantitative PCR using TaqMan TM universal master mix II with UNG in a realplex II master cycler, Eppendorf (Hamburg, Germany). The complete reactions were subjected to the following program of thermal cycling: 1 cycle of 2 minutes at 50°C; 1 cycle of 10 minutes at 95°C; 40 cycles of 15 seconds at 95°C and 1 minute at 60°C. The efficiencies of the TaqMan assays were checked using cDNA dilution series from extracts of hippocampal sample. Normalization was performed by amplifying 4 housekeeping genes (*Gnas, Pgk1, Actb* and Atp5b) in parallel and using the GeNorm procedure to correct the variations of the amount of source RNA in the starting material (39). All the samples were tested in triplicate.

### Gene expression analyses

The hippocampus and entorhinal cortex from males Ts65Dn (n=6) and control littermates (n=6), and Ts66Yah (n=6) and control littermates (n=5), at the age of 5-6 months old, were isolated and flash frozen in liquid nitrogen. Total RNA was prepared using an RNA extraction kit (Qiagen, Venlo, Netherlands) according to the manufacturer’s instructions. Sample quality was checked using an Agilent 2100 Bioanalyzer (Agilent Technologies, Santa Clara, California, USA). All the procedures and the analysis are detailed in the supplementary information.

The preparation of the libraries was done by the GenomEast platform, a member of the ‘France Génomique’ consortium (ANR-10-INBS-0009), using the TruSeq Stranded Total RNA Sample Preparation Guide - PN 15031048. Total RNA-Seq libraries were generated from a minimum of 150-300 ng of total RNA using TruSeq Stranded Total RNA LT Sample Prep Kit with Ribo-Zero Gold (Illumina, San Diego, CA), according to the manufacturer’s instructions. The molecules extracted from the biological material were polyA RNA. Whole genome expression sequencing was performed by the platform using Illumina Hiseq 4000 and generating single end RNA-Seq reads of 50 bps length. The raw sequenced reads were aligned by Hisat2 against the GRCm38.v99 mouse assembly. 55385 ENSEMBL Gene Ids were quantified aligning with the GRCm38.v99 assembly. HTSeq-count was used to generate the raw counts. The downstream analyses were carried on with in-house bash scripts and R version 3.6 scripts using FCROS (Dembele et al., 2014) and DESeq2 (Love et al., 2014) packages to identify the DEGs. Raw reads and normalized counts have been deposited in GEO (GSE213500 for Ts65Dn and GSE213502 for Ts66Yah).

We performed the functional differential analysis using the GAGE pathway analysis (Luo *et al*. 2009) and grouped all the pathways into 25 functional categories (noted meta-pathways). Then, to assess gene connectivity we built a minimum fully connected protein-protein interaction (PPI) network (noted MinPPINet) of genes known to be involved in the synaptic function as they were associated with synaptic pathways via the GO (Ashburner *et al*. 2000) and KEGG databases (Esling *et al*. 2015). Regulatory information was also added to build the final RegPPINet. We used the betweenness centrality analysis to identify hubs, keys for maintaining the network communication flow.

To further study the genotype-phenotype relationship in those models we combined the behavioral results and the RNA-Seq data to identify central genes altered in the models and linked to the phenotypes observed using the genotype-phenotype databases GO, KEGG and DisGeNET.

First, we downloaded the list of experimentally validated genes known to be involved in hyperactivity or locomotion behavior from the human disease database DIsGeNET (dataset: Hyperactive behavior, C0424295 with 1263 genes) and annotated the genes with a high confidence ortholog in mouse. We added all the mouse genes involved in GO genesets linked to locomotion or motor behavior (18 GO terms: GO:0007626, GO:0008344, GO:0031987, GO:0033058, GO:0035641, GO:0040011, GO:0040012, GO:0040013, GO:0040017, GO:0043056, GO:0045475, GO:0090325, GO:0090326, GO:0090327, GO:1904059, GO:1904060, GO:0036343, GO:0061744). Then, we queried our RNA-Seq data for these genes to identify those found deregulated in the datasets.

## DATA availability

All the data from the Behavioural studies and the MRI are available at the Zenodo repository: 10.5281/zenodo.7067252; where the RNASeq transcriptomes for both lines have been deposited in GEO (GEO (GSE213500 for Ts65Dn and GSE213502 for Ts66Yah)).

## ACKNOWLEDGMENTS

We would like to thank the members of the research group, of the IGBMC laboratory and of the ICS for their help in brain morphometric analysis. We extend our thanks to the animal caretakers of the ICS who are in charge of mice well-being. This work was supported by the Jérôme Lejeune foundation. We acknowledge the Sisley-d’Ornano and Jérôme Lejeune Foundations that awarded MF with a postdoctoral fellowship.

This work was supported by the National Centre for Scientific Research (CNRS), the French National Institute of Health and Medical Research (INSERM), the University of Strasbourg (Unistra), French government funds through the “Agence Nationale de la Recherche” in the framework of the Investissements d’Avenir program by IdEx Unistra (ANR-10-IDEX-0002), a SFRI-STRAT’US project (ANR 20-SFRI-0012) and EUR IMCBio (ANR-17-EURE-0023) and INBS PHENOMIN (ANR-10-IDEX-0002-02), and also provided by the Joint Programming Initiative Neurodegenerative Diseases (JPND) multinational research project HEROES (ANR-17-JPCD-0003) et DYRK-DOWN (ANR-18-CE16-0020). This project received funding from the European Union’s Horizon 2020 research and innovation program under grant agreement GO-DS21 No 848077. The funders had no role in the study design, data collection and analysis, decision to publish, or preparation of the manuscript.

## REFERENCES

Ashburner, M., C. A. Ball, J. A. Blake, D. Botstein, H. Butler et al., 2000 Gene ontology: tool for the unification of biology. The Gene Ontology Consortium. Nat Genet 25: 25–29.

Aziz, N. M., F. Guedj, J. L. A. Pennings, J. L. Olmos-Serrano, A. Siegel et al., 2018 Lifespan analysis of brain development, gene expression and behavioral phenotypes in the Ts1Cje, Ts65Dn and Dp(16)1/Yey mouse models of Down syndrome. Dis Model Mech 11.

Aït Yahya-Graison, E., J. Aubert, L. Dauphinot, I. Rivals, M. Prieur et al., 2007 Classification of human chromosome 21 gene-expression variations in Down syndrome: impact on disease phenotypes. Am J Hum Genet 81: 475–491.

Codner, G. F., L. Lindner, A. Caulder, M. Wattenhofer-Donze, A. Radage et al., 2016 Aneuploidy screening of embryonic stem cell clones by metaphase karyotyping and droplet digital polymerase chain reaction. BMC Cell Biol 17: 30.

Cohen, S. J., and R. W. Stackman, 2015 Assessing rodent hippocampal involvement in the novel object recognition task. A review. Behav Brain Res 285: 105–117.

Concordet, J. P., and M. Haeussler, 2018 CRISPOR: intuitive guide selection for CRISPR/Cas9 genome editing experiments and screens. Nucleic Acids Res 46: W242–W245.

Davisson, M. T., C. Schmidt and E. C. Akeson, 1990 Segmental trisomy of murine chromosome 16: a new model system for studying Down syndrome. Prog Clin Biol Res 360: 263–280.

Davisson, M. T., C. Schmidt, R. H. Reeves, N. G. Irving, E. C. Akeson et al., 1993 Segmental trisomy as a mouse model for Down syndrome. Prog Clin Biol Res 384: 117–133.

Duchon, A., M. Del Mar Muñiz Moreno, S. M. Lorenzo, M. P. S. de Souza, C. Chevalier et al., 2021 Multi-influential genetic interactions alter behaviour and cognition through six main biological cascades in Down syndrome mouse models. Hum Mol Genet.

Duchon, A., and Y. Herault, 2016 DYRK1A, a Dosage-Sensitive Gene Involved in Neurodevelopmental Disorders, Is a Target for Drug Development in Down Syndrome. Front Behav Neurosci 10: 104.

Duchon, A., M. Raveau, C. Chevalier, V. Nalesso, A. J. Sharp et al., 2011 Identification of the translocation breakpoints in the Ts65Dn and Ts1Cje mouse lines: relevance for modeling Down syndrome. Mamm Genome 22: 674–684.

Esling, P., F. Lejzerowicz and J. Pawlowski, 2015 Accurate multiplexing and filtering for high-throughput amplicon-sequencing. Nucleic Acids Res 43: 2513–2524.

Faundez, V., I. De Toma, B. Bardoni, R. Bartesaghi, D. Nizetic et al., 2018 Translating molecular advances in Down syndrome and Fragile X syndrome into therapies. Eur Neuropsychopharmacol 28: 675–690.

Fernandez, F., W. Morishita, E. Zuniga, J. Nguyen, M. Blank et al., 2007 Pharmacotherapy for cognitive impairment in a mouse model of Down syndrome. Nat Neurosci 10: 411–413.

Firth, H. V., S. M. Richards, A. P. Bevan, S. Clayton, M. Corpas et al., 2009 DECIPHER: Database of Chromosomal Imbalance and Phenotype in Humans Using Ensembl Resources. Am J Hum Genet 84: 524–533.

García-Cerro, S., N. Rueda, V. Vidal, S. Lantigua and C. Martínez-Cué, 2017 Normalizing the gene dosage of Dyrk1A in a mouse model of Down syndrome rescues several Alzheimer’s disease phenotypes. Neurobiol Dis 106: 76–88.

Goodliffe, J. W., J. L. Olmos-Serrano, N. M. Aziz, J. L. Pennings, F. Guedj et al., 2016 Absence of Prenatal Forebrain Defects in the Dp(16)1Yey/+ Mouse Model of Down Syndrome. J Neurosci 36: 2926–2944.

Guedj, F., J. L. Pennings, L. J. Massingham, H. C. Wick, A. E. Siegel et al., 2016 An Integrated Human/Murine Transcriptome and Pathway Approach To Identify Prenatal Treatments For Down Syndrome. Sci Rep 6: 32353.

Hart, A. W., L. McKie, J. E. Morgan, P. Gautier, K. West et al., 2005 Genotype-phenotype correlation of mouse pde6b mutations. Invest Ophthalmol Vis Sci 46: 3443–3450.

Hatano, R., A. Takeda, Y. Abe, K. Kawaguchi, I. Kazama et al., 2018 Loss of ezrin expression reduced the susceptibility to the glomerular injury in mice. Sci Rep 8: 4512.

Heller, H. C., A. Salehi, B. Chuluun, D. Das, B. Lin et al., 2014 Nest building is impaired in the Ts65Dn mouse model of Down syndrome and rescued by blocking 5HT2a receptors. Neurobiol Learn Mem 116: 162–171.

Herault, Y., J. M. Delabar, E. M. C. Fisher, V. L. J. Tybulewicz, E. Yu et al., 2017 Rodent models in Down syndrome research: impact and future opportunities. Dis Model Mech 10: 1165–1186.

Hoelter, S. M., C. Dalke, M. Kallnik, L. Becker, M. Horsch et al., 2008 “Sighted C3H” mice - a tool for analysing the influence of vision on mouse behaviour? Frontiers in Bioscience 13: 5810–5823.

Itohara, S., Y. Kobayashi and T. Nakashiba, 2015 Genetic factors underlying attention and impulsivity: mouse models of attention-deficit/hyperactivity disorder, pp. 46–51. Current Opinion in Behavioral Sciences.

Kazuki, Y., F. J. Gao, Y. Li, A. J. Moyer, B. Devenney et al., 2020 A non-mosaic transchromosomic mouse model of down syndrome carrying the long arm of human chromosome 21. Elife 9.

Leal, S. L., and M. A. Yassa, 2015 Neurocognitive Aging and the Hippocampus across Species. Trends Neurosci 38: 800–812.

Ling, K. H., C. A. Hewitt, K. L. Tan, P. S. Cheah, S. Vidyadaran et al., 2014 Functional transcriptome analysis of the postnatal brain of the Ts1Cje mouse model for Down syndrome reveals global disruption of interferon-related molecular networks. BMC Genomics 15: 624.

Lorenzi, H., N. Duvall, S. Cherry, R. Reeves and R. Roper, 2010 PCR prescreen for genotyping the Ts65Dn mouse model of Down syndrome. Biotechniques 48: 35–38.

Luo, W., M. S. Friedman, K. Shedden, K. D. Hankenson and P. J. Woolf, 2009 GAGE: generally applicable gene set enrichment for pathway analysis. BMC Bioinformatics 10: 161.

Marechal, D., V. Brault, A. Leon, D. Martin, P. L. Pereira et al., 2019 Cbs overdosage is necessary and sufficient to induce cognitive phenotypes in mouse models of Down syndrome and interacts genetically with Dyrk1a. Hum Mol Genet.

Martinez-Cue, C., C. Baamonde, M. Lumbreras, J. Paz, M. T. Davisson et al., 2002 Differential effects of environmental enrichment on behavior and learning of male and female Ts65Dn mice, a model for Down syndrome. Behav Brain Res 134: 185–200.

Matsumoto, S., S. Fujii, A. Sato, S. Ibuka, Y. Kagawa et al., 2014 A combination of Wnt and growth factor signaling induces Arl4c expression to form epithelial tubular structures. EMBO J 33: 702–718.

Moore, C. S., C. Hawkins, A. Franca, A. Lawler, B. Devenney et al., 2010 Increased male reproductive success in Ts65Dn “Down syndrome” mice. Mamm Genome 21: 543–549.

Muñiz Moreno, M. D. M., V. Brault, M. C. Birling, G. Pavlovic and Y. Herault, 2020 Modeling Down syndrome in animals from the early stage to the 4.0 models and next. Prog Brain Res 251: 91–143.

Nagamani, S. C., A. Erez, C. Eng, Z. Ou, C. Chinault et al., 2009 Interstitial deletion of 6q25.2-q25.3: a novel microdeletion syndrome associated with microcephaly, developmental delay, dysmorphic features and hearing loss. Eur J Hum Genet 17: 573–581.

Nakano-Kobayashi, A., T. Awaya, I. Kii, Y. Sumida, Y. Okuno et al., 2017 Prenatal neurogenesis induction therapy normalizes brain structure and function in Down syndrome mice. Proc Natl Acad Sci U S A 114: 10268–10273.

Neumann, F., S. Gourdain, C. Albac, A. D. Dekker, L. C. Bui et al., 2018 DYRK1A inhibition and cognitive rescue in a Down syndrome mouse model are induced by new fluoro-DANDY derivatives. Sci Rep 8: 2859.

Nguyen, T. L., A. Duchon, A. Manousopoulou, N. Loaëc, B. Villiers et al., 2018 Correction of cognitive deficits in mouse models of Down syndrome by a pharmacological inhibitor of DYRK1A. Dis Model Mech 11.

O’Doherty, A., S. Ruf, C. Mulligan, V. Hildreth, M. L. Errington et al., 2005 An aneuploid mouse strain carrying human chromosome 21 with Down syndrome phenotypes. Science 309: 2033–2037.

Olmos-Serrano, J. L., H. J. Kang, W. A. Tyler, J. C. Silbereis, F. Cheng et al., 2016 Down Syndrome Developmental Brain Transcriptome Reveals Defective Oligodendrocyte Differentiation and Myelination. Neuron 89: 1208–1222.

Pizzo, L., M. Jensen, A. Polyak, J. A. Rosenfeld, K. Mannik et al., 2019 Rare variants in the genetic background modulate cognitive and developmental phenotypes in individuals carrying disease-associated variants. Genetics in Medicine 21: 816–825.

Reeves, R. H., N. G. Irving, T. H. Moran, A. Wohn, C. Kitt et al., 1995 A mouse model for Down syndrome exhibits learning and behaviour deficits. Nat Genet 11: 177–184.

Richtsmeier, J., L. Baxter and R. Reeves, 2000 Parallels of craniofacial maldevelopment in Down syndrome and Ts65Dn mice. Dev Dyn 217: 137–145.

Ruby, N., F. Fernandez, P. Zhang, J. Klima, H. Heller et al., 2010 Circadian locomotor rhythms are normal in Ts65Dn “Down syndrome” mice and unaffected by pentylenetetrazole. J Biol Rhythms 25: 63–66.

Schultz, H., T. Sommer and J. Peters, 2015 The Role of the Human Entorhinal Cortex in a Representational Account of Memory. Front Hum Neurosci 9: 628.

Shaw, P. R., J. A. Klein, N. M. Aziz and T. F. Haydar, 2020 Longitudinal neuroanatomical and behavioral analyses show phenotypic drift and variability in the Ts65Dn mouse model of Down syndrome. Dis Model Mech 13.

Sierra, C., I. De Toma, L. L. Cascio, E. Vegas and M. Dierssen, 2021 Social Factors Influence Behavior in the Novel Object Recognition Task in a Mouse Model of Down Syndrome. Front Behav Neurosci 15: 772734.

Starbuck, J. M., T. Dutka, T. S. Ratliff, R. H. Reeves and J. T. Richtsmeier, 2014 Overlapping trisomies for human chromosome 21 orthologs produce similar effects on skull and brain morphology of Dp(16)1Yey and Ts65Dn mice. Am J Med Genet A 164A: 1981–1990.

Yonemura, S., T. Matsui and S. Tsukita, 2002 Rho-dependent and -independent activation mechanisms of ezrin/radixin/moesin proteins: an essential role for polyphosphoinositides in vivo. J Cell Sci 115: 2569–2580.

