## Supplementary information for "Ts66Yah, an upgraded Ts65Dn mouse model for Down syndrome, for only the region homologous to Human chromosome 21"

#### Supplementary Tables

| **First environment** | **Ts66Yah** | **Ts65Dn** |
| --- | --- | --- |
| **Sperm Analysis** |  |  |
| Number of individuals | 11 2n/11 Ts | 10 2n/10 Ts |
| Sexe | Male | Male |
| Nb of litters | 8 | 7 |
| Concentration(M/ml) | *t* (20) = 11.056 ; *p*<0.001 | *H*(1) = 14.286 ; *p*<0.001 |
| Motility % | *t* (20) = 7.287 ; *p*<0.001 | *t*(18) = 10.338 ; *p*<0.001 |
| Rapid cells % | *t* (20) = 7.323 ; *p*<0.001 | *t*(18) = 10.857 ; *p*<0.001 |
| Progressive % | *t* (20) = 3.964 ; *p*<0.001 | *t*(18) = 4.776 ; *p*<0.001 |
| **Nesting** |  |  |
| Sexe | Male | Male |
| Nb of litters | 15 | 7 |
| Number of individuals | 22 2n/25 Ts | 15 2n/12 Ts |
| Nesting score | *H*(1) = 12.006 ; *p*<0.001 | *H*(1) = 6.778 ; *p* = 0.012 |
| **Square Open Field** |  |  |
| Sexe | Male | Male |
| Nb of litters | 10 | 11 |
| Number of individuals | 12 2n/16 Ts | 16 2n/16 Ts |
| Distance total | *t* (26) = 0.713 ; *p* = 0.482 | *H*(1) = 6.763 ; *p* = 0.009 |
| Stereotypies | *H*(1) = 3.450; *p* = 0.063 | *t*(30) = 2.249 ; *p*= 0.032 |
| Velocity | *t* (26) = 0.708 ; *p*= 0.485 | *H*(1) = 6.767 ; *p* = 0.009 |
| Nb of rears | *H*(1) = 1.401 ; *p* = 0.236 | *t*(30) = 0.348 ; *p*= 0.731 |
| Nb of rears in center | *H*(1) = 4.180 ; *p* = 0.04 | *H*(1) = 4.63 ; *p* = 0.03 |
| Distance in peripheral zone | *t* (26) = 1.063 ; *p* = 0.297 | *H*(1) = 8.865 ; *p* = 0.003 |
| **Circadian Activity** |  |  |
| Sexe | Male | Male |
| Nb of litters | 10 | 8 |
| Number of individuals | 12 2n/16 Ts | 12 2n/12 ts |
| Total Habituation Locomotion | *H*(1) = 0.0344 ; *p* = 0.852 | *t*(22) = -1.072 ; *p* = 0.295 |
| Total Habituation Rear | *H*(1) = 1.346 ; *p* = 0.245 | *t*(22) = -2.309 ; *p* = 0.0307 |
| Total Light Locomotion | *t* (26) = 5.175 ; *p* = 0.022 | H(1) = 4.815 ; *p* = 0.028 |
| Total Light Rear | *H*(1) = 9.108 ; *p* = 0.002 | *H*(1) = 8.003 ; *p* = 0.005 |
| Total First Night Locomotion | *t* (26) = 0.036 ; *p* = 0.971 | *t*(22) = -2.761 ; *p* = 0.0114 |
| Total First Night Rear | *H*(1) =0.551 ; *p* = 0.457 | *H*(1) = 11.603 ; *p*<0.001 |
| Total second Night Locomotion | *t* (26) =-0.757 ; *p* = 0.455 | *t*(22) = -2.872 ; *p* = 0.0089 |
| Total second Night Rear | *H*(1) =0.422 ; *p* = 0.515 | *t*(22) = -3.783 ; *p* = 0.00102 |
| **Y maze** |  |  |
| Sexe | Male | Male |
| Nb of litters | 10 | 11 |
| Number of individuals | 12 2n/16 Ts | 16 2n/16 Ts |
| Arms visited within genotype | *t*(25) = -0.671 ; *p* = 0.508 | *H*(1) = 5.247 ; *p* = 0.022 |
| % spontaneous alternation within genotype | *t*(25) = 1.88 ; *p* = 0.827 | *t*(30) = 3.2 ; *p* = 0.003 |
| One sample t test spontaneous alternation vs 50% 2n | *t*(11) = 4.266 ; *p =* 0.00133 | *t*(15) = 5.371 ; *p*<0.001 |
| One sample t test spontaneous alternation vs 50% Ts | *t*(11) = 0.117 ; *p* = 0.117 | *t*(15) = 2.623 ; *p* = 0.019 |
| **NOR 24H** |  |  |
| Sexe | Male | Male |
| Nb of litters | 10 | 11 |
| Number of individuals | 13 2n/10 Ts | 16 2n/16 Ts |
| One sample t test %NO vs 50% 2n | *t*(12) = 2.744 ; *p =* 0.0178 | *t*(15) = 2.211 ; *p* = 0.043 |
| One sample t test %NO vs 50% Ts | *t*(9) = 1.056 ; *p =* 0.318 | *t*(15) = 1.155 ; *p* = 0.266 |
| Object within genotype in habituation | *t*(21) =-2.358 ; *p* = 0.0281 | *H*(1) = 5.460, *p* = 0.019 |
| **Morris Water Maze** |  |  |
| Sexe | Male | Male |
| Nb of litters | 16 | 16 |
| Number of individuals | 22 2n/22 Ts | 22 2n/18 Ts |
| **Hidden version** |  |  |
| ANOVA RM Latency within genotype | *F*(1,252) = 6.066, *p = 0.018* | *F*(1,210) = 10.841, *p* = 0.002 |
| ANOVA RM Velocity within genotype | *F*(1,252) = 2.595, *p = 0.115* | *F*(1,210) = 0.198, *p* = 0.659 |
| ANOVA RM thigmotaxis within genotype | *F*(1,252) = 1.011, *p = 0.321* | *F*(1,210) = 5.408, *p* = 0.028 |
| One sample t test %TQ vs 50% 2n | *t*(21) = 5.966; *p*<0.001 | *t*(21) = 3.085 ; *p* = 0.00561 |
| One sample t test %TQ vs 50% Tg | *t*(21) = 2.852; *p* = 0.009 | *t*(17) = 0.341 ; *p* = 0.737 |
| **Reversal** |  |  |
| ANOVA RM Latency within genotype | *F*(1,126) = 3.923, *p = 0.054* | *F*(1,105) = 16.361, *p*<0.001 |
| ANOVA RM Velocity within genotype | *F*(1,126) = 0.0253 *p = 0.874* | *F*(1,105) = 0.000143, *p*<0.991 |
| ANOVA RM thigmotaxis within genotype | *F*(1,126) = 0.358 *p = 0.553* | *F*(1,105) = 3.918, *p = 0.056* |
| One sample t test %TQ vs 50% 2n | *t*(21) = 2.871; *p*<0.009 | *t*(20) = 2.388; *p* = 0.00269 |
| One sample t test %TQ vs 50% Ts | *t*(21) = 2.796; *p*<0.011 | *t*(17) = 0.847 ; *p* = 0.409 |
| **Cue version** |  |  |
| ANOVA Latency within genotype | *H*(1) = 4.414, *p*= 0.036 | *H*(1) = 0.728, *p*= 0.393 |
| **Fear Conditioning** |  |  |
| Sexe | Male | Male |
| Nb of litters | 10 | 11 |
| Number of individuals | 12 2n/16 Ts | 16 2n /16 Ts |
| **FC Hab1** |  |  |
| Hab1 session | *H*(1) = 3.95 ; *p* = 0.047 | *H*(1) = 0.347 ; *p* = 0.852 |
| Hab2 session | *H*(1) = 4.078 ; *p* = 0.043 | *t*(22) = 3.264; *p*=0.00356 |
| **FC Context** |  |  |
| Cont1 | *t*(26) = 0.006 ; *p* = 0.995 | *H*(1) = 0.0128 ; *p* = 0.910 |
| Cont2 | *t*(26) = -1.74 ; *p* = 0.093 | *t(*30) = 1.056; *p* = 0.299 |
| Cont3 | *H*(1) = 0.026 ; *p* = 0.870 | *H*(1) = 4.031 ; *p* = 0.045 |
| **IRM analysis** |  |  |
| Sexe | Male | Male |
| Nb of litters | 5 | 3 |
| number of individuals | 6 2n/7 Ts | 5 2n/6 Ts |
| Amygdala | *t*(11) = 0.277; *p*= 0.786 | *t*(9) =-0.129; *p*=0.899 |
| Ant commissure | *t*(11) = -0.959; *p*= 0.357 | *H*(1) = 0.410; *p=* 0.521 |
| BasalForebrain Septum | *t*(11) = -0.05; *p*= 0.955 | *t*(9) = -1.145; *p*= 0.281 |
| Brain stem | *t*(11) = -0.785 ; *p*= 0.448 | *t*(9) = 1.325; *p*= 0.217 |
| Caudate putamen | *t*(11) = -0.193 ; *p*= 0.850 | *t*(9) = -0.463; *p*= 0.654 |
| Central gray | *H*(1) = 5.22 ; *p =* 0.022 | *t*(9) = -2.151; *p*= 0.059 |
| Cerrebelum | *t*(11) = -1.22; *p*= 0.245 | *t*(9) = 1.522; *p*= 0.162 |
| External capsule | *t*(11) = -1.058; *p*= 0.312 | *t*(9) = -0.643; *p*= 0.535 |
| Fimbria | *t*(11) = -2.81; *p*= 0.016 | *t*(9) = -2.18; *p*= 0.056 |
| Globus pallidus | *t*(11) = -1.40 ; *p*= 0.187 | *t*(9) = -1.021; *p*= 0.333 |
| Hippocampus | *t*(11) = -1.27 ; *p*= 0.227 | *t*(9) =- -0.757; *p*= 0.467 |
| Hypothalamus | *H*(1) = -1.313 ; *p =* *0.215* | *t*(9) = -0.633; *p*= 0.542 |
| Inferior colliculi | *t*(11) = -0.72 ; *p*= 0.484 | *t*(9) = -1.341; *p*= 0.212 |
| Internal Capsule | *t*(11) = -1.712 ; *p*= 0.114 | *t*(9) = -1.60; *p*= 0.142 |
| Rest of Midbrain | *H*(1) = 5.224; *p=* 0.022 | *t*(9) = -1.00 ; *p*= 0.341 |
| Superior colliculi | *H*(1) = 4.59; *p=* 0.032 | *t*(9) = -2.173; *p*= 0.057 |
| Thalamus | *t*(11) = -2.077 ; *p* = 0.062 | *t*(9) = -1.287; *p*= 0.230 |
| Ventricles | *t*(11) = -2.222; *p* = 0.048 | *t*(9) = -1.546; *p*= 0.156 |
| WholeBrain | *t*(11) = -0.531 ; *p* = 0.605 | *t*(9) = 0.089; *p*= 0.930 |
| **qPCR analysis** |  |  |
| Sexe | Male | Male |
| Nb of litters | 5 | 7 |
| number of individuals | 5 2n/6 Ts | 6 2n/5 Ts |
| *Dyrk1a* | *t*(9) = -6.033; *p* = 0.000236 | *t*(9) =- 4.0958; *p* = 0.00594 |
| *Pde10A* | *t*(9) = 0.316; *p* = 0.759 | *t*(9) = -2.729; *p* = 0.0238 |
| *Sod1* | *t*(9) = -4.892; *p* = 0.00086 | *t*(9) = -3.675; *p* = 0.00522 |
| *Stat4* | *t*(9) = -0.0297; *p* = 0.977 | *H*(1) = 0.033 ; *p = 0.855* |
| *Ccr6* | *t*(9) = 3.267; *p* = 0.029 | *t*(9) = -2.729; *p* = 0.0238 |
| *Ezr* | *H*(1) = 0.3 ; *p = 0.584* | *t*(9) = -4.058; *p* = 0.00627 |
| *Sh3bgr* | *t*(9) = -4.995; *p* = 0.00362 | *t*(9) = -2.786; *p* = 0.0295 |
| **Second environment** | **Ts66Yah** | **Ts65Dn** |
| **Y maze** |  | **Not done** |
| Sexe | Male |  |
| Nb of litters | 8 |  |
| number of individuals | 13 2n/11 Ts |  |
| Arms visited within genotype | *t*(22) = 1.141; *p* = 0.266 |  |
| % spontaneous alternation within genotype | *t*(22) = 4.436; *p* = 0.0002 |  |
| One sample t test spontaneous alternation vs 50% 2n | *t*(12) = 4.348; *p =* 0.0009 |  |
| One sample t test spontaneous alternationvs 50% Ts | *t*(10) = 1.814; *p* = 0.0997 |  |
| **Y maze** |  | **Not done** |
| Sexe | Female |  |
| Nb of litters | 8 |  |
| number of individuals | 14 2n/12 Ts |  |
| Arms visited within genotype | *t*(24)= 2.122; *p* = 0.044 |  |
| % spontaneous alternation within genotype | *t*(24) = 2.694; *p* = 0.0127 |  |
| One sample t test spontaneous alternation vs 50% 2n | *t*(13) = 3.235; *p =* 0.0065 |  |
| One sample t test spontaneous alternationvs 50% Ts | *t*(11) = 0.1530; *p* = 0.881 |  |
| ANOVA "nbr of entries" genotype/sexe for sexe | *F*_(1,46)_ = 0.01; *p* = 0.921 |  |
| ANOVA "Spontaneous alternation" genotype/sexe for sexe | *F*_(1,46)_ = 0.25; *p* = 0.619 |  |
| **NOR 24H** |  | **Not done** |
| Sexe | Male |  |
| Nb of litters | 8 |  |
| number of individuals | 13 2n/11 Ts |  |
| Object within genotype in habituation | *t*(22) = 2.091; *p* = 0.048 |  |
| One sample t test %NO vs 50% 2n | *t*(12) = 2.25; *p =* 0.044 |  |
| One sample t test %NO vs 50% Ts | *t*(10) = 2.017; *p* = 0.0713 |  |
| **NOR 24H** |  | **Not done** |
| Sexe | Female |  |
| Nb of litters | 8 |  |
| number of individuals | 14 2n/12 Ts |  |
| Object within genotype in habituation | *t*(24) = 3.676; *p* = 0.0012 |  |
| One sample t test %NO vs 50% 2n | *t*(13) = 4.088; *p* = 0.0013 |  |
| One sample t test %NO vs 50% Ts | *t*(11) = 0.9938; *p* = 0.341 |  |
| ANOVA "sniffing time presentation" genotype/sexe for sexe | *F*_(1,46)_ = 0.411; *p* = 0.525 |  |
| ANOVA "RI" genotype/sexe for sexe | *F*_(1,46)_ = 0.2009; *p* = 0.66 |  |

**Table S1**. **Detailed behaviour statistics results (statistically significant results are highlighted).**

| **Variable** | **Detailed name** | **unit (SI)** |
| --- | --- | --- |
| OF:Total_distance | total distance travelled in the open field test | m |
| OF:Rears | total number of rears in the open field |  |
| OF:Sterotypies | total number of stereotypies |  |
| OF:V | Mean velocity during the open field session | cm/s |
| CA:Locomotion:Hab | nb of back and forth during the habituation period of the circadian activity |  |
| CA:Rears:Hab | total number of rears during the circadian activity |  |
| CA:Locomotion:Light | nb of back and forth during the light period of the circadian activity |  |
| CA:Rears:Light | total number of rears during the light phase of the circadian activity |  |
| CA:Locomotion:N1 | nb of back and forth during the night phase 1 of the circadian activity |  |
| CA:Rears:N1 | total number of rears during the night phase 1 of the circadian activity |  |
| CA:Locomotion:N2 | nb of back and forth during the night phase 2 of the circadian activity |  |
| CA:Rears:N2 | total number of rears during the night phase 2 of the circadian activity |  |
| YM:Arm_Entries | total number of arm entries in the Y maze |  |
| YM:Spont_Alter | Percentage of spontaneous alternation in the Y maze |  |
| NOR:Pres_HG | time sniffing the left object in the presentation of the NOR | s |
| NOR:Pres_HD | time sniffing the right object in the presentation of the NOR | s |
| NOR:Test_FO | time sniffing the familiar object in the test phase of the NOR | s |
| NOR:Test_NO | time sniffing the novel object in the test phase of the NOR | s |
| NOR:Test_NO | Percentage of time spent exploring the novel object in the NOR |  |
| MWM:PT1_TQ | Percentage of time spent in the targeted quadrant of the probe test 1 |  |
| MWM:PTRev_TQ | Percentage of time spent in the targeted quadrant of the probe test for the reversal |  |
| FC:Hab1 | Freezing time during the habituation phase 1 of the fear conditioning | s |
| FC:Hab2 | Freezing time during the habituation phase 2 of the fear conditioning | s |
| FC:Cont1 | Freezing time during the contextual phase 1 of the fear conditioning | s |
| FC:Cont2 | Freezing time during the contextual phase 2 of the fear conditioning | s |
| FC:Cont3 | Freezing time during the contextual phase 3 of the fear conditioning | s |
| FC:Precue1 | Freezing time during the precued phase 1 of the fear conditioning |  |
| FC:Cue1 | Freezing time during the cued phase 1 of the fear conditioning |  |
| FC:Precue2 | Freezing time during the precued phase 2 of the fear conditioning |  |
| FC:Cue2 | Freezing time during the cued phase 2 of the fear conditioning |  |
| Sperm:C° | Sperm concentration | 10^6^/ml |
| Sperm:Motility | Percentage of motile spermatozoa |  |
| Sperm:Rapid_cells | Percentage of rapid spermatozoa |  |
| Sperm:Progressive | Percentage of progressive spermatozoa |  |
| Nesting:Score | Standardized core of nesting activities |  |

**Table S2.** **Phenotypic variables used for the Gdaphen analysis.**

| **Models Analysed** | **Variable 1** | **Variable 2** | **Correlation coefficient** |
| --- | --- | --- | --- |
| Ts66Yah vs wt | CA:Rears:N1 | CA:Locomotion:N1 | 0.808 |
| Ts66Yah vs wt | CA:Rears:Hab | CA:Locomotion:Hab | 0.796 |
| Ts66Yah vs wt | CA:Locomotion:Light | CA:Locomotion:N2 | 0.853 |
| Ts66Yah vs wt | Sperm:Motility | Sperm:Rapid_cells | 0.890 |
| Ts66Yah vs wt | Sperm:Motility | Sperm:Progressive | 0.779 |
| Ts66Yah vs wt | Sperm:Rapid_cells | Sperm:Motility | 0.890 |
| Ts66Yah vs wt | Sperm:Rapid_cells | Sperm:Progressive | 0.927 |
| Ts66Yah vs wt | FC:Cue2 | FC:Cue1 | 0.822 |
| Ts66Yah vs wt | FC:Cue2 | FC:Precue2 | 0.806 |
| Ts66Yah vs wt | CA:Locomotion:Hab | CA:Rears:Hab | 0.796 |
| Ts66Yah vs wt | CA:Locomotion:Hab | CA:Locomotion:N1 | 0.770 |
| Ts66Yah vs wt | CA:Rears:Light | CA:Rears:N2 | 0.967 |
| Ts66Yah vs wt | OF:V | OF:Total_distance | 1.000 |
| Ts66Yah vs wt | FC:Cue1 | FC:Precue2 | 0.811 |
| Ts66Yah vs wt | FC:Cue1 | FC:Cue2 | 0.822 |
| Ts66Yah vs wt | FC:Hab1 | FC:Hab2 | 0.801 |
| Ts65Dn vs wt | CA:Rears:N1 | CA:Rears:Hab | 0.800 |
| Ts65Dn vs wt | Sperm:Motility | Sperm:C° | 0.820 |
| Ts65Dn vs wt | Sperm:Motility | Sperm:Rapid_cells | 0.981 |
| Ts65Dn vs wt | Sperm:Motility | Sperm:Progressive | 0.865 |
| Ts65Dn vs wt | Sperm:Rapid_cells | Sperm:C° | 0.775 |
| Ts65Dn vs wt | Sperm:Rapid_cells | Sperm:Motility | 0.981 |
| Ts65Dn vs wt | Sperm:Rapid_cells | Sperm:Progressive | 0.863 |
| Ts65Dn vs wt | CA:Locomotion:N2 | CA:Locomotion:Hab | 0.782 |
| Ts65Dn vs wt | CA:Locomotion:N2 | CA:Locomotion:N1 | 0.782 |
| Ts65Dn vs wt | CA:Locomotion:Light | CA:Rears:Light | 0.925 |
| Ts65Dn vs wt | OF:Total_distance | OF:V | 1.000 |
| Ts65Dn vs wt | CA:Locomotion:Hab | CA:Locomotion:N1 | 1.000 |
| Ts65Dn vs wt | CA:Locomotion:Hab | CA:Locomotion:N2 | 0.782 |
| Ts65Dn vs wt | FC:Cue2 | FC::Cue1 | 0.767 |
| Ts65Dn, Ts66Yah vs wt | Sperm:Rapid_cells | Sperm:Motility | 0.987 |
| Ts65Dn, Ts66Yah vs wt | Sperm:Rapid_cells | Sperm:Progressive | 0.872 |
| Ts65Dn, Ts66Yah vs wt | Sperm:Motility | Sperm:Rapid_cells | 0.987 |
| Ts65Dn, Ts66Yah vs wt | Sperm:Motility | Sperm:Progressive | 0.849 |
| Ts65Dn, Ts66Yah vs wt | CA:Rears:N1 | CA:Locomotion:N1 | 0.763 |
| Ts65Dn, Ts66Yah vs wt | CA:Locomotion:N2 | CA:Locomotion:N1 | 0.777 |
| Ts65Dn, Ts66Yah vs wt | CA:Rears:Light (nb) | CA:Rears:N2 | 0.864 |
| Ts65Dn, Ts66Yah vs wt | CA:Locomotion:N1 | CA:Locomotion:Hab | 0.869 |
| Ts65Dn, Ts66Yah vs wt | CA:Locomotion:N1 | CA:Rears:N1 | 0.763 |
| Ts65Dn, Ts66Yah vs wt | CA:Locomotion:N1 | CA:Locomotion:N2 | 0.777 |
| Ts65Dn, Ts66Yah vs wt | OF:Total_distance | OF:V | 1.000 |
| Ts65Dn, Ts66Yah vs wt | FC:Cue2 | FC:Cue1 | 0.794 |

**Table S3. Highly correlated variables identified in the GDAPHEN analysis of Ts66Yah, Ts65Dn and the joint analysis of Ts65Dn and Ts66Yah behavioral data.**

|  | Ts65Dn | | Ts66Yah | |
| --- | --- | --- | --- | --- |
| Variable | 2n | DS | 2n | DS |
| GLMNET |  |  |  |  |
| Sperm:Progressive | 17.6 | 27.5 | 15.7 | 15.7 |
| FC:Hab2 | 10.1 | 19.1 | 18.0 | 10.1 |
| MWM:PTRev_TQ | 5.2 | 22.4 | 8.4 | 5.2 |
| NOR:Test_NO_% | 12.1 | 25.9 | 12.3 | 12.1 |
| NOR:Test_NO_T | 0.3 | 19.1 | 12.5 | 0.3 |
| YM:Spont_Alter | 8.1 | 17.6 | 7.1 | 7.1 |
| CA:Rears:N2 | 25.1 | 29.0 | 50.0 | 25.1 |
| CA:Locomotion:Light | 3.6 | 2.4 | 3.5 | 2.4 |
| CA:Locomotion:Hab | 0.0 | 0.0 | 11.2 | 23.7 |
| OF:Sterotypies | 22.1 | 21.7 | 21.7 | 100.0 |
| OF:Rears | 6.4 | 40.8 | 15.4 | 6.4 |
| Random Forest |  |  |  |  |
| Sperm:Progressive | 30.3 | 58.2 | 39.6 | 18.0 |
| FC:Hab2 | 50.0 | 13.9 | 13.4 | 19.5 |
| MWM:PTRev_TQ | 10.9 | 29.7 | 13.9 | 6.9 |
| NOR:Test_NO_% | 11.6 | 16.6 | 29.6 | 25.1 |
| NOR:Test_NO_T | 48.4 | 15.1 | 12.6 | 41.5 |
| YM:Spont_Alter | 34.0 | 31.1 | 15.5 | 21.9 |
| CA:Rears:N2 | 27.9 | 72.4 | 25.3 | 13.6 |
| CA:Locomotion:Light | 24.4 | 14.0 | 8.5 | 21.0 |
| CA:Locomotion:Hab | 21.5 | 33.3 | 1.5 | 6.6 |
| OF:Sterotypies | 75.7 | 47.3 | 0.0 | 100.0 |
| OF:Rears | 18.5 | 5.3 | 15.4 | 5.1 |

##### Table S4a. Discriminating variables identified from the GLM-Net and RF analysis combining the Ts66Yah &Ts65Dn data.

| **Ts66Yah vs Wt** | | | |
| --- | --- | --- | --- |
| **GLM-Net** | **Sel model >30%: 9 Variables** **Accuracy: 0.941** | **RF** | **Sel model >30%: 9 Variables** **Accuracy: 0.925** |
| Overall | Variable | Overall | Variable |
| 100 | Sperm:C° | 100 | Sperm:C° |
| 72.57 | Sperm:Progressive | 52.56 | Sperm:Progressive |
| 39.21 | MWM:PTRev_TQ | 30.58 | Nesting:Score |
| 33.28 | Nesting:Score | 29.14 | MWM:PT1_TQ |
| 25.62 | CA:Locomotion:N1 | 22.07 | OF:Total_distance |
| 5.86 | CA:Rears:N2 | 10.33 | CA:Rears:N2 |
| 5.35 | OF:Total_distance | 8.66 | FC:Cont3 (s) |
| 1.94 | FC:Cont3 | 3.64 | CA:Locomotion:N1 |
| 0 | MWM:PT1_TQ | 0 | MWM:: PTRev TQ |
| **Ts65Dn vs wt** | | | |
| **GLM-Net** | **Sel model >30%: 11 Variables** **Accuracy: 0.7** | **RF** | **Sel model >30%: 11 Variables** **Accuracy: 0.891** |
| 100 | FC:Precue1 | 100 | Sperm:C° |
| 57.02 | FC:Cont3 | 65.11 | FC:Cont3 |
| 52.43 | MWM:PT1_TQ | 42.15 | FC:Precue1 |
| 46.81 | YM:Arm_Entries | 36.11 | Nesting:Score |
| 30.66 | Nesting:Score | 29.68 | CA:Rears:Hab |
| 17.96 | Sperm:C° | 11.99 | YM:Arm_Entries |
| 16.82 | NOR:Test_NO | 7.39 | NOR:Pres_HD |
| 12.17 | FC:Hab1 | 7.09 | FC:Hab1 |
| 5.75 | OF:Rears | 3.13 | NOR:Test_NO |
| 5.75 | NOR:Pres_HD | 1.13 | OF:Rears |
| 0 | CA:Rears:Hab | 0 | MWM:PT1_TQ (%) |

**Table S4b. Comparison of the most discriminative variables identified between the genotypes for Ts66Yah vs wt, or for Ts65Dn vs wt, using the GLM-Net or RF methods.**

| Cranium | | | Description |
| --- | --- | --- | --- |
| Right | Median | Left |  |
|  | 1 |  | Nasale |
|  | 2 |  | Nasion |
|  | 3 |  | Bregma |
|  | 4 |  | Parietal-occipital junction |
|  | 5 |  | Midline of the interparietal-occipital junction |
|  | 6 |  | Dorsal midpoint of the foramen magnum |
| 7 |  | 15 | Dorsal-most point of the incisor alveoli |
| 8 |  | 16 | Parietal-premaxillar-maxillar junction |
| 9 |  | 17 | Anterior-most point of the zygomatic spine |
| 10 |  | 18 | Posterior-most point of the frontal-maxillary dorsal junction |
| 11 |  | 19 | Anterior-most point of the squamosal-parietal junction |
| 12 |  | 20 | Anterior-most point of the zygomatic-squamosal junction |
| 13 |  | 21 | Posterior-most point of the zygomatic-squamosal junction |
| 14 |  | 22 | Tip of the post-tympanic hook. |
| 23 |  | 31 | Anterior-most point of the anterior palatine foramen |
| 24 |  | 32 | Posterior-most point of the anterior palatine foramen |
| 25 |  | 33 | Ventral-most point of the premaxillar, maxillar and anterior palatine foramen junction |
| 26 |  | 34 | Mesial-most point of the first upper molar cervix |
| 27 |  | 35 | Point of greatest curvature of the posterior margin of malar process |
| 28 |  | 36 | Distal-most point of the third upper molar cervix |
| 29 |  | 37 | Point of greatest curvature of the squamosal retroversus process |
| 30 |  | 38 | Antero-medial projection of ectotympanic in basicranial |
|  | 39 |  | ventral midpoint of the foramen magnum |
| Mandibule |  |  |  |
| 1 |  | 12 | Tip of the coronoid process |
| 2 |  | 13 | Distal-most point of the third lower molar cervix |
| 3 |  | 14 | Mesial-most point of the first lower molar cervix |
| 4 |  | 15 | Dorsal-most point of the incisor alveoli |
| 5 |  | 16 | Inferior-most point of the incisor alveoli |
| 6 |  | 17 | Inferior-most point on border of ramus inferior to incisor alveolar |
| 7 |  | 18 | Superior-most point on inferior border of mandibular ramus |
| 8 |  | 19 | Tip of the mandibular angle |
| 9 |  | 20 | Ventral-most point of the mandibular condyle |
| 10 |  | 21 | Anterior-most point of the mandibular condyle |
| 11 |  | 22 | Mandibular foramen |

**Table S5. Landmarks used for craniofacial analysis of mouse models.**

|  | Ts65Dn EC | Ts65Dn HC | Ts66Yah EC | Ts66Yah HC |
| --- | --- | --- | --- | --- |
| Differentially Expressed Genes  (DEGs) identified by FCROS | 1691 | 1836 | 2220 | 1902 |
| Genes from the Mmu17 centromeric genes |  |  |  |  |
| Expressed genes | 53 | 53 | 53 | 53 |
| Upregulated Triplicated (TEGs) | 41 | 37 | 7 | 10 |
| Genes from the Mmu16 homologous region to Hsa21 |  |  |  |  |
| Expressed genes | 94 | 94 | 94 | 94 |
| Upregulated Triplicated (TEGs) | 81 | 85 | 82 | 84 |
| Number of GAGE KEGG and GOs (CC.BP.MF) terms dis-regulated in the trisomic models (FDR<0.1) | 135 | 493 | 40 | 323 |
| Number of GAGE KEGG and GOs upregulated in the trisomic models (FDR<0.1) | 95 | 459 | 1 | 260 |

**Table S6a:** **Summary of the differential expression analysis and differential functional analysis for each DS model dataset**.

| **Chr** | **Chr length (pb)** | **DEGs Ts66Yah HIP** | **DEGs Ts66Yah EC** | **DEGs Ts65Ds HIP** | **DEGs Ts65Dn EC** | **%DEGs Ts66Yah HIP /10^6^bp** | **%DEGs Ts66Yah EC/10^6^bp** | **%DEGs Ts65Dn HIP 10^6^bp** | **%DEGs Ts65Dn EC/10^6^bp** |
| --- | --- | --- | --- | --- | --- | --- | --- | --- | --- |
| 1 | 195154279 | 142 | 138 | 132 | 117 | 0.73 | 0.71 | 0.68 | 0.60 |
| 2 | 181755017 | 152 | 183 | 110 | 116 | 0.84 | 1.01 | 0.61 | 0.64 |
| 3 | 159745316 | 123 | 136 | 90 | 103 | 0.77 | 0.85 | 0.56 | 0.64 |
| 4 | 156860686 | 77 | 106 | 97 | 94 | 0.49 | 0.68 | 0.62 | 0.60 |
| 5 | 151758149 | 118 | 117 | 124 | 105 | 0.78 | 0.77 | 0.82 | 0.69 |
| 6 | 149588044 | 115 | 127 | 91 | 87 | 0.77 | 0.85 | 0.61 | 0.58 |
| 7 | 144995196 | 140 | 161 | 162 | 139 | 0.97 | 1.11 | 1.12 | 0.96 |
| 8 | 130127694 | 91 | 111 | 80 | 64 | 0.70 | 0.85 | 0.61 | 0.49 |
| 9 | 124359700 | 98 | 109 | 96 | 88 | 0.79 | 0.88 | 0.77 | 0.71 |
| 10 | 130530862 | 84 | 104 | 80 | 78 | 0.64 | 0.80 | 0.61 | 0.60 |
| 11 | 121973369 | 96 | 133 | 120 | 92 | 0.79 | 1.09 | 0.98 | 0.75 |
| 12 | 120092757 | 73 | 94 | 64 | 75 | 0.61 | 0.78 | 0.53 | 0.62 |
| 13 | 120883175 | 75 | 83 | 83 | 83 | 0.62 | 0.69 | 0.69 | 0.69 |
| 14 | 125139656 | 62 | 90 | 72 | 59 | 0.50 | 0.72 | 0.58 | 0.47 |
| 15 | 104073951 | 72 | 102 | 49 | 58 | 0.69 | 0.98 | 0.47 | 0.56 |
| 16 | 98008968 | 109 | 119 | 101 | 78 | 1.11 | 1.21 | 1.03 | 0.80 |
| 17 | 95294699 | 106 | 118 | 100 | 100 | 1.11 | 1.24 | 1.05 | 1.05 |
| 18 | 90720763 | 56 | 51 | 52 | 39 | 0.62 | 0.56 | 0.57 | 0.43 |
| 19 | 61420004 | 49 | 68 | 39 | 44 | 0.80 | 1.11 | 0.63 | 0.72 |
| X | 169476592 | 60 | 69 | 79 | 61 | 0.35 | 0.41 | 0.47 | 0.36 |
| Y | 91455967 | 2 | 1 | 2 | 1 | 0.02 | 0.01 | 0.02 | 0.01 |

**Table S6b. Number of Differentially Expressed Genes (DEGs) found per chromosome and percentage of DEGs per chromosome after correcting by chromosome length.**

| **Gene** | **Chr** | **Start** | **Feature Type MGI** | **MGI Gene/ Marker ID** | **Ts65Dn EC** | **Ts65Dn HC** | **Ts66Yah EC** | **Ts66Yah HC** |
| --- | --- | --- | --- | --- | --- | --- | --- | --- |
| *Gm10232* | 17 | 3044014 | pseudogene | MGI:3641637 | 1.23 | 1.11 | 0.86 | 0.83 |
| *Pisd-ps2* | 17 | 3076578 | pseudogene | MGI:3612472 | 1.41 | 1.43 | 0.96 | 1.1 |
| *Scaf8* | 17 | 3114972 | protein coding gene | MGI:1925212 | 1.35 | 1.4 | 1.02 | 1.03 |
| *Tiam2* | 17 | 3326573 | protein coding gene | MGI:1344338 | 1.73 | 1.49 | 1.01 | 1.01 |
| *Gm7043* | 17 | 3461800 | pseudogene | MGI:3647242 | 0.68 | 0.71 | 0.84 | 0.89 |
| *Tfb1m* | 17 | 3519256 | protein coding gene | MGI:2146851 | 1.47 | 1.55 | 1.04 | 1.13 |
| *Cldn20* | 17 | 3532554 | protein coding gene | MGI:3646757 | 1.55 | 1.45 | 0.82 | 1.39 |
| *1700102H20Rik* | 17 | 3557824 | lncRNA gene | MGI:1915480 | 1.03 | 1.54 | 0.58 | 1.15 |
| *Arid1b* | 17 | 4994332 | protein coding gene | MGI:1926129 | 1.42 | 1.48 | 1.02 | 1.01 |
| *Gm15599* | 17 | 5110766 | pseudogene | MGI:3783046 | 0.82 | 0.78 | 1.63 | 0.71 |
| *Gm29050* | 17 | 5388763 | lncRNA gene | MGI:5579756 | 2.49 | 1.01 | 1.82 | 0.64 |
| *Tmem242* | 17 | 5410870 | protein coding gene | MGI:1917794 | 1.38 | 1.34 | 0.94 | 0.93 |
| *Zdhhc14* | 17 | 5492557 | protein coding gene | MGI:2653229 | 1.71 | 1.63 | 0.94 | 0.98 |
| *Gm26595* | 17 | 5817472 | pseudogene | MGI:5477089 | 0.91 | 1.01 | 1.00 | 1.04 |
| *Gm26622* | 17 | 5837968 | pseudogene | MGI:5477116 | 1.31 | 0.86 | 0.85 | 1.33 |
| *Snx9* | 17 | 5841329 | protein coding gene | MGI:1913866 | 1.47 | 1.39 | 0.99 | 0.94 |
| *Synj2* | 17 | 5941280 | protein coding gene | MGI:1201671 | 1.48 | 1.62 | 1.12 | 1.18 |
| *Serac1* | 17 | 6042196 | protein coding gene | MGI:2447813 | 1.6 | 1.58 | 1.00 | 1.01 |
| *Gtf2h5* | 17 | 6079786 | protein coding gene | MGI:107227 | 1.52 | 1.54 | 0.96 | 1.01 |
| *Tulp4* | 17 | 6106437 | protein coding gene | MGI:1916092 | 1.3 | 1.3 | 1.13 | 1.08 |
| *Gm15590* | 17 | 6131854 | pseudogene | MGI:3831433 | 0.84 | 0.87 | 0.82 | 1.08 |
| *Tmem181a* | 17 | 6256860 | protein coding gene | MGI:1924356 | 0.92 | 1.09 | 1.67 | 1.34 |
| *Dynlt1a* | 17 | 6306340 | protein coding gene | MGI:3807506 | 0.65 | 0.78 | 1.39 | 1.47 |
| *Dynlt1b* | 17 | 6430112 | protein coding gene | MGI:98643 | 1.54 | 1.29 | 0.88 | 0.86 |
| *Tmem181b-ps* | 17 | 6438524 | pseudogene | MGI:3779544 | 1.42 | 1.28 | 0.71 | 0.73 |
| *Dynlt1c* | 17 | 6601671 | protein coding gene | MGI:3807476 | 2.76 | 1.01 | 0.62 | 0.56 |
| *Tmem181c-ps* | 17 | 6613753 | pseudogene | MGI:3780993 | 2.6 | 2.16 | 1.02 | 0.57 |
| *Dynlt1f* | 17 | 6646602 | protein coding gene | MGI:3780996 | 1.62 | 1.38 | 1.15 | 1.05 |
| *Sytl3* | 17 | 6659093 | protein coding gene | MGI:1933367 | 1.5 | 2.01 | 0.59 | 0.54 |
| *Ezr* | 17 | 6738041 | protein coding gene | MGI:98931 | 1.31 | 1.44 | 0.9 | 0.96 |
| *Rsph3b* | 17 | 6904413 | protein coding gene | MGI:3630308 | 1.26 | 1.15 | 0.99 | 1.00 |
| *Tagap1* | 17 | 6955011 | protein coding gene | MGI:1919786 | 1.23 | 1.17 | 1.02 | 1.04 |
| *Rnaset2b* | 17 | 6970634 | protein coding gene | MGI:3702087 | 0.89 | 1.12 | 1.04 | 1.71 |
| *Rps6ka2* | 17 | 7170115 | protein coding gene | MGI:1342290 | 1.4 | 1.21 | 1.04 | 1.17 |
| *Fndc1* | 17 | 7738569 | protein coding gene | MGI:1915905 | 1.7 | 1.35 | 1.14 | 0.88 |
| *E430024P14Rik* | 17 | 7881106 | lncRNA gene | MGI:2445079 | 2.06 | 1.55 | 0.89 | 1.04 |
| *Tagap* | 17 | 7926000 | protein coding gene | MGI:3615484 | 2.6 | 2.65 | 0.68 | 0.67 |
| *Rsph3a* | 17 | 7945614 | protein coding gene | MGI:1914082 | 1.96 | 1.53 | 1.09 | 1.25 |
| *Rnaset2a* | 17 | 8115445 | protein coding gene | MGI:1915445 | 1.54 | 1.74 | 0.74 | 0.95 |
| *Fgfr1op* | 17 | 8165501 | protein coding gene | MGI:1922546 | 1.51 | 1.33 | 1.01 | 0.96 |
| *Ccr6* | 17 | 8236043 | protein coding gene | MGI:1333797 | 9.69 | 9.74 | 0.58 | 0.5 |
| *Mpc1* | 17 | 8282904 | protein coding gene | MGI:1915240 | 1.15 | 1.22 | 1.18 | 1.42 |
| *4930506C21Rik* | 17 | 8293366 | lncRNA gene | MGI:1922310 | 1.75 | 1.51 | 1.11 | 1.00 |
| *Sft2d1* | 17 | 8311102 | protein coding gene | MGI:1918689 | 1.34 | 1.31 | 0.96 | 0.88 |
| *Prr18* | 17 | 8337459 | protein coding gene | MGI:2443403 | 1.51 | 1.56 | 1.2 | 0.97 |
| *T2* | 17 | 8355992 | protein coding gene | MGI:104658 | 2.22 | 1.22 | 0.85 | 1,00 |
| *T* | 17 | 8434423 | protein coding gene | MGI:98472 | 1.71 | 1.89 | 1.00 | 1.34 |
| *Pde10a* | 17 | 8525372 | protein coding gene | MGI:1345143 | 1.69 | 1.48 | 0.91 | 0.98 |
| *Gm15425* | 17 | 8564258 | pseudogene | MGI:3705642 | 0.79 | 0.84 | 1.05 | 1.00 |
| *Gm17087* | 17 | 8565852 | protein coding gene | MGI:4937914 | 1.23 | 1.19 | 1.08 | 0.92 |
| *1700010I14Rik* | 17 | 8988333 | protein coding gene | MGI:1914181 | 1.37 | 1.29 | 1.39 | 1.32 |
| *6530411M01Rik* | 17 | 9147719 | lncRNA gene | MGI:1915041 | 0.98 | 1.27 | 1.35 | 1.28 |
| *Gm8492* | 17 | 9320395 | pseudogene | MGI:3643249 | 1.03 | 0.97 | 1.05 | 1.05 |
| *Mrpl39* | 16 | 84717576 | protein coding gene | MGI:1351620 | 1.41 | 1.44 | 1.49 | 1.50 |
| *Jam2* | 16 | 84774123 | protein coding gene | MGI:1933820 | 1.43 | 1.49 | 1.63 | 1.56 |
| *Atp5j* | 16 | 84827866 | protein coding gene | MGI:107777 | 1.42 | 1.56 | 1.57 | 1.45 |
| *Gabpa* | 16 | 84834925 | protein coding gene | MGI:95610 | 1.38 | 1.22 | 1.42 | 1.48 |
| *App* | 16 | 84949685 | protein coding gene | MGI:88059 | 1.44 | 1.57 | 1.49 | 1.53 |
| *Gm10791* | 16 | 84972214 | lncRNA gene | MGI:3641949 | 1.13 | 1.48 | 1.38 | 1.39 |
| *Cyyr1* | 16 | 85421533 | protein coding gene | MGI:2152187 | 1.24 | 1.42 | 1.56 | 1.53 |
| *Adamts1* | 16 | 85793827 | protein coding gene | MGI:109249 | 1.30 | 1.46 | 1.81 | 1.56 |
| *Adamts5* | 16 | 85856173 | protein coding gene | MGI:1346321 | 1.48 | 1.72 | 1.59 | 1.60 |
| *N6amt1* | 16 | 87354185 | protein coding gene | MGI:1915018 | 1.38 | 1.26 | 1.15 | 1.11 |
| *Ltn1* | 16 | 87376651 | protein coding gene | MGI:1926163 | 1.38 | 1.41 | 1.72 | 1.58 |
| *Rwdd2b* | 16 | 87433407 | protein coding gene | MGI:1858215 | 1.57 | 1.48 | 1.62 | 1.59 |
| *Usp16* | 16 | 87454703 | protein coding gene | MGI:1921362 | 1.50 | 1.36 | 1.51 | 1.50 |
| *Cct8* | 16 | 87483326 | protein coding gene | MGI:107183 | 1.38 | 1.46 | 1.61 | 1.52 |
| *Rpl31-ps4* | 16 | 87549062 | pseudogene | MGI:3649126 | 0.78 | 0.93 | 0.92 | 1.44 |
| *Map3k7cl* | 16 | 87553330 | protein coding gene | MGI:2446584 | 1.20 | 1.78 | 1.29 | 0.97 |
| *Bach1* | 16 | 87698945 | protein coding gene | MGI:894680 | 1.41 | 1.37 | 1.50 | 1.35 |
| *2810407A14Rik* | 16 | 87784075 | lncRNA gene | MGI:1917461 | 1.46 | 1.47 | 1.98 | 2.04 |
| *Grik1* | 16 | 87895900 | protein coding gene | MGI:95814 | 1.49 | 1.40 | 1.37 | 1.39 |
| *Cldn17* | 16 | 88505807 | protein coding gene | MGI:2652030 | 1.00 | 2.05 | 1.00 | 1.63 |
| *Cldn8* | 16 | 88560828 | protein coding gene | MGI:1859286 | 1.00 | 0.92 | 1.00 | 0.96 |
| *Tiam1* | 16 | 89787111 | protein coding gene | MGI:103306 | 1.56 | 1.36 | 1.60 | 1.51 |
| *Sod1* | 16 | 90220754 | protein coding gene | MGI:98351 | 1.62 | 1.74 | 1.43 | 1.39 |
| *Scaf4* | 16 | 90225680 | protein coding gene | MGI:2146350 | 1.42 | 1.42 | 1.24 | 1.28 |
| *Hunk* | 16 | 90386013 | protein coding gene | MGI:1347352 | 1.27 | 1.53 | 1.71 | 1.60 |
| *1110008E08Rik* | 16 | 90554164 | lncRNA gene | MGI:1915750 | 0.94 | 1.21 | 1.03 | 2.88 |
| *Mis18a* | 16 | 90719312 | protein coding gene | MGI:1913828 | 1.63 | 1.29 | 1.56 | 1.45 |
| *Mrap* | 16 | 90738207 | protein coding gene | MGI:1924287 | 1.45 | 1.29 | 1.06 | 1.80 |
| *Urb1* | 16 | 90751527 | protein coding gene | MGI:2146468 | 1.40 | 1.49 | 1.55 | 1.72 |
| *Eva1c* | 16 | 90826719 | protein coding gene | MGI:1918217 | 1.50 | 1.75 | 1.35 | 1.74 |
| *Synj1* | 16 | 90936092 | protein coding gene | MGI:1354961 | 1.54 | 1.59 | 1.64 | 1.62 |
| *Paxbp1* | 16 | 91014037 | protein coding gene | MGI:1914617 | 1.44 | 1.42 | 1.40 | 1.59 |
| *4932438H23Rik* | 16 | 91053935 | protein coding gene | MGI:1921637 | 1.26 | 1.41 | 1.63 | 1.72 |
| *H3f3a-ps2* | 16 | 91114189 | pseudogene | MGI:1101758 | 1.00 | 0.85 | 0.87 | 0.60 |
| *Olig2* | 16 | 91225457 | protein coding gene | MGI:1355331 | 1.47 | 1.42 | 1.24 | 0.94 |
| *Olig1* | 16 | 91269772 | protein coding gene | MGI:1355334 | 1.47 | 1.43 | 1.41 | 0.97 |
| *Ifnar2* | 16 | 91372783 | protein coding gene | MGI:1098243 | 1.46 | 1.55 | 1.54 | 1.56 |
| *Il10rb* | 16 | 91406164 | protein coding gene | MGI:109380 | 1.34 | 1.53 | 1.42 | 1.58 |
| *Ifnar1* | 16 | 91485238 | protein coding gene | MGI:107658 | 1.49 | 1.48 | 1.62 | 1.60 |
| *Ifngr2* | 16 | 91547072 | protein coding gene | MGI:107654 | 1.69 | 1.65 | 1.65 | 1.60 |
| *Tmem50b* | 16 | 91574503 | protein coding gene | MGI:1925225 | 1.45 | 1.58 | 1.55 | 1.46 |
| *Dnajc28* | 16 | 91614254 | protein coding gene | MGI:2181053 | 1.33 | 1.31 | 1.48 | 1.57 |
| *Gart* | 16 | 91621186 | protein coding gene | MGI:95654 | 1.48 | 1.59 | 1.48 | 1.50 |
| *Son* | 16 | 91647506 | protein coding gene | MGI:98353 | 1.21 | 1.30 | 1.30 | 1.52 |
| *Donson* | 16 | 91676808 | protein coding gene | MGI:1890621 | 1.53 | 1.36 | 1.55 | 1.55 |
| *Atp5o* | 16 | 91684398 | protein coding gene | MGI:106341 | 1.43 | 1.46 | 1.61 | 1.62 |
| *Gm10785* | 16 | 91688898 | lncRNA gene | MGI:3642149 | 1.32 | 1.36 | 1.87 | 1.35 |
| *Cryzl1* | 16 | 91689322 | protein coding gene | MGI:1913859 | 1.36 | 1.42 | 1.55 | 1.45 |
| *Itsn1* | 16 | 91729281 | protein coding gene | MGI:1338069 | 1.52 | 1.49 | 1.46 | 1.41 |
| *Mrps6* | 16 | 92058270 | protein coding gene | MGI:2153111 | 1.73 | 1.60 | 1.44 | 1.47 |
| *Slc5a3* | 16 | 92058322 | protein coding gene | MGI:1858226 | 1.22 | 1.15 | 1.51 | 1.52 |
| *Kcne2* | 16 | 92292389 | protein coding gene | MGI:1891123 | 0.99 | 1.99 | 0.98 | 1.55 |
| *Smim11* | 16 | 92301286 | protein coding gene | MGI:1916186 | 1.40 | 1.26 | 1.47 | 1.51 |
| *Kcne1* | 16 | 92345982 | protein coding gene | MGI:96673 | 1.15 | 2.93 | 1.30 | 1.33 |
| *Rcan1* | 16 | 92391953 | protein coding gene | MGI:1890564 | 1.55 | 1.56 | 1.47 | 1.58 |
| *2410124H12Rik* | 16 | 92478742 | lncRNA gene | MGI:1924035 | 1.16 | 1.11 | 0.93 | 1.94 |
| *Clic6* | 16 | 92485736 | protein coding gene | MGI:2146607 | 1.73 | 1.87 | 1.31 | 1.49 |
| *Runx1* | 16 | 92601466 | protein coding gene | MGI:99852 | 1.16 | 1.08 | 0.99 | 1.08 |
| *Setd4* | 16 | 93583457 | protein coding gene | MGI:2136890 | 1.53 | 1.41 | 1.52 | 1.45 |
| *Cbr1* | 16 | 93605853 | protein coding gene | MGI:88284 | 1.51 | 1.47 | 1.59 | 1.32 |
| *Cbr3* | 16 | 93683215 | protein coding gene | MGI:1309992 | 1.51 | 1.57 | 1.39 | 1.42 |
| *Morc3* | 16 | 93832121 | protein coding gene | MGI:2136841 | 1.33 | 1.33 | 1.53 | 1.56 |
| *Chaf1b* | 16 | 93883901 | protein coding gene | MGI:1314881 | 1.22 | 1.98 | 1.67 | 1.48 |
| *Cldn14* | 16 | 93919031 | protein coding gene | MGI:1860425 | 1.90 | 1.39 | 1.61 | 1.51 |
| *Sim2* | 16 | 94084931 | protein coding gene | MGI:98307 | 1.20 | 1.56 | 1.90 | 1.71 |
| *Hlcs* | 16 | 94128882 | protein coding gene | MGI:894646 | 1.59 | 1.40 | 1.44 | 1.43 |
| *Ripply3* | 16 | 94328420 | protein coding gene | MGI:2181192 | 2.01 | 1.57 | 1.43 | 2.00 |
| *Pigp* | 16 | 94358763 | protein coding gene | MGI:1860433 | 1.45 | 1.45 | 1.45 | 1.41 |
| *Ttc3* | 16 | 94370618 | protein coding gene | MGI:1276539 | 1.49 | 1.50 | 1.51 | 1.53 |
| *Gm23692* | 16 | 94429073 | snRNA gene | MGI:5453469 | 1.00 | 1.00 | 1.14 | 1.00 |
| *Dyrk1a* | 16 | 94570010 | protein coding gene | MGI:1330299 | 1.36 | 1.41 | 1.53 | 1.45 |
| *Kcnj6* | 16 | 94748636 | protein coding gene | MGI:104781 | 1.54 | 1.50 | 1.55 | 1.71 |
| *Kcnj15* | 16 | 95257558 | protein coding gene | MGI:1310000 | 1.69 | 3.34 | 1.90 | 1.80 |
| *Erg* | 16 | 95359169 | protein coding gene | MGI:95415 | 1.06 | 1.19 | 0.99 | 1.11 |
| *Ets2* | 16 | 95702075 | protein coding gene | MGI:95456 | 1.38 | 1.38 | 1.37 | 1.43 |
| *Psmg1* | 16 | 95979933 | protein coding gene | MGI:1860263 | 1.51 | 1.56 | 1.40 | 1.31 |
| *Brwd1* | 16 | 95992092 | protein coding gene | MGI:1890651 | 1.43 | 1.43 | 1.50 | 1.47 |
| *Hmgn1* | 16 | 96120618 | protein coding gene | MGI:96120 | 1.40 | 1.40 | 1.51 | 1.49 |
| *Wrb* | 16 | 96145407 | protein coding gene | MGI:2136882 | 1.53 | 1.55 | 1.58 | 1.57 |
| *Lca5l* | 16 | 96158407 | protein coding gene | MGI:3041157 | 1.60 | 1.41 | 1.59 | 1.62 |
| *Sh3bgr* | 16 | 96200450 | protein coding gene | MGI:1354740 | 2.87 | 1.29 | 1.34 | 1.68 |
| *B3galt5* | 16 | 96235801 | protein coding gene | MGI:2136878 | 1.31 | 1.33 | 1.54 | 1.49 |
| *Igsf5* | 16 | 96361668 | protein coding gene | MGI:1919308 | 1.42 | 1.36 | 2.12 | 2.05 |
| *Itgb2l* | 16 | 96422288 | protein coding gene | MGI:1277979 | 1.63 | 1.34 | 0.94 | 1.12 |
| *Pcp4* | 16 | 96467606 | protein coding gene | MGI:97509 | 1.57 | 1.12 | 1.15 | 1.18 |
| *Dscam* | 16 | 96590840 | protein coding gene | MGI:1196281 | 1.50 | 1.50 | 1.63 | 1.53 |
| *Bace2* | 16 | 97356742 | protein coding gene | MGI:1860440 | 1.45 | 1.66 | 1.61 | 1.76 |
| *Mx1* | 16 | 97447035 | polymorphic pseudogene | MGI:97243 | 1.55 | 1.50 | 2.22 | 1.89 |
| *Mx2* | 16 | 97535308 | polymorphic pseudogene | MGI:97244 | 1.54 | 1.55 | 1.95 | 1.59 |
| *Tmprss2* | 16 | 97564682 | protein coding gene | MGI:1354381 | 1.19 | 0.74 | 1.42 | 2.29 |
| *Ripk4* | 16 | 97741933 | protein coding gene | MGI:1919638 | 3.05 | 2.38 | 2.44 | 2.01 |
| *Prdm15* | 16 | 97791467 | protein coding gene | MGI:1930121 | 1.24 | 1.22 | 1.42 | 1.36 |
| *C2cd2* | 16 | 97855209 | protein coding gene | MGI:1891883 | 1.37 | 1.44 | 1.72 | 1.61 |
| *Zbtb21* | 16 | 97943357 | protein coding gene | MGI:1927240 | 1.22 | 1.30 | 1.45 | 1.42 |

**Table S7. Expression of genes from the centromeric part of the Mmu17 and the telomeric part of Mmu16 in the Ts66Yah and Ts65Dn Hippocampy (HC) or Entorhinal cortex (EC) (in red with a FC >1.2).**

|  | Ts66Yah HIP | | Ts66Yah EC | | Ts65Dn HIP | | Ts65Dn EC | |
| --- | --- | --- | --- | --- | --- | --- | --- | --- |
| Group of pathways | pathways/ group | % pathways per group | nb total pathways per group | % pathways per group | nb total pathways per group | % pathways per group | nb total pathways per group | % pathways per group |
| Synaptic | 12 | 3,72 | 4 | 10 | 8 | 1,51 | 10 | 6,62 |
| GABA system | 1 | 0,31 |  |  | 1 | 0,19 | 5 | 3,31 |
| Myelin sheat:SNARE complex |  |  |  |  | 2 | 0,38 |  |  |
| Behaviour | 3 | 0,93 |  |  | 2 | 0,38 | 1 | 0,66 |
| Synaptic: Other pathways | 12 | 3,72 | 4 | 10 | 4 | 0,75 | 10 | 6,62 |
| Transcription & epigenomic regulation | 22 | 6,81 | 2 | 5 | 24 | 4,53 | 10 | 6,62 |
| Enzymes activity | 52 | 16,1 |  |  | 88 | 16,6 | 40 | 26,49 |
| Mitochondria | 1 | 0,31 |  |  |  |  | 5 | 3,31 |
| Ribosome | 3 | 0,93 |  |  | 6 | 1,13 | 4 | 2,65 |
| Compounds binding | 16 | 4,95 | 3 | 7,5 | 28 | 5,28 | 19 | 12,58 |
| Cell Structure & organelles | 12 | 3,72 | 7 | 17,5 | 82 | 15,47 | 40 | 26,49 |
| Phospho-Kinase | 8 | 2,48 |  |  | 8 | 1,51 | 3 | 1,99 |
| Hormone regulation | 4 | 1,24 |  |  |  |  |  |  |
| Post-translation modification | 1 | 0,31 |  |  | 4 | 0,75 |  |  |
| Replication | 4 | 1,24 | 1 | 2,5 | 7 | 1,32 |  |  |
| Morphogenesis & development | 32 | 9,91 | 14 | 35 | 89 | 16,79 | 9 | 5,96 |
| Host response & defense | 127 | 39,32 | 2 | 5 | 174 | 32,83 | 18 | 11,92 |
| Metabolism | 67 | 20,74 | 9 | 22,5 | 118 | 22,26 | 33 | 21,85 |
| Degradation system | 1 | 0,31 |  |  | 7 | 1,32 | 7 | 4,64 |
| Cardiovascular & homeostasis | 16 | 4,95 |  |  | 19 | 3,58 | 4 | 2,65 |
| Apoptosis & cell death | 2 | 0,62 |  |  | 9 | 1,7 |  |  |
| Connective-tissue cells |  |  | 2 | 5 | 11 | 2,08 | 1 | 0,66 |
| Fibers and cytoskeleton | 5 | 1,55 |  |  | 5 | 0,94 | 18 | 11,92 |
| Stimulus detection | 2 | 0,62 |  |  | 4 | 0,75 |  |  |
| Signaling pathways | 14 | 4,33 | 2 | 5 | 30 | 5,66 | 6 | 3,97 |
| Channels and location signals | 55 | 17,03 |  |  | 81 | 15,28 | 14 | 9,27 |
| Sexual development & embryogenesis | 2 | 0,62 |  |  |  |  | 3 | 1,99 |
| Aging |  |  | 1 | 2,5 |  |  |  |  |

**Table S08 Summary of the functional pathways found in the models, the number of pathways grouped on each meta-pathway and the percentage of pathways inside each meta-pathway considering the total number of pathways.**

### Supplementary figure

A

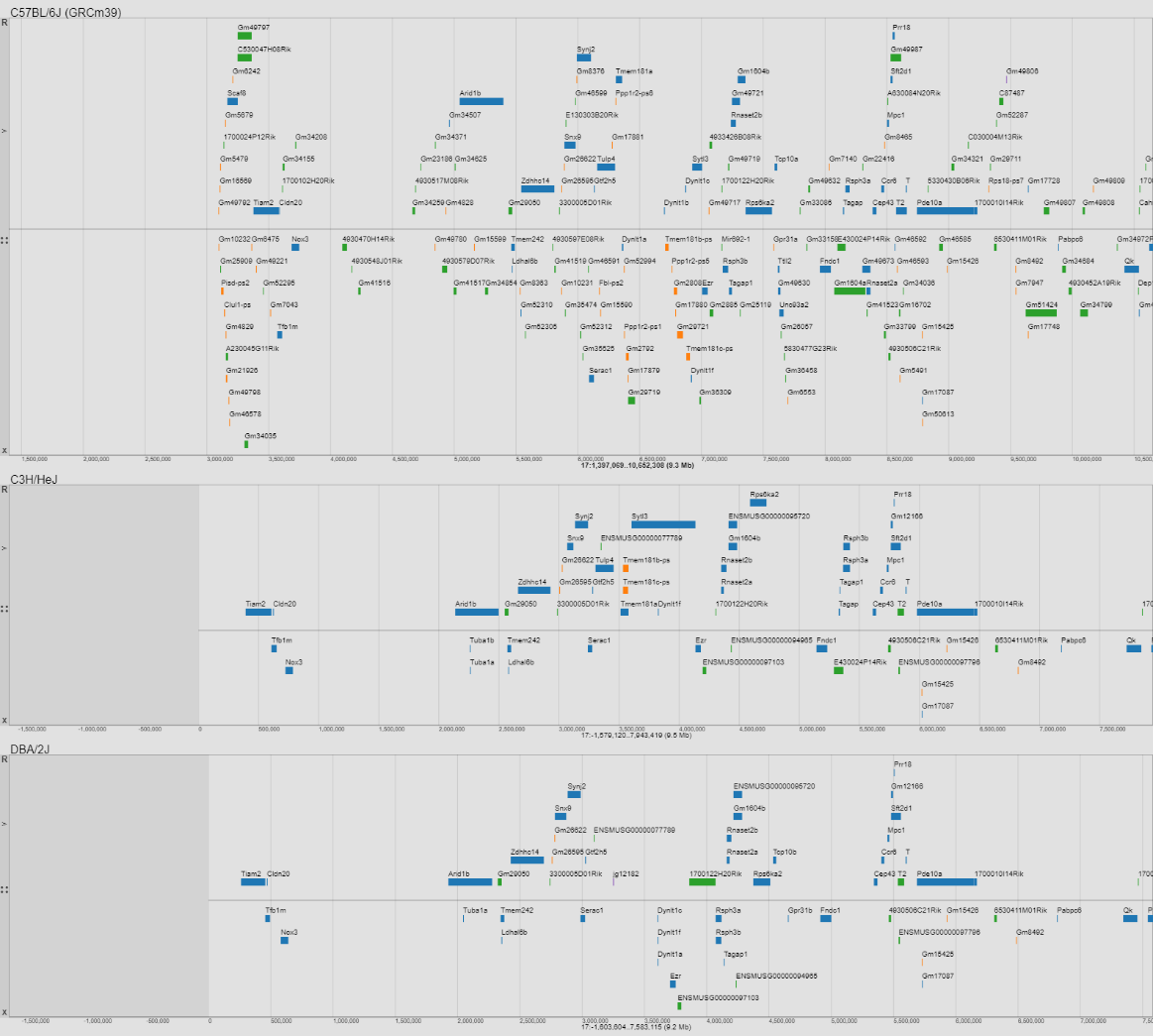

B

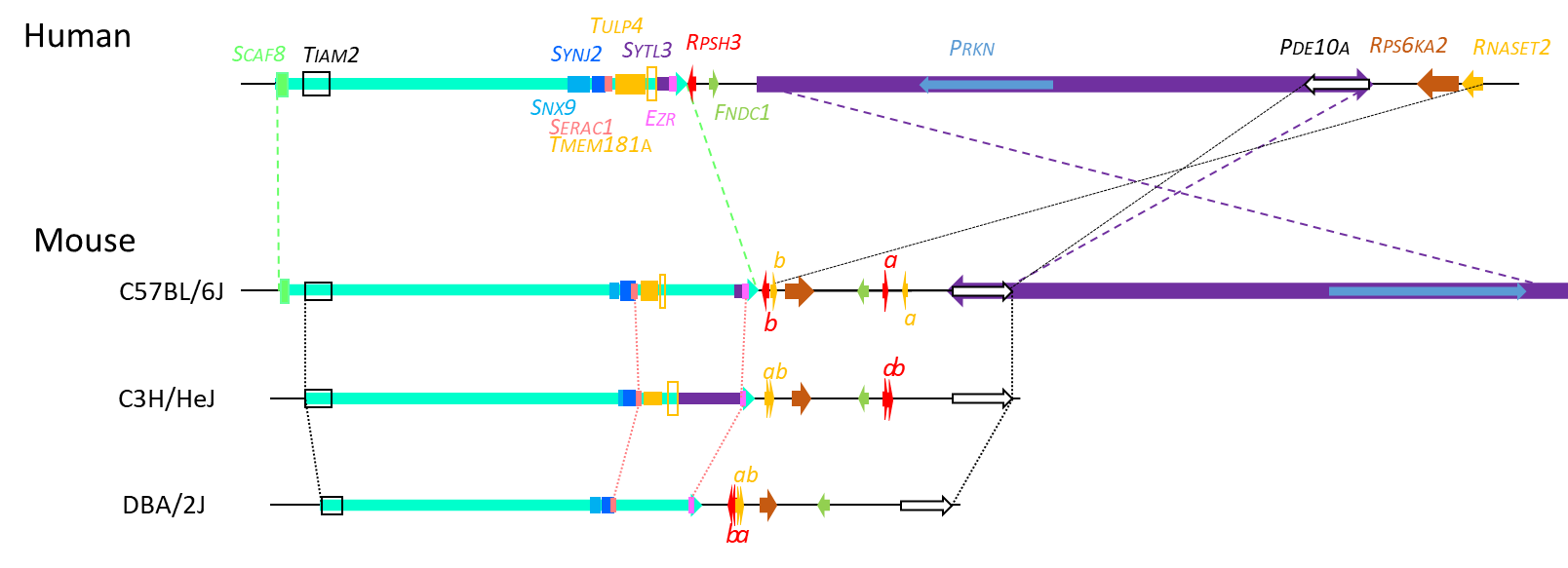

**Fig. S1: The genomic organisation of the *Scaf8- Pde10a* region found on the centromeric part of the mouse chromosome 17 is conserved, but reorganized, in the C57BL/6J, C3H/HeJ and DBA/2J mouse strains, and showed several gene reorganization in the human genome.** A. The copy was derived from the multiple genome viewer available at the Mouse genome informatics ([www.informatics.jax.org/Mgv](http://www.informatics.jax.org/Mgv)) including the region sequenced on C3H/HeJ::17:-1579120..7943419, on the C57BL/6::17:1397069..10652308 and DBA/2J::17:-1603604..7583115. with the GRCm39 reference genome. B. Schematic representation of the general organization of the *Scaf8-Pde10a* in the 3 mouse lines and in the human genome showing both preserved and altered loci.

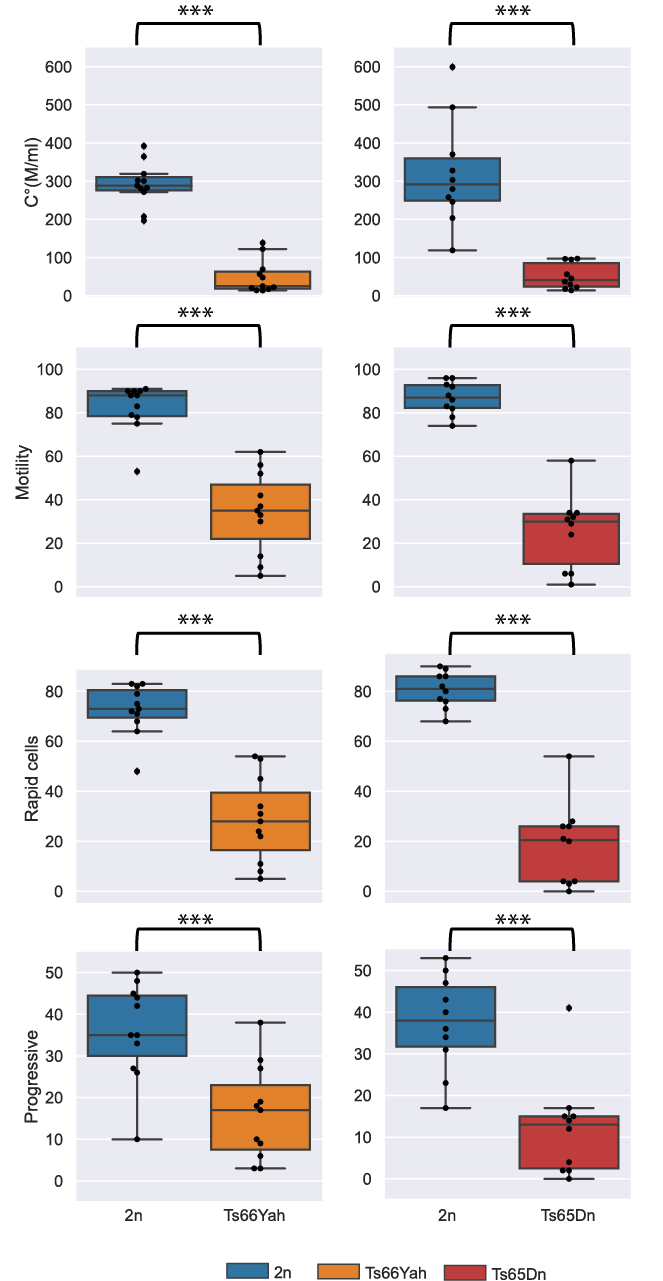

**Fig. S2. Comparison of sperm analysis in the Ts66Yah and Ts65Dn DS models**. Both models presented significant lower pattern of sperm quality in males as observed here with the spermatozoa concentration (C°, millions/ml), the percentage of motile cells (Motility %), the rapid cells (%) and the progressivity (%) in Ts66Yah (n = 11 for 2n control littermates and n = 11 for Ts animals; left panel) and Ts65Dn (n = 10 for 2n control littermates and n=10 for Ts animals; right panel) respectively. All parameters were significantly lower in trisomic mice. Box plots with the median and quartiles. Statistical significance of differences between genotype was inferred by a two-tailed T test or Kruskal-Wallis non-parametric test.***p<0.001)

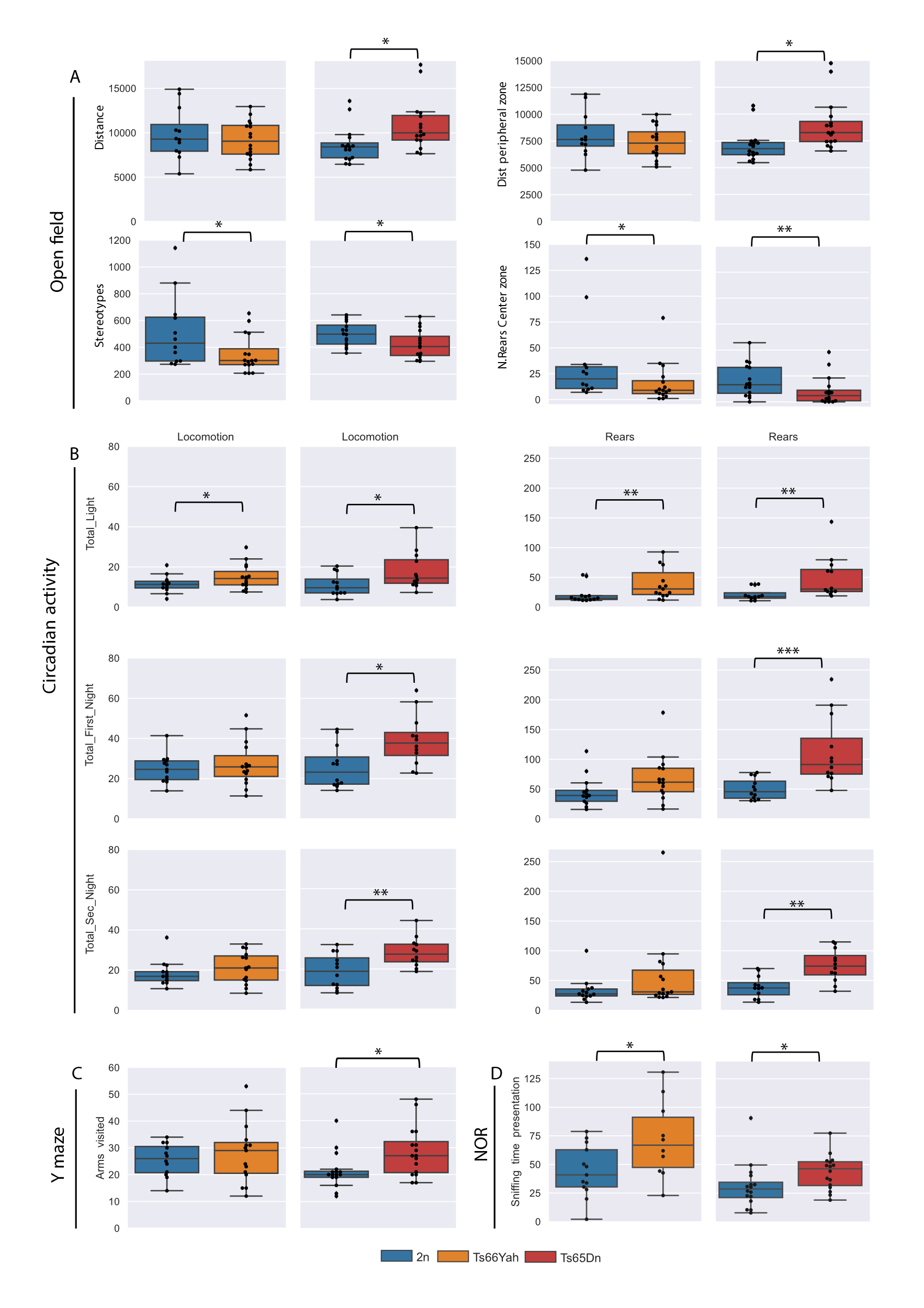

**Fig. S3: Activities measured in different paradigms for the Ts66Yah and Ts65Dn males.** A. In the open field, while Ts66Yah trisomic mice travelled the same distance as the 2n control mice, the Ts65Dn mice showed more activity in this paradigm. The number of stereotypes was increased in both trisomic mice. Only the Ts65Dn mice presented an increase in anxiety-like behaviour with a reduced activity in the centre and increased activity in the peripheral zone. The experiment was done with 2n (n = 12) and trisomic (Ts; n = 16) males for Ts66Yah and 2n (n = 16) and Ts (n = 16) males for Ts65Dn. B. left and right panels illustrate the locomotion and the number of rears respectively in the circadian activity during light, first and second night phases of the test. The locomotion pattern of Ts66Yah mice was similar to the 2n control mice regardless of test phases. On the contrary, Ts65Dn mice exhibited an increase in locomotion activity during light, first and second night phases. Similarly, rearing activity (right panel) was increased for the Ts65Dn trisomic mice during all phases of the test, whereas Ts66Yah trisomic mice showed the same activity level than the 2n control group of mice. This experiment was done with 2n (n = 12) and trisomic (Ts; n = 16) males for Ts66Yah and 2n (n = 12) and Ts (n = 12) males for Ts65Dn. C. Only the Ts65Dn mice showed hyperactivity in the Y maze test with increased arms entries compare to the 2n group of mice. This experiment was done with 2n (n = 12) and trisomic (Ts; n = 16) males for Ts66Yah and 2n (n = 16) and Ts (n = 16) males for Ts65Dn. D. in the novel object recognition test, both group of mice with trisomy (13 2n/10 Ts66Yah and 16 2n/16 Ts65Dn) presented an increased sniffing time during the presentation phase of the test. Data obtained in activity tests highlight a hyperactivity and an increased in anxiety-like behavior of Ts65Dn trisomic mice, observations that were not found in the Ts66Yah trisomic mice. Box plot representation with the median and quartiles. Statistical significance of differences between genotype was inferred by a two-tailed T test or Kruskal-Wallis non-parametric test,* p<0.05, **p<0.01, ***p<0.001.

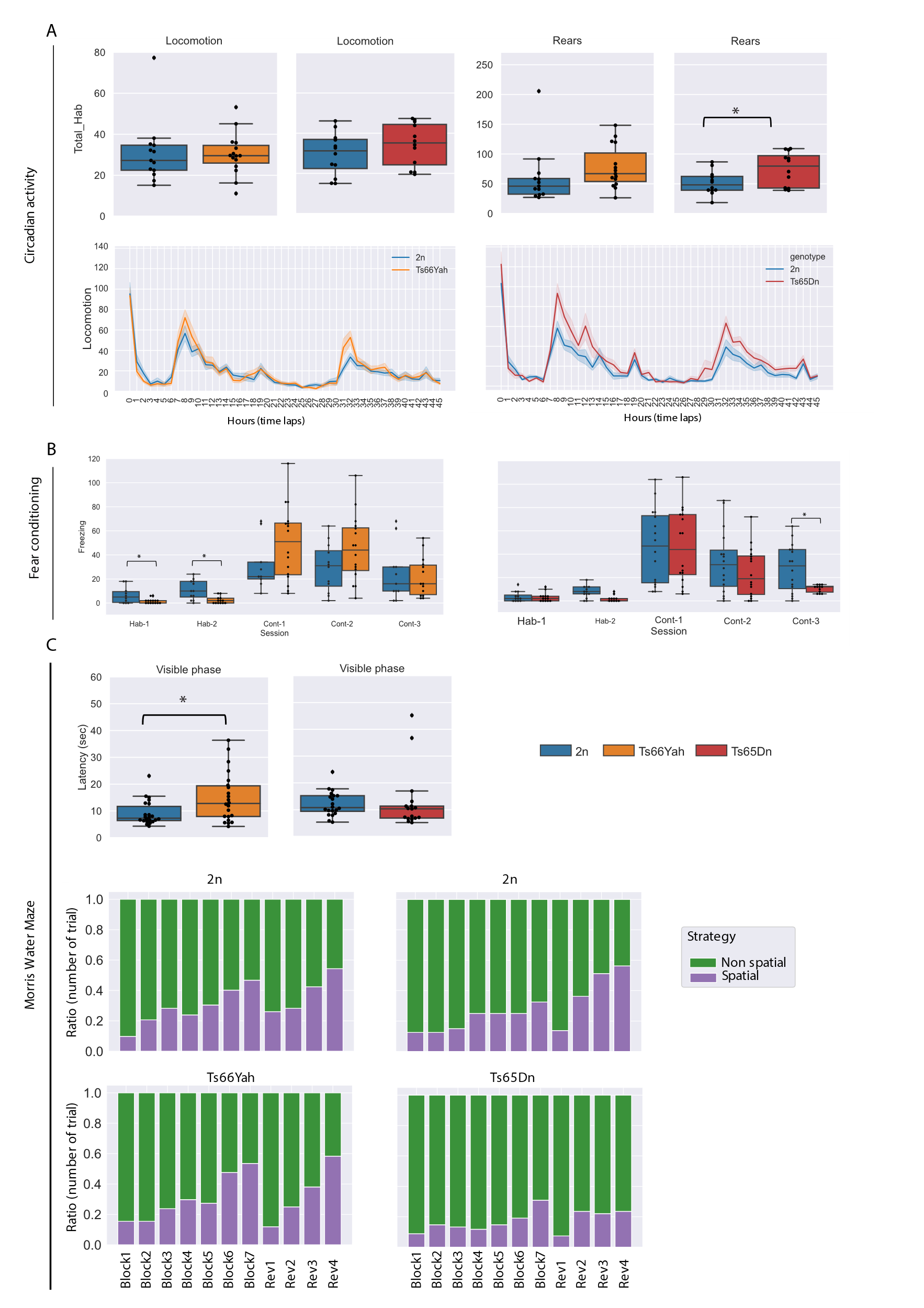

**Fig. S4: Investigating circadian activity, Morris water maze and fear conditioning comparing males from Ts66Yah and Ts65Dn DS models.** A. upper panel illustrate the locomotion and number of rears respectively in the circadian activity during habituation phases of the test. There was no difference between the 2n control and trisomic mice for the locomotion pattern. Rearing activity was increased for the Ts65Dn trisomic mice only. Lower panel represent locomotion circadian activity for Ts66Yah and Ts65Dn mutants, both lines had a similar pattern than the control littermates. B. freezing time in seconds during the different phases of the FC test. All animals increased their freezing time after electrical shock. During the context session, 24 hours after the conditioning, all the mice showed a high level of freezing time, except for the Ts65Dn in the last minutes of the test (Cont-3), when mice recovered their activity at normal level. Ts66Yah, n = 12 for 2n and n = 16 for Ts individuals and Ts65Dn n = 16 for control and n = 16 for Ts individuals. C. upper panel represents latency taking by the mice to locate the PF during the MWM visible session. Lower panel represents the PF searching strategies undertaken by each mouse models along the test. Only the Ts65Dn mice showed a low and constant level of spatial navigation path compared to the control mice and the Ts66Yah line. This experiments were done with 2n (n = 22) and trisomic (Ts; n = 22) Ts66Yah males and 2n (n = 20) and Ts (n = 18) Ts65Dn males. Box plots with the median and quartiles. Statistical significance of differences between genotype was inferred by a two-tailed T test or Kruskal-Wallis non-parametric test or ANOVA for repeated measures. * p<0.05, **p<0.01, ***p<0.001).

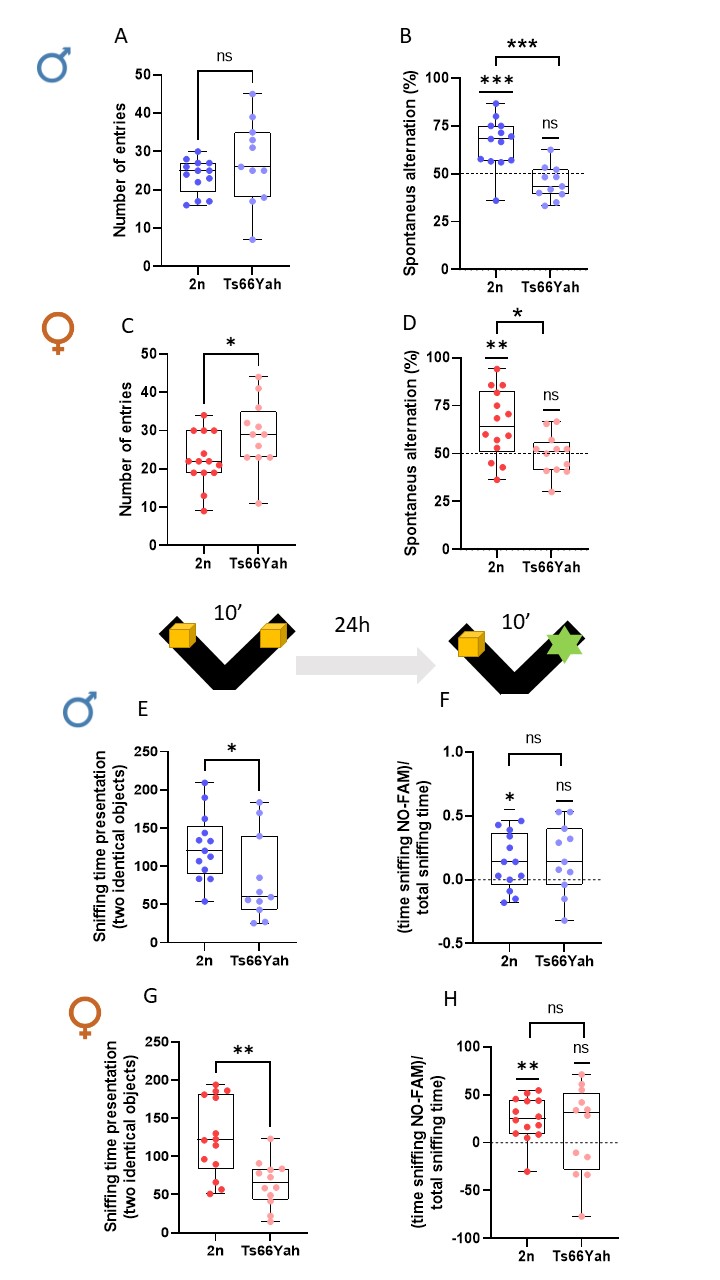

**Fig. S5 Reproducibility of memory phenotypes affecting spontaneous alternation and object discrimination in both males and females from the Ts66Yah mouse line with no major sex effect.**

During the Y maze test, the total number of entries (A-C) was not significantly different whereas the alternation rate (B-D) was lower in the Ts66Yah mice compared to control 2n littermates, and both sexes responded similarly. In the novel object recognition with 24-hour retention time, conducted in a V-maze (E-G), the total sniffing time of identical objects during the familiarization session was similar in males but not in females. During the test phase (F-H), the discrimination index showed no significant exploration of the novel object in both males and females Ts66Yah mice with no sex effect (ANOVA). Two tail one-sample t-test vs. 0% (hazard) showed that Ts66Yah male (p = 0.07) and female (p = 0.341) mice failed to recognize the new object. Two tail T-test for 2n vs. Ts66Yah comparisons. * p<0.05, **p<0.01, ***p<0.001. Each dot represents an individual. Blue dots represent male mice, and red dots represent female mice; dark colors for 2n mice and light colors for Ts66Yah mice (Ts66Yah, Male, n = 13 for 2n and n = 11 for Ts; Female n = 14 for 2n and n = 12 for Ts). Box plots represent the median and quartiles.

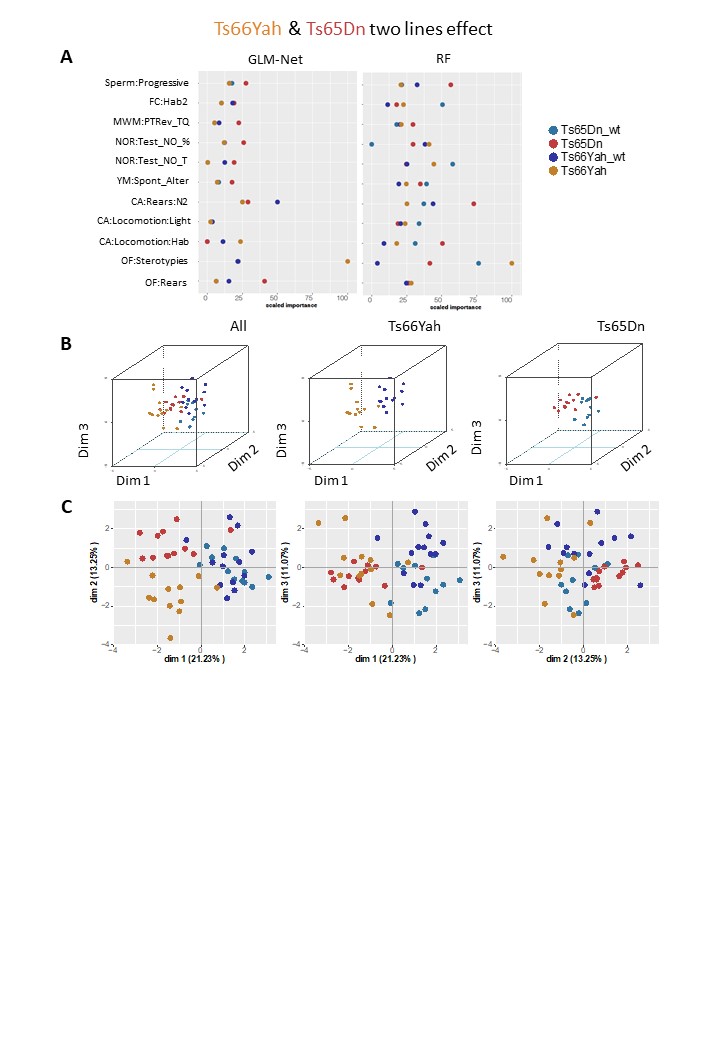

**Figure S6: Identification of discriminating phenotypic variables contributing the most to the genotype discrimination considering the two models Ts65Dn and Ts66Yah models together.** A. Identification of the power of each explanatory phenotypic variable to the genotype discrimination. The explanatory variables selected were the ones known to contribute more than 30% to the genotype discrimination. Those variables were identified using a principal component analysis (PCA) after applying a multi-factor analysis of mixed data (MFA) implemented using MFAmix function from the PCAmixdata R package. The relevance of those variables to the genotype discrimination was analyzed using two different statistical classifiers: Lasso and Elastic-Net Regularized Generalized Linear Models, noted as GLM-Net taken from the caret R package, and Random forest, noted RF taken from the caret R package. All measures of importance are scaled to have a maximum value of 100 in the variable contributing the most to the discrimination in the 2n control (wild type, wt) Ts66Yah. With GLM-Net and RF we can identify the importance of each variable to discriminate each genotype. B. 3D-PCA plots showing the individual animals clustering on the 3D space based on the PCA analyses performed with all the phenotypic variables and colored based on genotype and model as follows, in soft blue Ts65Dn control (wt), in red Ts65Dn DS trisomic, in dark blue the Ts66Yah wild-types (wt) and in medallion yellow Ts66Yah DS mutants. The upper panel shows all individuals from the two models, mixed genotypes, the middle plot shows only the Ts66Yah model and the bottom plot only the Ts65Dn. **C.** Individual component map. The distribution in 2D space of the individual observation coordinates calculated based on the PCA analysis performed after the MFA implemented using MFAmix function from the PCAmixdata R package.

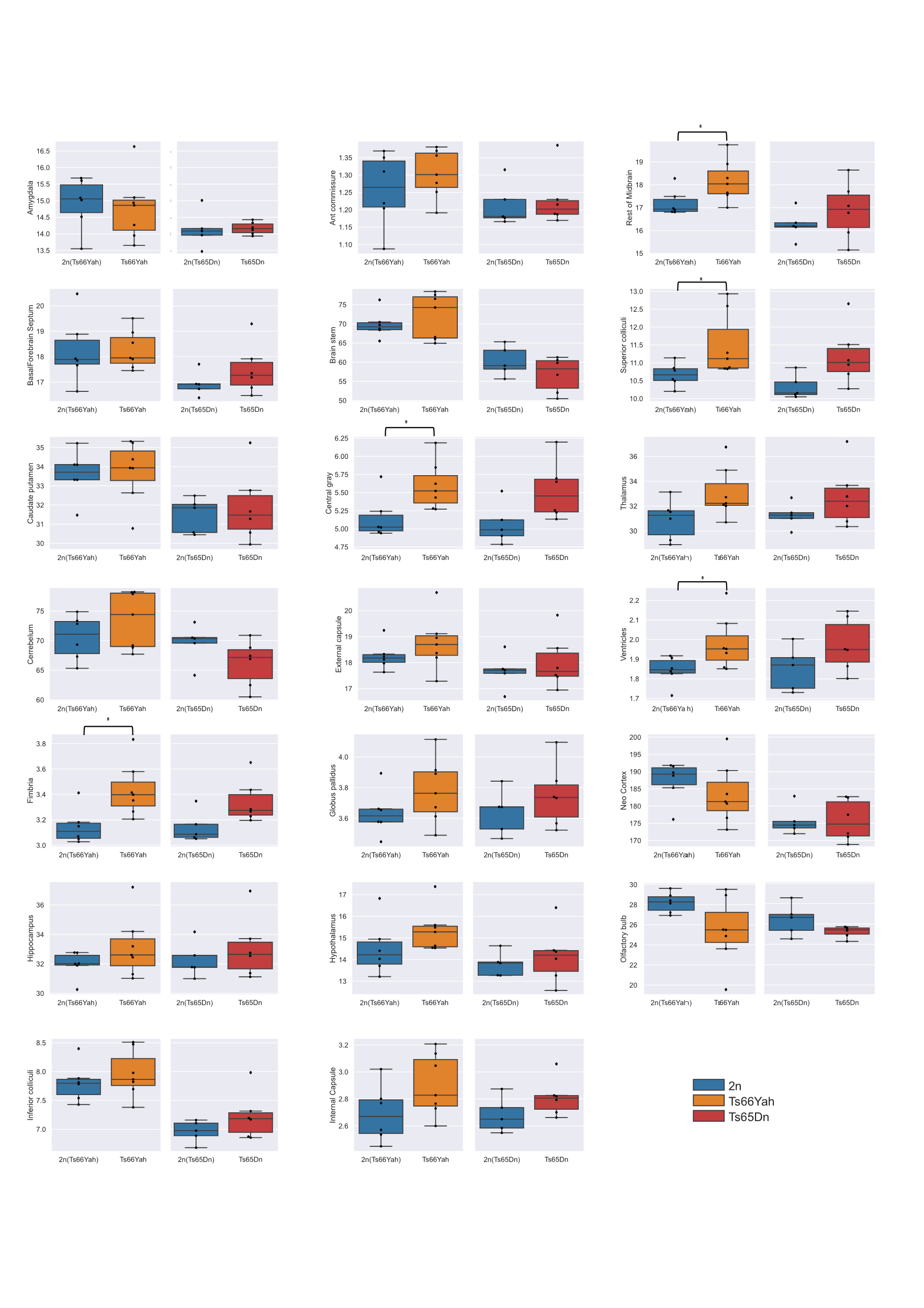

**Fig. S7. Changes observed by brain structures using MRI.** The changes in the morphology of specific brain structures and the direction of the changes (increase or decrease of volume) were the same in both DS lines. However, the amplitude of the changes was less severe in the Ts66Yah than in the Ts65Dn mice. Central grey, olfactory bulb or thalamus were not affected in the Ts66Yah mutants compared to the controls. Ts66Yah, n = 6 for 2n and n = 7 for Ts individuals and Ts65Dn n = 5 for control and n = 6 for Ts individuals. Box plots represent the median and quartiles. Statistical significance of differences between genotype was inferred by a two-tailed T test or Kruskal-Wallis non-parametric test. * p<0.05, **p<0.01, ***p<0.001.

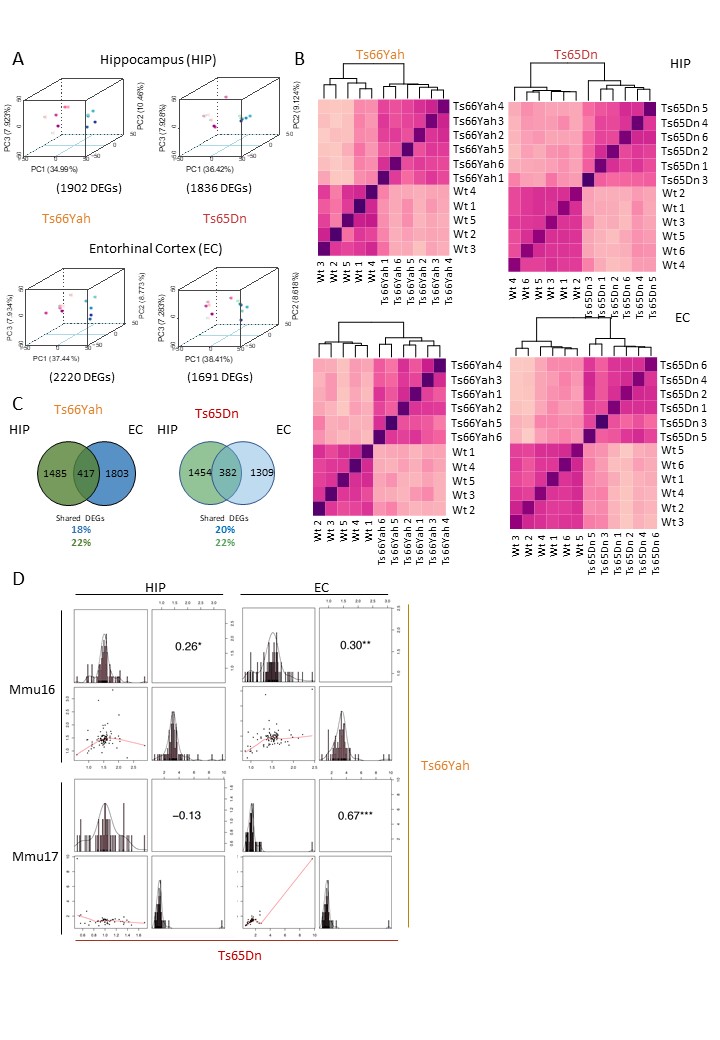

**Fig. S8 Gene expression analysis in the HIP and EC of Ts66Yah and Ts65Dn mice**. A. 3D-PCA on the DEGs for each adult hippocampal sample or entorhinal cortex sample allows to separate the animals carrying the Ts65Dn or Ts66Yah trisomies (pink) in comparison with the wild-type littermates (wt, blue). B. Homogenicity plot showing on the left how the Ts66Yah samples and the wt littermates cluster by Euclidean distance, on the right how the Ts65Dn samples and wt littermates cluster by Euclidian distance. C. Venn diagram showing the DEGs in common between Ts66Yah entorhinal cortex and hippocampi samples in the upper panel, and between Ts65Dn entorhinal cortex and hippocampi samples on the bottom panel. The shared DEGs correspond to 18% and 22% of the total DEGs identified respectively for the Ts66Yah entorhinal cortex and hippocampi samples; and 20% and 22 % for the Ts65Dn entorhinal cortex and hippocampi samples. In the intersection between the Venn diagrams circles are highlighted in the middle in bold the common genes regulated in mirroring sense, instead the common DEGs deregulated following the same regulatory sense specified in the upper (upregulated shared DEGs) and bottom (for the downregulated shared DEGs) part of the intersection between the Venn diagrams circles. D. Spearman correlation between Ts65Dn and Ts66Yah models based on TEGs identified in the Hippocampus (Hip) or the enthorinal cortices (EC) located in the Mmu16 homologous to Hsa21 or in the Mmu17 centromeric region.

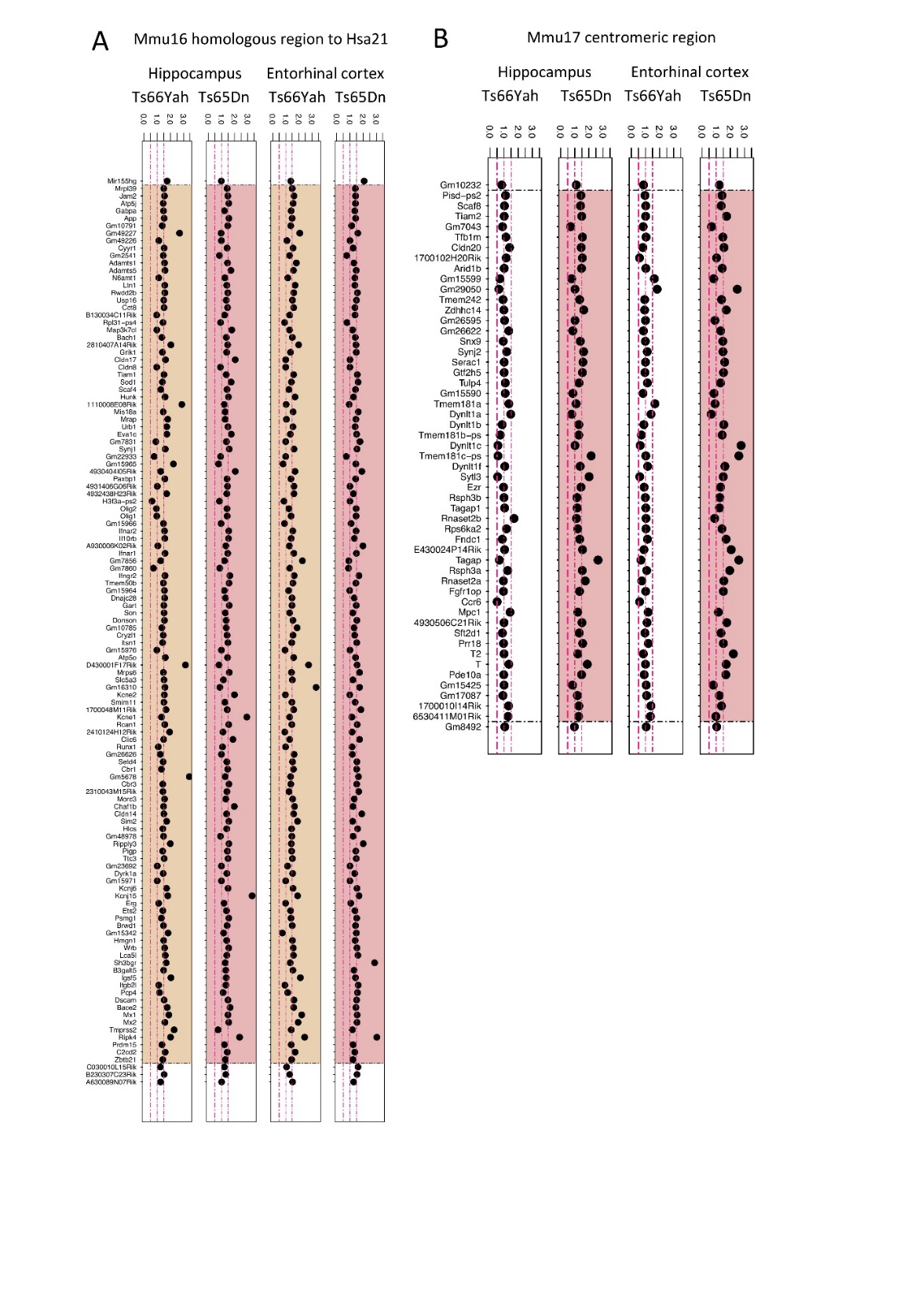

**Fig. S9. Expression of genes found on mouse chromosome 16 homologous region to Hsa21 (A) and on the 17 centromeric region (B) in Ts65Dn and Ts66Yah** **models**. **A**. The duplicated regions on Mmu16 found in the Ts65Dn and Ts66Yah models are highlighted in red or orange respectively whereas in **B**, the duplicated region on Mmu17 for the Ts65Dn model is highlighted in red and also for the Ts66Yah (no color). The genes are displayed following the order of their genomic start site coordinates from top to bottom.

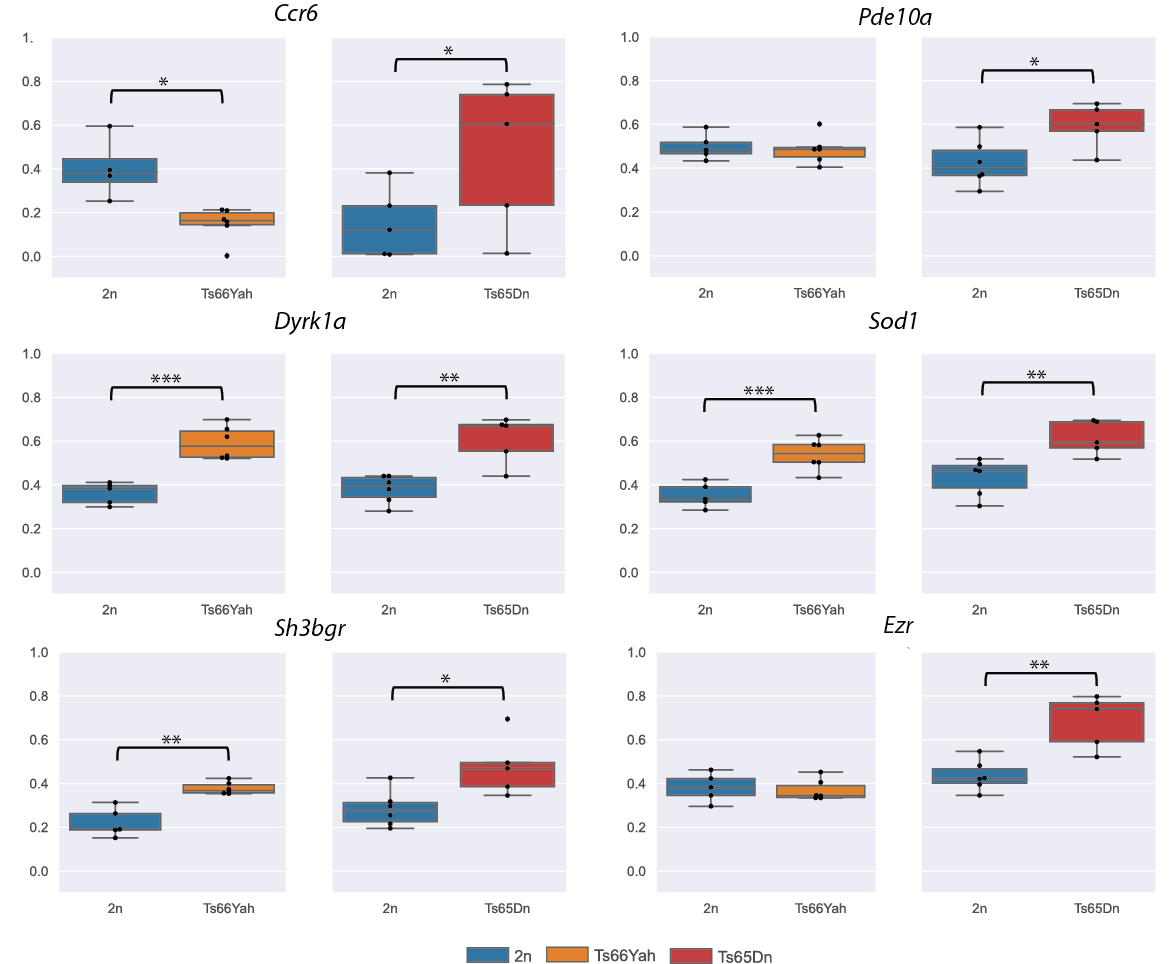

**Fig. S10. Quantification of the relative expression of selected messenger RNA isolated from the hippocampi of the Ts65Dn and Ts66Yah mouse models.**

Quantitative gene expression in trisomic mice compared to controls (2n: n=5; Ts66Yah n=6; 2n: n=6; Ts65Dn n=5) is reported here as boxplot. *Ccr6* was down-regulated in Ts66Yah and up-regulated in Ts65Dn. *Pde10a* was only upregulated in Ts65Dn, because is located in the Mmu17 deleted region. *Dyrk1a* and *Sod1* genes located in triplicated Mmu16 region were up regulated in both models. *Sh3bgr* and *Ezr* non-triplicated genes were dysregulated in different ways depending of the trisomic models (Box plots represent the median and quartiles, Statistical significance of differences between genotype was inferred by a two-tailed T test or Kruskal-Wallis non-parametric test. * p<0.05, **p<0.01, ***p<0.001).

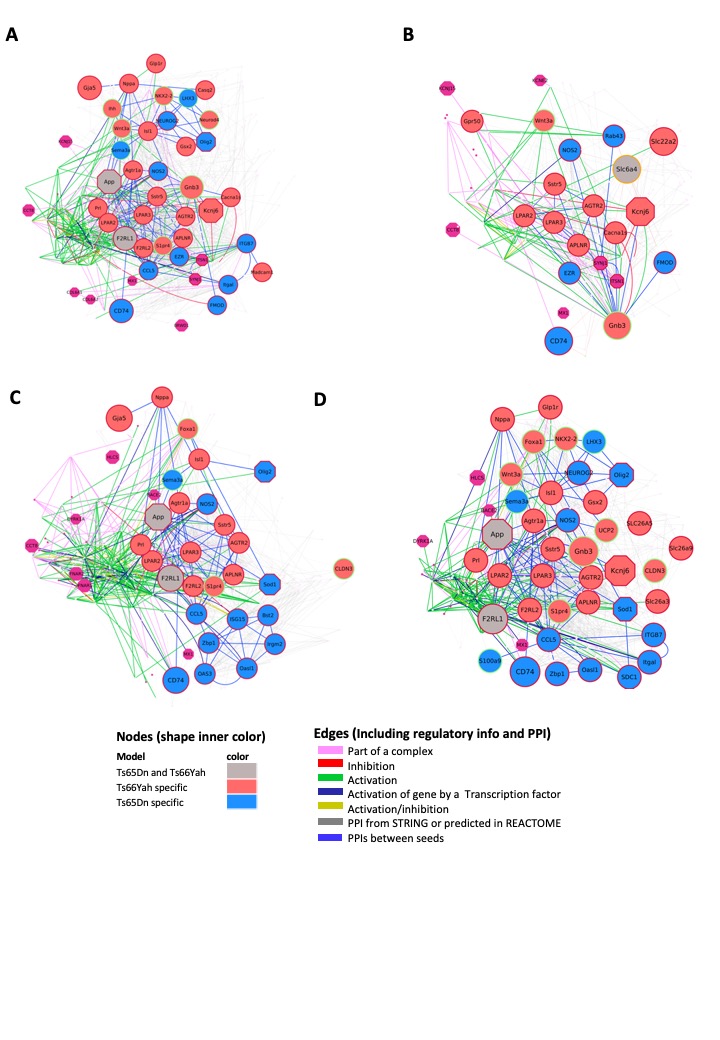

**Fig. S11. Ts65Dn and Ts66Yah Protein-protein interaction and regulatory gene connectivity network (RegPPINets) of genes involved in the synaptic meta-pathway identified in Ts65Dn or Ts66Yah highlighting the main biological cascades with altered connectivity**. These results in Ts66Yah and Ts65Dn support the role of the DS subnetworks centered on RHOA, SNARE proteins, NPY and DYRK1A cascades. A. Highlighting RHOA/RHOC/RHO 2^nd^ interactors. B. Highlighting SNARE VAMPS/SEC proteins 2^nd^ interactors. C. Highlighting DYRK1A 2^nd^ interactors. D. Highlighting NPY 2^nd^ interactors.

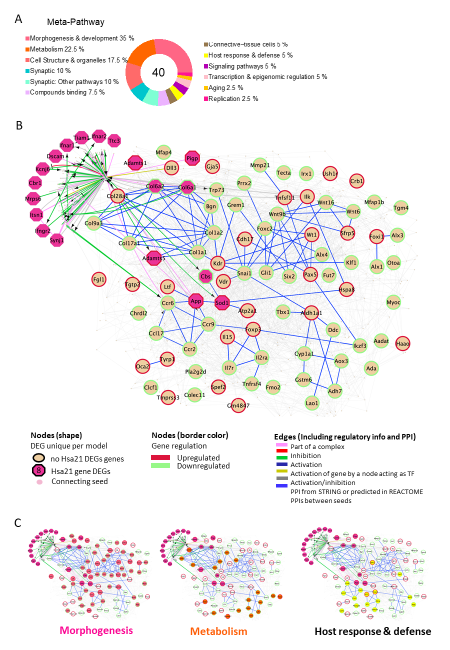

**Fig. S12. Ts66Yah entorhinal cortex functional Network analysis**. A. Donut plot representing the functional alterations grouped per meta-pathway in the Ts66Yah entorhinal cortex (noted as EC). In the centre of the donut plot the number of total altered pathways identified by gage; at left and right a summary showing the percentage of pathways included on each of the more populated meta-pathways. B. Full Ts66Yah entorhinal cortex RegPPINet network built using as seeds all the altered genes identified by gage in the Ts66Yah entorhinal cortex model visualized using the edge weighed spring embedded layout with the nodes ordered by betweenness index in Cytoscape. The full RegPPINet was built by querying STRING and selecting the PPIs with a medium confidence score (CS=0.4) coming from all sources of evidence. The shapes of the nodes and the inner colour represents the following information: Mustard circles: DEGs found altered in the model; Magenta octagons: represent HSA21 syntenic genes in mouse, some identified as contributing to those meta-pathway dysregulations by GAGE; Pallid pink Ellipses: In smaller size than the rest of the node shapes, represent connecting proteins added in the network to not have unconnected nodes assure the full connectivity of the network. The edge colour represents the type of interaction annotated by the PathPPI classification (Tang *et al.* 2015), and the ReactomeFIViz annotations as follows i) The GErel edges indicating expression were colored in blue and repression in yellow. ii) PPrel edges indicating activation were coloured in green, inhibition in red. Iii) Interactions between proteins known to be part of complexes in violet. Iv) Predicted interactions were represented in grey including the PPI interactions identified by STRING DB (Szklarczyk *et al.* 2017) after merging both networks. C. Ts66Yah entorhinal cortex RegPPINet network colored highlighting genes known to be involved in the following comorbidities, Host response & Immune system (yellow), genes involve in metabolism (orange), morphogenesis (magenta pink).

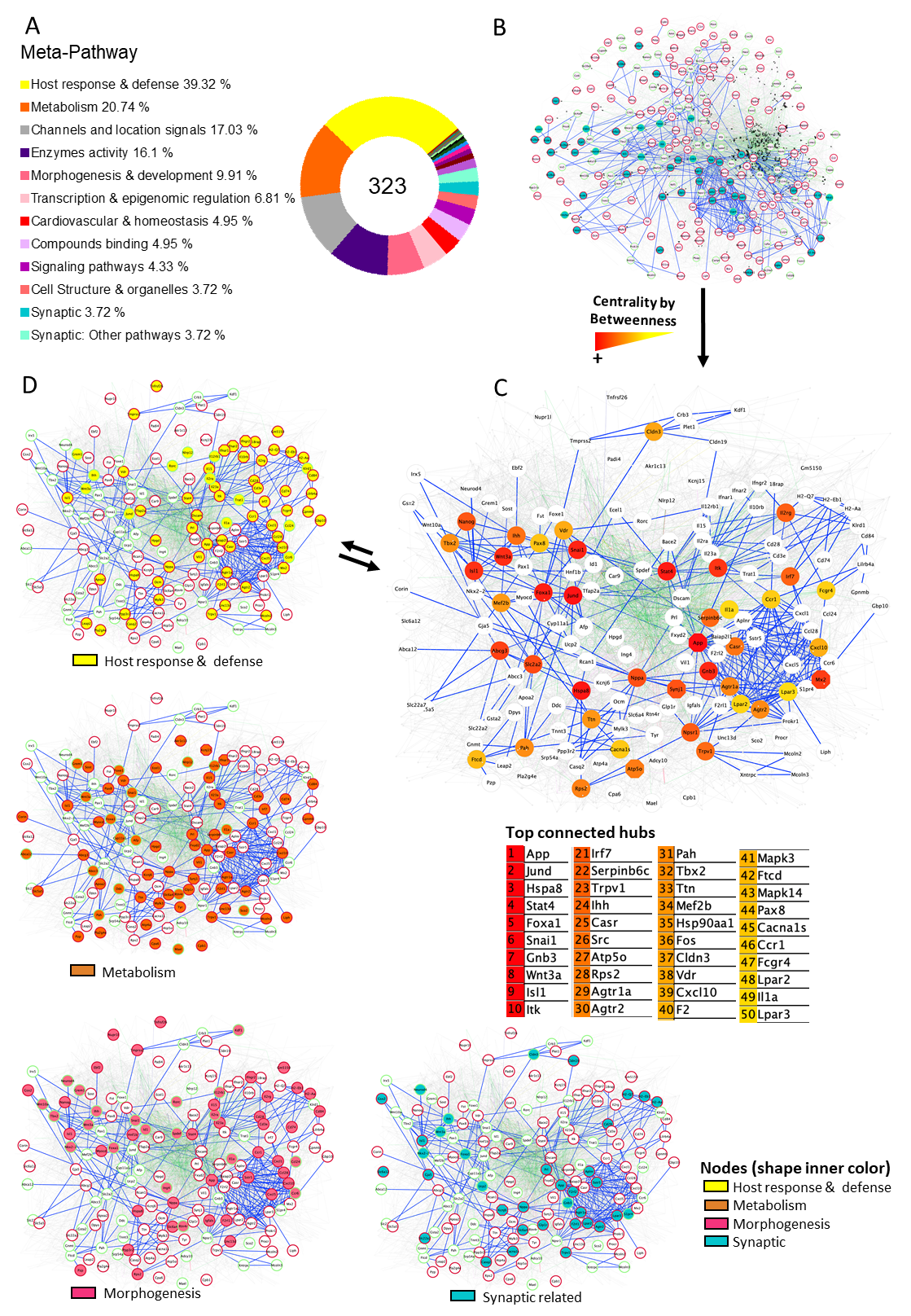

**Fig. S13. Ts66Yah Hippocampi functional Network analysis.** A, Donut plot representing the functional alteration per meta-pathway. In the centre the number of total altered pathways of the Ts66Yah Hippocampi and then the summary in percentages of the more populated meta-pathways. On the right, specifying the total number of pathways included on the synaptic meta-pathway**.** B, The full Ts66Yah Hippocampi RegPPINet network was built using as seeds all the genes identified by gage as altered in the Ts66Yah Hippocampi model visualized using the edge weighed spring embedded layout by betweenness index in Cytoscape. The full RegPPINet was built by querying STRING and selecting the PPIs with a medium confidence score (CS=0.4) coming from all sources of evidence. The shapes of the nodes represent the following information: Shapes: i) Pallid small pink ellipses: represent connecting proteins added to assure the full connectivity of the network; ii) Rest of the elliptic nodes represent genes identified by GAGE after q-val <0.1 cut off to be contributing even slightly, to any pathway of those found dysregulated inside the meta-pathway. Ii) Octagons represent the mouse Hsa21 syntenic genes. The edge colour represents the type of interaction annotated by following the PathPPI classification (Tang *et al.* 2015), and the ReactomeFIViz annotations as follows i) The GErel edges indicating expression were coloured in blue and repression in yellow. ii) PPrel edges indicating activation were coloured in green, inhibition in red. Iii) Interactions between proteins known to be part of complexes in violet. Iv) Predicted interactions were represented in grey including the PPI interactions identified by STRING DB (Szklarczyk *et al.* 2017) after merging both networks. * Represents those genes from the top 50 central genes list on the right side, known to be involved in synaptic pathways (they were linked to those functionalities in the GO/KEGG databases). C, Central Ts66Yah Hippocampi RegPPINet network colored i) in the upper panel highlighting the top 50 genes with the highest betweenness index value from red (higher index) to yellow (lower index) ii) in the lower panel highlighting the genes identified by gage to contribute to the alteration of synaptic related pathways. The list of the most central genes is included below the figure. D, Ts66Yah Hippocampi RegPPINet network colored highlighting genes known to be involved in comorbidities linked to morphogenesis, metabolism, and immune system highlighted in pink, orange and yellow on each network respectively.

#### References

Avants, B. B., C. L. Epstein, M. Grossman and J. C. Gee, 2008 Symmetric diffeomorphic image registration with cross-correlation: Evaluating automated labeling of elderly and neurodegenerative brain. Medical Image Analysis 12**:** 26-41.

Birling, M. C., L. Schaeffer, P. Andre, L. Lindner, D. Marechal *et al.*, 2017 Efficient and rapid generation of large genomic variants in rats and mice using CRISMERE. Sci Rep 7**:** 43331.

Duchon, A., M. Del Mar Muniz Moreno, S. Martin Lorenzo, M. P. Silva de Souza, C. Chevalier *et al.*, 2021 Multi-influential genetic interactions alter behaviour and cognition through six main biological cascades in Down syndrome mouse models. Hum Mol Genet 30**:** 771-788.

Duchon, A., M. Raveau, C. Chevalier, V. Nalesso, A. J. Sharp *et al.*, 2011 Identification of the translocation breakpoints in the Ts65Dn and Ts1Cje mouse lines: relevance for modeling down syndrome. Mammalian Genome 22**:** 674-684.

Ma, Y., P. R. Hof, S. C. Grant, S. J. Blackband, R. Bennett *et al.*, 2005 A three-dimensional digital atlas database of the adult C57BL/6J mouse brain by magnetic resonance microscopy. Neuroscience 135**:** 1203-1215.

Sawiak, S. J., N. I. Wood, G. B. Williams, A. J. Morton and T. A. Carpenter, 2009 SPMMouse: A new toolbox for SPM in animal brain. Proc. Int'l. Soc. Mag. Res. Med.**:** 1086.

Szklarczyk, D., J. H. Morris, H. Cook, M. Kuhn, S. Wyder *et al.*, 2017 The STRING database in 2017: quality-controlled protein-protein association networks, made broadly accessible. Nucleic Acids Res 45**:** D362-D368.

Tang, H., F. Zhong, W. Liu, F. He and H. Xie, 2015 PathPPI: an integrated dataset of human pathways and protein-protein interactions. Sci China Life Sci 58**:** 579-589.

Tustison, N. J., B. B. Avants, P. A. Cook, Y. J. Zheng, A. Egan *et al.*, 2010 N4ITK: Improved N3 Bias Correction. Ieee Transactions on Medical Imaging 29**:** 1310-1320.

Warfield, S. K., K. H. Zou and W. M. Wells, 2004 Simultaneous truth and performance level estimation (STAPLE): An algorithm for the validation of image segmentation. Ieee Transactions on Medical Imaging 23**:** 903-921.
